# A neural network for information seeking

**DOI:** 10.1101/720433

**Authors:** J. Kael White, Ethan S. Bromberg-Martin, Sarah R. Heilbronner, Kaining Zhang, Julia Pai, Suzanne N. Haber, Ilya E. Monosov

## Abstract

Humans and other animals often show a strong desire to know the uncertain rewards their future has in store, even when they cannot use this information to influence the outcome. However, it is unknown how the brain predicts opportunities to gain information and motivates this information seeking behavior. Here we show that neurons in a network of interconnected subregions of primate anterior cingulate cortex and basal ganglia predict the moment of gaining information about uncertain rewards. Spontaneous increases in their information prediction signals are followed by gaze shifts toward objects associated with resolving uncertainty, and pharmacologically disrupting this network reduces the motivation to seek information. These findings demonstrate a cortico-basal ganglia mechanism responsible for motivating actions to resolve uncertainty by seeking knowledge about the future.

## INTRODUCTION

Humans and other animals often express a strong desire to know the uncertain rewards their future has in store, even when there is no way to use this information to influence the outcome ^1, 2^. This information seeking behavior is not predicted by standard theories of reinforcement learning and reward-seeking, and has been the subject of investigation in the fields of psychology, economics, and neuroscience ^3–8^. These studies have revealed several neuronal populations suitable for regulating specific aspects of information seeking behavior. Certain prefrontal cortical neurons in monkeys and areas in humans are transiently activated by visual cues associated with future information gain about uncertain outcomes ^9, 10^. In addition, evidence from both monkeys and humans suggests that gaining information about uncertain outcomes can be a form of reward, since it activates the same ‘reward prediction error’ circuitry as primary rewards like food and water ^6, 10–12^.

These findings, however, leave a major question unanswered: how does the brain motivate online information seeking behavior? That is, how does the brain ‘bridge the gap’ to sustain our motivation to seek information, in between the time when we first learn of an opportunity to gain information and the time when we finally obtain it? At present, it is completely unknown what neural networks are causally responsible for information seeking behavior, and it is unknown what neural code they could use to regulate the strength of information seeking from moment to moment in response to environmental demands.

To gain insight into this question, we took inspiration from a field of study that has performed extensive investigations into the analogous question for conventional primary reward seeking behavior. These studies have revealed that the brain contains neural populations encoding *reward predictions*. Many such neurons have sustained activity that starts from the moment when reward can first be predicted, scales with the expected amount of reward, and ramps up to the expected time when the reward become available ^13^. In many cases, these reward prediction signals have been directly linked to online reward seeking behavior: the strength of the signals are correlated with reward seeking behaviors ^14–17^, and perturbing the signals alters reward seeking ^18–20^.

We therefore hypothesized that information seeking behavior is motivated by an analogous neural network that encodes *information predictions*. Specifically, in order for a neural network to motivate ongoing information seeking behavior, it must (A) monitor the level of uncertainty about future events, (B) anticipate the time when information will become available to resolve the uncertainty, (C) activate before information-seeking behaviors, such as gaze shifts to inspect the source of uncertainty, (D) causally motivate behavior to obtain information.

Here we demonstrate that these criteria are met by an anatomically interconnected network comprising three areas of the primate brain: the anterior cingulate cortex (ACC) and two subregions of the basal ganglia (BG), the internal-capsule-bordering portion of the dorsal striatum (icbDS) and the anterior pallidum including anterior globus pallidus and the ventral pallidum (Pal).

## RESULTS

### A cortico-basal ganglia network monitors reward uncertainty

To identify neural networks that are selectively responsive to reward uncertainty, we presented monkeys with fractal visual conditioned stimuli (CSs) predicting the delivery of a future juice reward with 0%, 25%, 50%, 75%, and 100% probabilities ^21, 22^. All three areas contained numerous neurons that were strongly activated or inhibited by all of the CSs that cued uncertain rewards (Fig 1B; Fig 1C, cyan, blue, turquoise; 25%, 50%, and 75% reward CSs). These responses were primarily excitatory in ACC and icbDS and often inhibitory in Pal (Fig 1B). The average responses consisted of sustained ramping to the moment when the uncertain outcome would occur (Fig 1C,D). Importantly, unlike conventional reward-related neurons in these areas ^21–23^, these neurons were more responsive to reward uncertainty than reward value; their responses were substantially lower for the CSs that cued certain outcomes, even though they had the highest and lowest values in the task (black, 100%, certain reward; gray, 0%, certain no-reward; Fig 1C-E). Furthermore, many of these neurons responded to uncertainty in a graded manner ^24^: they responded most in the condition with maximal uncertainty (50% reward), less in conditions with intermediate uncertainty (75% and 25% reward), and least in conditions with no uncertainty (100% and 0% reward) ^21, 22^. Specifically, neurons with a significantly greater average response to the 50% CS than to the 75 and 25% CSs were found in all three areas (Fig 1F, dark blue; p < 0.05, signed-rank test), with much greater prevalence than expected by chance (p < 0.001 in each area, binomial tests), and were significantly more prevalent than the opposite response pattern (Fig 1F, cyan; p < 0.05 in each area, binomial tests).

**Figure 1.**
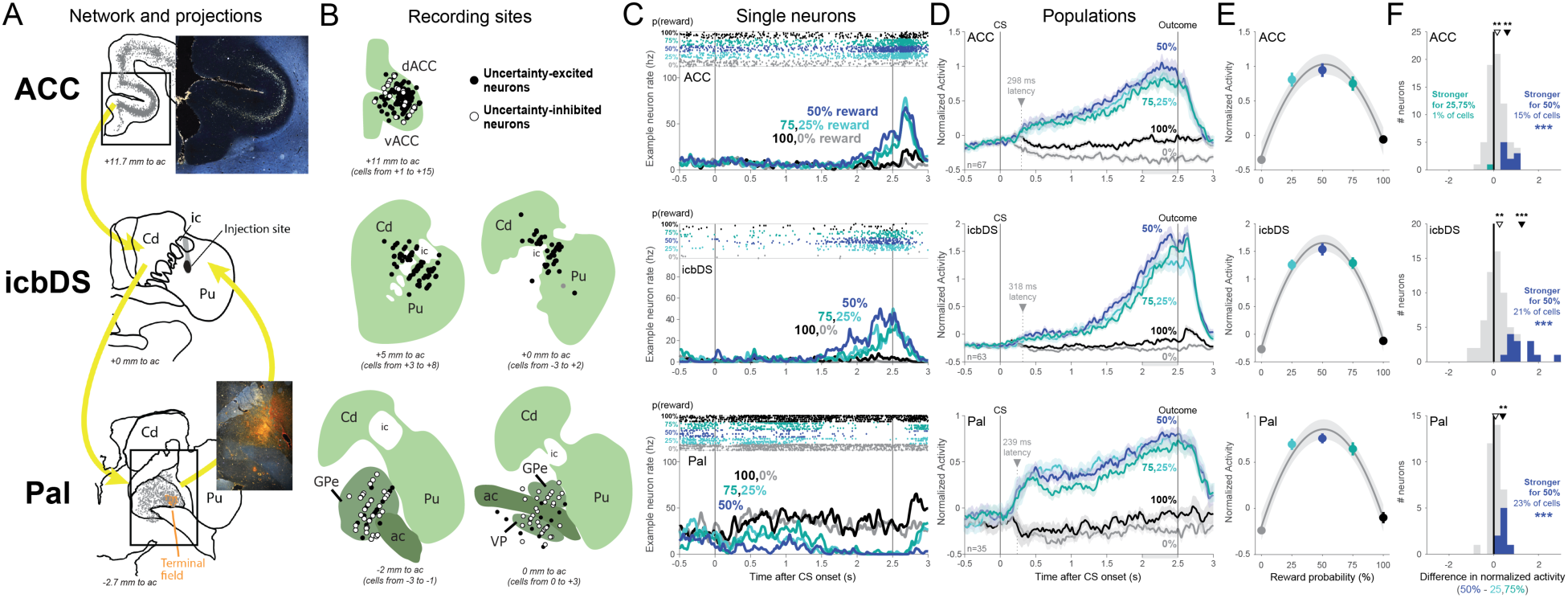
A cortico-basal ganglia network signals reward uncertainty. (**A**) ACC-icbDS-Pal network. Lucifer yellow was injected within icbDS, i.e. the internal-capsule bordering region of dorsal striatum (middle, black area). Retrogradely labeled cell bodies (gray) were found in ACC (top; inset) and Pal (bottom). Anterogradely-labeled fibers (orange area) were found in Pal (bottom; inset). Abbreviations: ic, internal capsule; Cd, caudate nucleus; Pu, putamen; dACC, dorsal ACC; Pal, pallidum; icbDS, vACC, ventral ACC; ac, anterior commissure; GPe, globus pallidus external segment; VP, ventral pallidum. Text indicates the plane’s anterior-posterior position relative to the midline anterior commissure. (**B**) Reconstruction of recording sites in ACC (top), icbDS (middle), and Pal (bottom). Circles indicate locations of neurons that responded to uncertainty with significant excitation (black) or inhibition (white). Structures are shown in the coronal plane. Neurons are projected onto the nearest shown plane; text indicates the range of neuron locations. (**C**) Example responses to the uncertainty task from neurons in ACC (top), icbDS (middle), and Pal (bottom). The ACC and icbDS neurons have excitatory ramping activity up to the time of uncertain reward: strongest for highly uncertain rewards (blue, 50%), strong for moderately uncertain rewards (turquoise colors, 25% and 75%), and weak or absent for certain outcomes (black, 100%; gray, 0%). The Pal neuron has similar ramping activity with an opposite, inhibitory direction of response. *Top of panel:* times of each spike (dots) on each trial (rows). *Bottom of panel:* smoothed firing rate for each CS. (**D**) The population average normalized activity of uncertainty coding neurons in each area ramps up to the time of the uncertain outcome. Shaded areas are ± 1 SE. Arrow, dashed line, and text indicate each area’s latency of uncertainty coding. Gray area below x-axis is the pre-outcome analysis time window. (**E**) Population average pre-outcome normalized activity is well fitted with a second-order polynomial function of reward probability (gray lines; shaded areas are ± 1 bootstrap SE) indicating an inverted-U relationship between reward probability and neural responses, as expected for uncertainty coding. (**F**) Graded coding of reward uncertainty. Histograms show each neuron’s difference in normalized activity between CSs with high (50%) vs. moderate (25,75%) reward uncertainty. Colored neurons have significantly differential activity. In all areas, more neurons are significantly more active for high uncertainty (blue) than moderate uncertainty (turquoise). *** indicates more neurons than expected by chance (p ≤ 0.001). Arrows indicate the mean of all neurons (open arrow) and all neurons with significant differential activity (filled arrows); *, **, *** indicate the significance of their difference from zero, p < 0.05, 0.01, 0.001 (signed-rank tests).

The network refined its uncertainty signals over time. Uncertainty signals emerged markedly earlier in Pal (Fig 1D, arrows; Fig S1; significantly shorter latency in Pal than ACC and icbDS, both p ≤ 0.005, permutation tests; Methods) but this initial signal did not yet encode the graded level of uncertainty (i.e. similar activity for 25, 50, and 75%; Fig S1). Uncertainty signals later emerged in both ACC and icbDS at roughly similar latencies (Fig 1D; no significant latency difference, p = 0.35), and those two areas first significantly encoded the graded level of uncertainty, doing so before Pal (50 > 25,75%, Fig 1D; Fig S1). These results indicate that a rapid but rough Pal uncertainty signal is followed by a slower, graded signal in cortico-striatal areas.

Given that these uncertainty-related regions have closely related neural signals, we tested whether they form an interconnected network. We injected the bidirectional tracer Lucifer yellow (LY) into the internal capsule-bordering regions of DS where uncertainty-responsive neurons were found (Fig 1A; Fig S2). This produced a large number of retrogradely labeled cells in the ACC (Fig 1A, gray) including the subregion containing uncertainty-responsive neurons (Fig 1B), indicative of a unidirectional ACC→icbDS projection. In addition, this produced both retrogradely labeled cells (Fig 1A, gray) and anterogradely labeled fibers in Pal (Fig 1A, orange area) including the region containing uncertainty-responsive neurons (Fig 1B), indicative of bidirectional icbDS→Pal and Pal→icbDS projections. The presence of bidirectional icbDS→Pal and Pal→icbDS projections were confirmed by tracer injection into Pal (Fig S2). These connections are consistent with established cortico-BG circuits ^25–27^ and are ideally suited to support the forms of uncertainty coding we observed in all three areas. Notably, these connections are consistent with uncertainty coding being primarily excitatory in ACC and icbDS and more commonly inhibitory in Pal (Fig 1B, S1) since cortex-striatum projections are excitatory and striatum-pallidum projections are mutually inhibitory ^28^. They are also consistent with uncertainty signals emerging first in Pal, as Pal can communicate with ACC and icbDS via direct Pal→icbDS projections (Fig 1A) and classic cortico-basal ganglia-thalamic-cortical loops ^25^.

### The network anticipates gaining information to resolve reward uncertainty

Our findings thus far identify an interconnected cortico-BG network that signals reward uncertainty with ramping anticipatory activity. This raises a key question: what event is the network anticipating? Most crucially, does the network anticipate the moment of receiving an uncertain outcome *per se*, or the moment of receiving *information* to resolve the uncertainty?

To answer this question, we designed a task to separate the time of receiving information from the time of receiving the outcome (information task, Fig 2A). On each trial the monkey was shown a fractal CS that indicated that a reward would be delivered in 3 seconds with 100%, 50%, or 0% probability. There were two types of CSs. *Information-predictive CSs* (Info CSs) were followed after 1 second by an informative visual cue whose color indicated the upcoming outcome (e.g. orange → reward, gray → no reward). *Non-information-predictive CSs* (Noinfo CSs) were followed by a non-informative cue whose color was randomized on each trial and hence did not indicate the upcoming outcome. Note that in this terminology the terms Info CS and Noinfo CS refer to whether the CS is followed by an informative cue (not to whether the CS itself conveys information about reward). Importantly, there was no way for animals to use the information to control or influence the outcome. Thus, neurons that anticipate the moment of receiving information to resolve uncertainty (Fig 2B, “Hypothesis 1”) should be activated at distinct times on Info and Noinfo CS trials. On Info CS trials they should activate in anticipation of receiving information from the informative cue (Fig 2B, left, red arrow). On Noinfo CS trials they should activate in anticipation of outcome delivery, because that is when the animal is first informed of the outcome (by receiving either juice or no juice; Fig 2B, left, blue arrow). On the other hand, neurons that simply anticipate uncertain outcomes (Fig 2B, “Hypothesis 2”) should respond identically during the Info and Noinfo CSs because both types of CSs are associated with identical future reward outcomes (the same reward probability, amount, and timing). They should only differentiate between Info and Noinfo trials in anticipation of the outcome, when outcomes are certain on Info trials but uncertain on Noinfo trials (Fig 2B, right, blue arrow).

**Figure 2.**
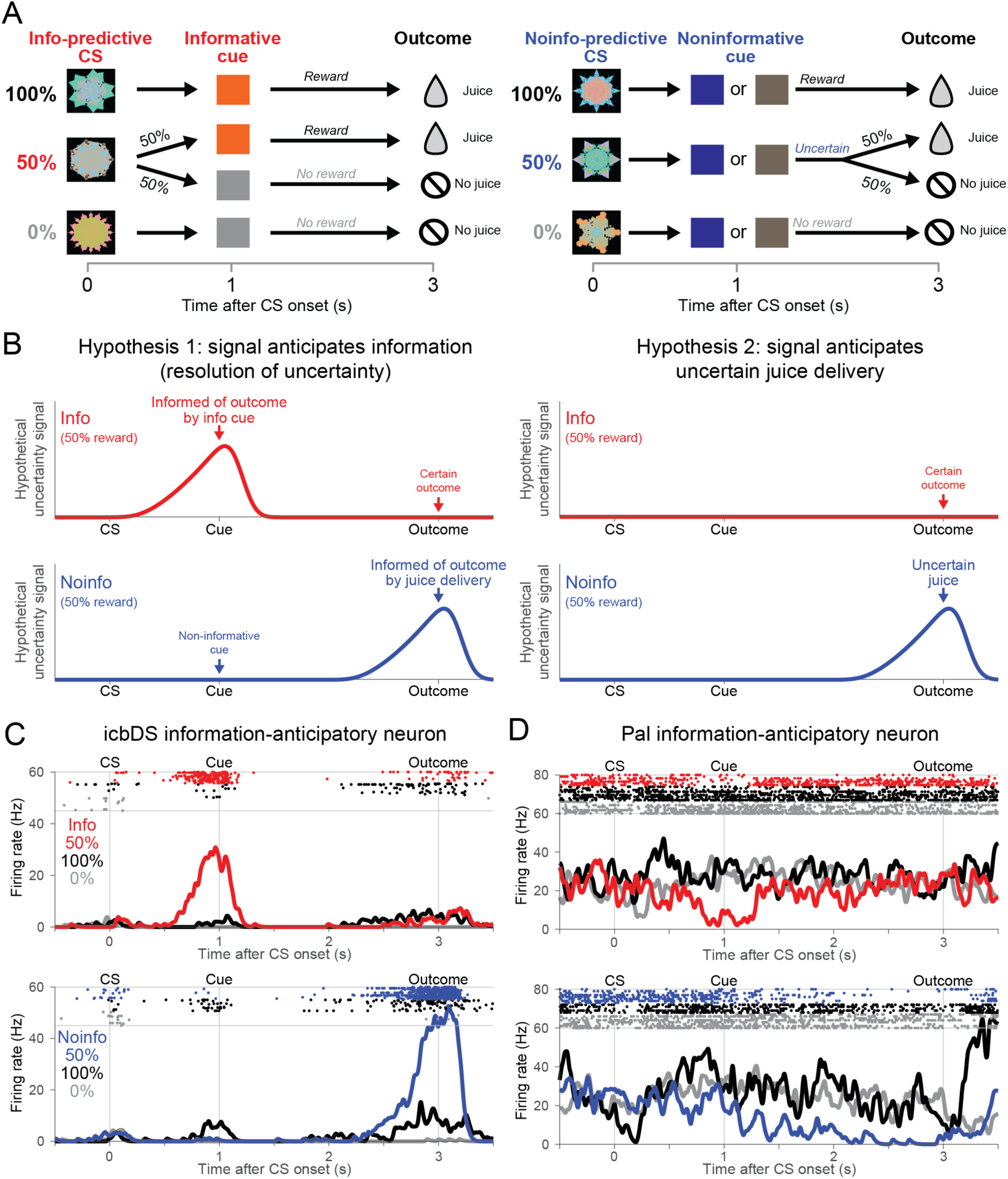
Neuronal activity anticipating the moment of gaining information to resolve reward uncertainty. (**A**) Information task. On Info CS trials, CSs predict 100, 50, or 0% reward, and are followed by an informative cue that indicates the outcome with certainty. On Noinfo CS trials, analogous CSs are followed by one of two non-informative cues that are randomized and hence leave the outcome uncertain until it is delivered at the end of the trial. (**B**) Testing two hypotheses about the origin of uncertainty-related activity. In Hypothesis 1 (left), activity on 50% reward trials anticipates the moment of gaining information about the uncertain outcome, and hence anticipates both informative cues (top, red, Info CS) and uncertain outcomes (bottom, blue, Noinfo CS). In Hypothesis 2 (right), activity simply anticipates uncertain outcome delivery, and hence has no differential activity during the CS period because it only anticipates uncertain outcomes (bottom, blue, Noinfo CS). (**C**) An icbDS neuron with information-anticipatory activity. This neuron has strong ramping activation on 50% reward trials that anticipates informative cues (top, red, Info CS) and uncertain outcomes (bottom, blue, Noinfo CS). Its activity is greatly reduced or absent when the outcome is certain (100% reward, black; 0% reward, gray). (**D**) An example Pal neuron with information-anticipatory activity. This neuron has a similar response pattern but with ramping inhibition rather than excitation.

Indeed, the cortico-BG network contained a substantial population of neurons that anticipated the moment of gaining information to resolve reward uncertainty. For example, the icbDS neuron in Fig 2C bore a close resemblance to the hypothetical information anticipatory signal (compare to Fig 2B). This neuron was strongly activated in advance of receiving information about uncertain rewards, both on Info CS trials during the CS period in anticipation of viewing an informative cue (top, red) and on Noinfo CS trials during the cue period in anticipation of reward delivery (bottom, blue). These activations were highly similar even though the information on these two trial types was conveyed through different modalities (visual cue vs. juice delivery). Importantly, this neuron had much lower activity during the same time periods on other trial types when information was not expected (e.g. during the Noinfo CS in advance of non-informative cues, and during the informative cues in advance of the already-known outcome). Similarly, the Pal neuron in Fig 2D had a comparable response pattern except that it anticipated information with inhibitions rather than excitations. Furthermore, as in the original uncertainty task, these responses cannot be explained as merely anticipating juice reward or as encoding the expected amount of juice associated with each CS; responses were strong when reward was uncertain (50% reward CS trials), but were weak or absent when reward was certain (100% reward CS trials).

We recorded from 154 uncertainty coding neurons using information tasks (ACC n=63, icbDS n=24, Pal n=67; Methods). We defined a neuron’s uncertainty signal as its ROC area for using its firing rate to distinguish trials with uncertain rewards vs. certain rewards (e.g. 50% vs. 100% and 0%). Crucially, we classified neurons as uncertainty coding solely based on whether they had a significant uncertainty signal during a 0.5 sec time window before outcome delivery on Noinfo trials (p < 0.05, rank-sum test). We then independently asked whether the same neurons also had uncertainty signals before cue onset and on Info trials. Specifically, we calculated an “Informative Cue Anticipation Index” defined as the difference between the uncertainty signal on Info and Noinfo CS trials. In many neurons, this index became different from zero shortly after CS onset and built up to the time of the cue, in either an excitatory or inhibitory manner (Fig 3A, red or blue). We defined a cell as information-anticipatory if the index was significantly different from zero during the 0.5 sec immediately before cue onset. Information-anticipatory neurons were highly prevalent in all three areas of the network (Fig 3B; 41%, 54%, and 39% of uncertainty-related neurons in ACC, icbDS, and Pal (n=26/63, 13/24, and 26/67, respectively); more than expected by chance, all p < 0.001, binomial tests).

**Figure 3.**
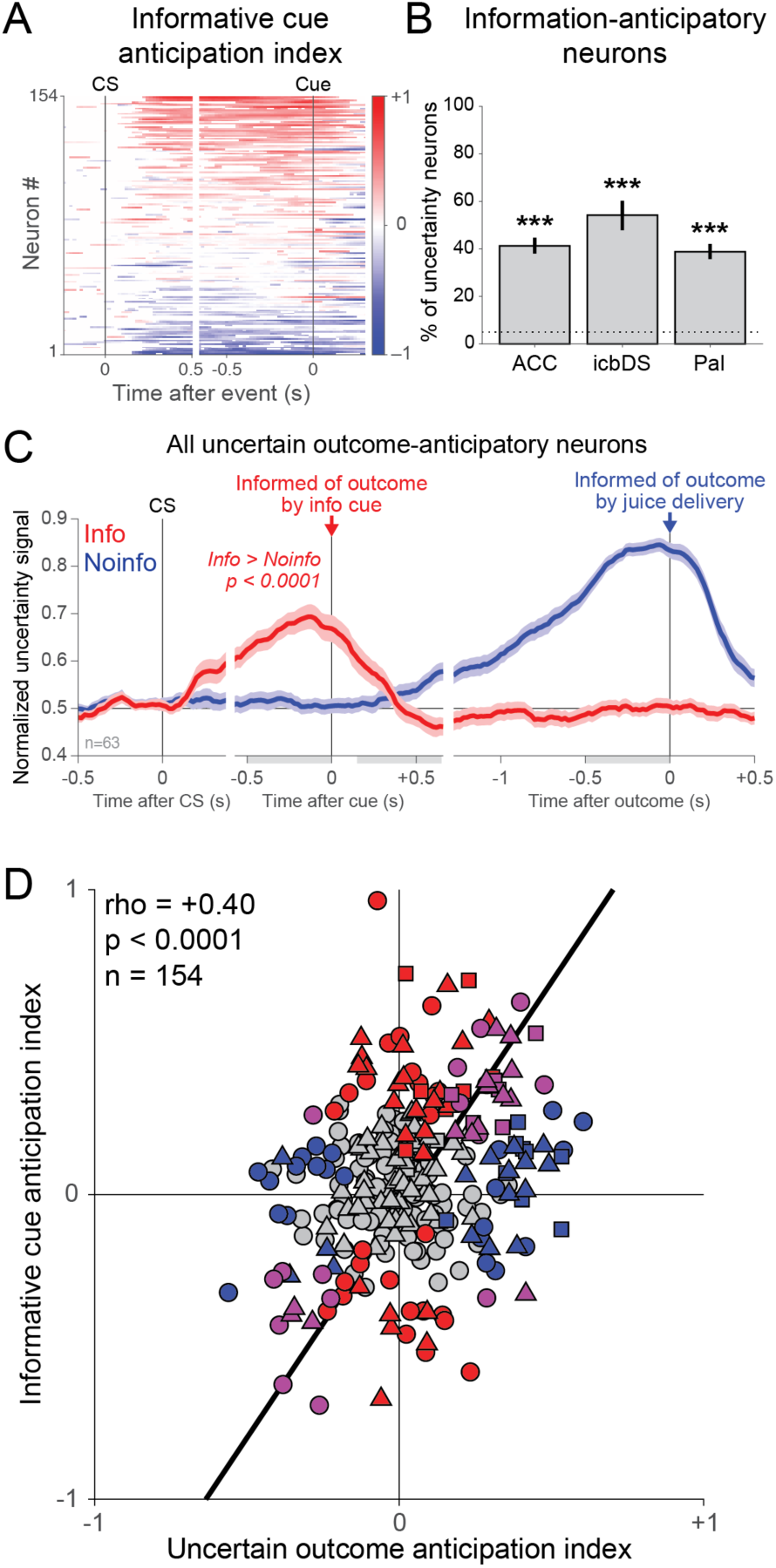
Prevalence of information-anticipatory activity in the cortico-basal ganglia network. (**A**) Uncertainty-related activity emerges preferentially during the Info CS in anticipation of informative cues. Each row is a neuron, and the color at each time point indicates the Informative Cue Anticipation Index (red: more positive uncertainty signal for Info, top; blue: more negative uncertainty signal for Info, bottom; color bar indicates scale). (**B**) Information-anticipatory activity was significantly present in approximately half of uncertainty-related neurons in each area. *** indicates significantly more neurons than expected by chance, p < 0.001. (**C**) Information-anticipatory activity was prevalent in neurons anticipating uncertain outcomes. Shown are the population average uncertainty signals on Info CS (red) and Noinfo CS (blue) trials from the subset of neurons with a significant Uncertain Outcome Anticipation Index (blue, index measured only using Noinfo CS trials). The population has clear activity on Info CS trials in anticipation of viewing information cues (red, text indicates p-value, rank-sum test). Error bars are ± 1 SE. Gray shaded areas are the time windows for calculating the indexes of cue anticipation (pre-cue window) and outcome anticipation (post-cue, pre-outcome windows). (**D**) Correlated anticipation of the two reward-informative task events. Many neurons have significant Informative Cue Anticipation Indexes (red, y-axis, p < 0.05, permutation test), Uncertain Outcome Anticipation Indexes (blue, x-axis), or both (purple). The two indexes are highly correlated; text indicates rank correlation and its p-value. ACC, icbDS, and Pal neurons are indicated by triangles, squares, and circles. Black line is a linear fit with type 2 regression.

Importantly, as expected for an information-anticipatory signal, the same neurons that anticipated informative cues on Info trials also commonly anticipated the time of uncertain outcomes on Noinfo trials, and did so in similar manners. To quantify this, we defined an analogous “Uncertain Outcome Anticipation Index” as the change in a neuron’s uncertainty signal from the beginning to the end of the cue period on Noinfo trials. This index was significant in substantial number of single neurons in all three areas (37%, 63%, and 34% of neurons in ACC, icbDS, and Pal; more neurons than expected by chance in all areas, binomial tests, all p < 0.001) and tended to be most prevalent in icbDS (higher fraction of significant neurons than ACC or Pal, p = 0.0586 vs. ACC, p = 0.0313 vs. Pal, permutation tests). There was a strong correlation between the two neural anticipation indexes (rank correlation = 0.40, p < 0.001, Fig 3D). That is, many neurons, especially in ACC and icbDS, were activated in anticipation of receiving information from both cues and outcomes, while other neurons especially in Pal were inhibited for both. Thus, when examining neurons whose uncertainty signals on Noinfo trials significantly anticipated the time of the outcome, the average timecourse of their uncertainty signals on Info trials bore a strong resemblance to a hypothetical information-anticipatory signal (Fig 3C, compare to Fig 2B). Notably, an explicit test of competing hypotheses indicated that this activity was significantly different from hypothetical encoding of simple anticipation of uncertain outcome delivery or of the current level of uncertainty (Fig S3). Similar results were found in all three areas (Fig S4).

### Monkeys anticipate gaining information by gazing at objects associated with the uncertain outcome

Given the cortico-BG network’s strong information-predictive signal, we next asked whether information predictions evoke information seeking behavior in monkeys. Since monkeys, like humans, scan uncertain environments for information with their eyes ^6, 7, 22^, we hypothesized that monkeys may anticipate information by directing their gaze to objects in their environment associated with the uncertainty to be resolved. Consistent with previous work, we found that monkeys licked in anticipation of juice rewards (Fig S4) and that their gaze was attracted to visual objects based on their expected reward value (Fig 4A,B, 100% CS > 0% CS (black > gray), informative reward cue > no-reward cue (dark red > light red)).

**Figure 4.**
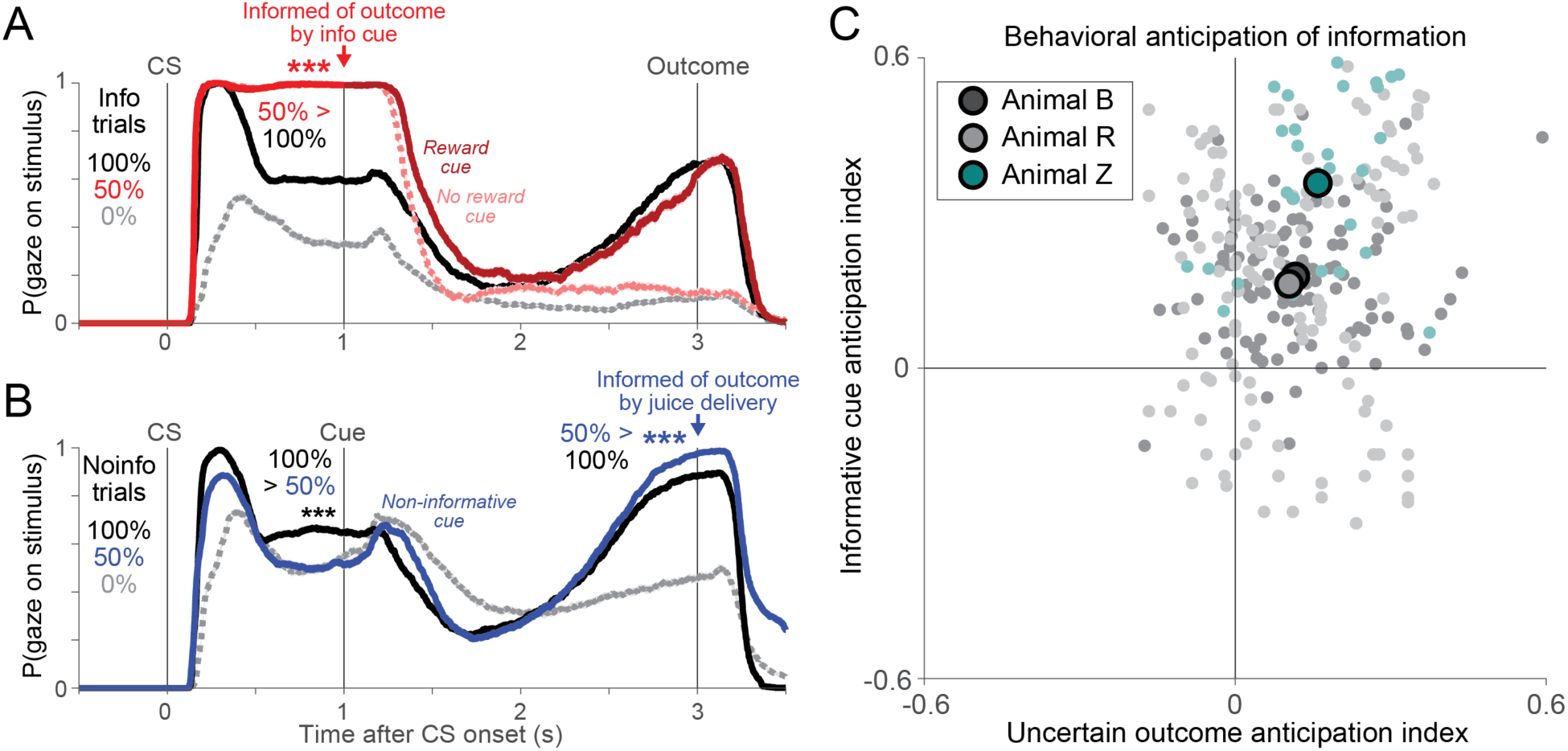
Monkeys anticipate the moment of gaining information by gazing at objects associated with the uncertain outcome. (**A**) Monkeys’ gaze on Info trials is attracted to CSs in anticipation of receiving informative cues about uncertain rewards. Lines indicate the probability at each millisecond that the animal gazes at the stimulus. Shaded areas indicate ± 1 SE (most too small to see). Monkeys gazed more at the 100% CS (black) than the 0% CS (gray), but gazed most of all at the uncertain 50% CS (red) which could be followed by informative cues indicating either reward (dark red) or no reward (pink). Gaze at reward cues then ramped up to the time of reward delivery, while gaze at no-reward cues was minimal. *** indicates p < 0.001 (signed-rank test) of the difference indicated by the text at the moment of a task event. (**B**) Gaze on Noinfo trials is attracted to objects in anticipation of outcome delivery. Same format as (A). During the CS period monkeys had roughly value-based gaze behavior (100% > 50% > 0%), but during the cue period their gaze on 50% reward trials ramped up in anticipation of the uncertain outcome until it became greater than all other conditions (50% > 100% > 0%). (**C**) Uncertainty-related gaze behavior in each animal significantly anticipated informative cues (y-axis) and uncertain outcomes (x-axis). Same format as Fig 3D, but analyzing gaze instead of neural activity. Colors indicate animals; light dots are single sessions; dark circles are means in each animal, error bars are ± 1 SE (most too small to see).

Strikingly, however, monkeys’ gaze was even more strongly attracted to objects based on their uncertainty, especially in the moments before receiving information to resolve that uncertainty. On Info trials, monkeys could anticipate receiving information during the Info 50% reward CS as they awaited the upcoming informative cue (Fig 4A, red arrow). Monkeys were substantially more likely to gaze at the Info 50% reward CS than any other CS at the moment the cue was about to appear (signed-rank tests, all p < 0.001). Importantly, this attraction of gaze was specifically related to upcoming information rather than to reward value or uncertainty per se. Monkeys gazed at the Info 50% reward CS far more than the Noinfo 50% reward CS, which was associated with exactly the same reward value and uncertainty but was not followed by information (Fig 3B, blue; signed-rank test, p < 0.001), and far more than the 100% reward CSs, which were associated with *double* the reward value but had no uncertainty to resolve (Fig 4A,B, black; signed-rank test, p < 0.001). Furthermore, this avid gaze at the Info 50% reward CS occurred despite near-zero licking, indicating that animals were anticipating the delivery of information, not juice reward (Fig S4).

Similarly, on Noinfo trials, monkeys could anticipate receiving information at the end of the cue period as they awaited the upcoming reward delivery or omission (Fig 4B, blue arrow). Again, at that time the monkeys gazed at the cue more during Noinfo 50% reward trials than all other conditions (signed-rank tests, all p < 0.001), even 100% reward trials that had double the reward value (Fig 4A,B, black). They did so even though the upcoming information about the outcome was delivered through a non-visual modality (receipt of juice or no juice), and even though 100%, 50%, and 0% Noinfo trials all used exactly the same set of visual cue stimuli; they gazed at those cues most avidly on Noinfo 50% reward trials when they were associated with an uncertain reward, and did so specifically in the moments before the uncertainty was going to be resolved. These monkeys also behaved consistently in standard uncertainty-related tasks that lacked informative cues (essentially treating all trials as Noinfo because no cues were available; Fig S4; Methods). Thus, when we analyzed monkeys’ gaze behavior in the same way that we analyzed neural spiking activity, we found that all monkeys had significantly positive Info Cue Anticipation and Uncertain Outcome Anticipation indexes, indicative of information-anticipatory behavior (Fig 4C, all p < 0.001, signed-rank test; Fig S5).

### Spontaneous fluctuations in network activity predict future information-anticipatory gaze behavior

Further investigation revealed that the monkeys’ information-anticipatory gaze behavior was linked to moment-to-moment variability in the cortico-BG network’s information anticipatory signals. Examining time windows immediately before receipt of information, we found that neural information signals were present even at moments when the monkey’s gaze was away from the visual stimulus, but were significantly stronger during matched time points from other trials when the monkey’s gaze was on that visual stimulus (Fig 5A). This occurred consistently in both the information task and standard probabilistic reward tasks (Fig 5A; Methods).

**Figure 5.**
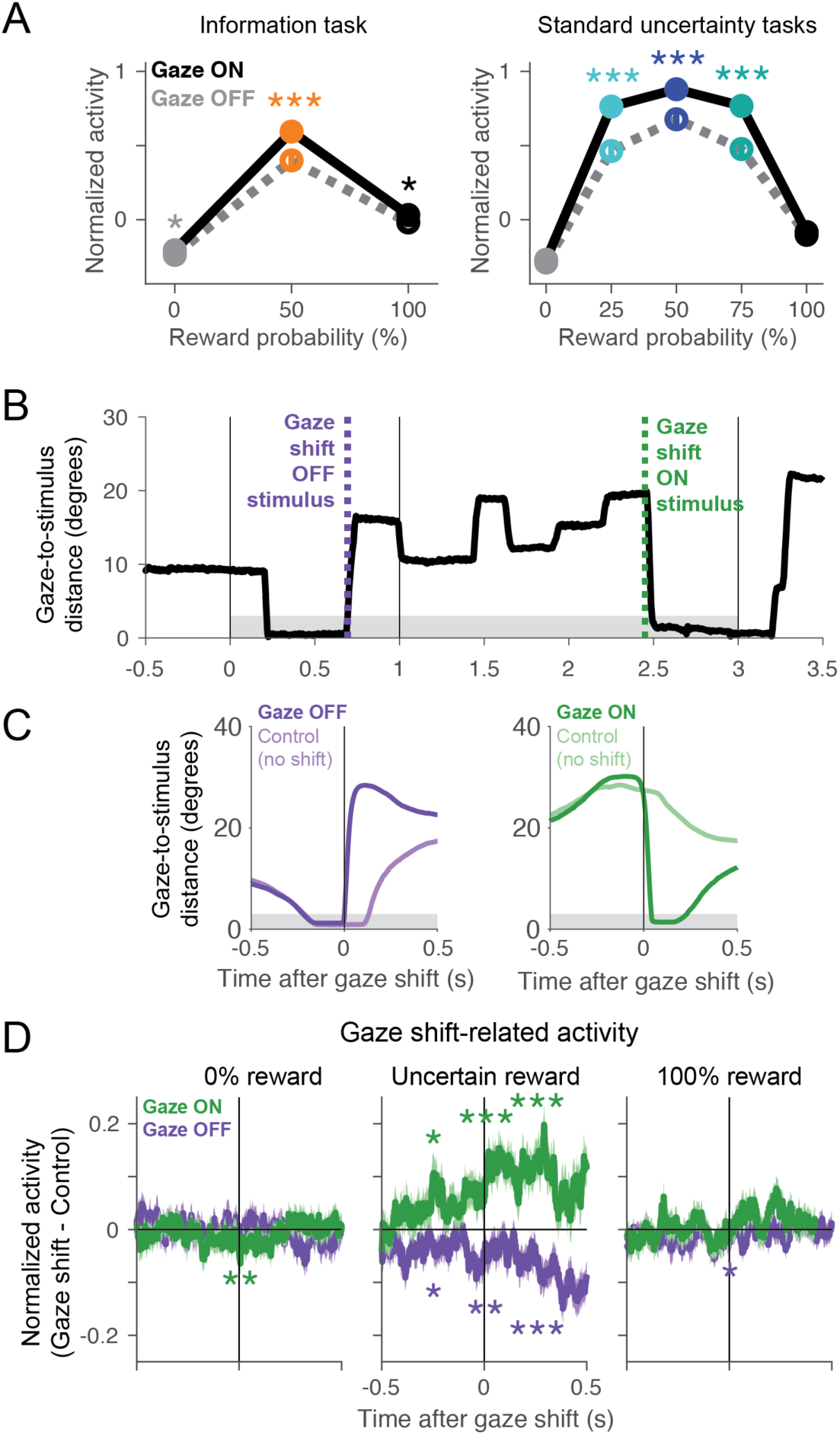
Fluctuations in cortico-BG network signals predict immediate gaze shifts toward or away from objects associated with resolving uncertainty. (**A**) Mean normalized activity before receipt of information is enhanced when gaze is on the stimulus (dark dots/lines) compared to when gaze is off the stimulus (light dots/lines), especially for intermediate reward probabilities when reward is uncertain. Data is shown for information tasks (left) and standard uncertainty tasks (right). This analysis was performed on all neurons where there was trial-to-trial variance in gaze behavior before receipt of information for all of the CSs (information task n=117: ACC n=62, icbDS n=24, Pal = 31; standard uncertainty tasks n=130: ACC n=53, icbDS n=53, Pal = 24). Error bars are ± 1 SE; asterisks indicate significance. (**B**) Timecourse of gaze distance to the center of the stimulus (black) on an example trial with the Noinfo 50% reward CS. Gray shaded area indicates the window for gaze to be on the stimulus. Green and purple dashed lines indicate gaze shifts off of and onto the stimulus. (**C**) Mean timecourse of gaze distance to the stimulus aligned on gaze shifts off and on the stimulus (purple, green) and aligned on control non-shift events (light purple, light green), excluding blinks. Both gaze shift and control events have similar gaze trajectories until the moment of the gaze shift. (**D**) Mean timecourse of gaze shift-related normalized activity aligned on gaze shifts onto the stimulus (green) and off the stimulus (purple). This analysis was performed on all neurons where there was at least one gaze shift onto the stimulus and one gaze shift off of the stimulus, as well as their appropriately matched control non-shift events, in each of three conditions: 0% reward (left), uncertain reward (middle), and 100% reward (right) (n=300: ACC n=156, icbDS n=78, Pal n=66). Shaded areas are ± 1 SE; asterisks indicate significance in time windows before, during, and after the gaze shift. Uncertainty-related activity was significantly enhanced before gaze shifts on the stimulus and reduced before gaze shifts off the stimulus, but only when animals were anticipating information about an uncertain reward (middle). Gaze-related activity was weak or absent when the outcome was certain, regardless of whether that outcome was no reward (left) or reward (right).

Importantly, neural activity was not simply enhanced in a generalized manner whenever gaze was on any stimulus, nor was the enhancement the result of encoding simple visuomotor variables such as gaze position or saccade direction (Fig S6). Instead, the enhancement took the form of a gain-like increase in the strength of neural information signals: activity was primarily enhanced when gazing at stimuli associated with uncertainty and its resolution, and was less affected when gazing at stimuli associated with certain outcomes (Fig 5A; significantly greater effect of gaze on normalized activity on uncertain trials than on certain trials, information task: p = 0.0009, standard uncertainty tasks: p < 0.0001, signed-rank tests). Thus, while information-anticipatory signals were present even after controlling for gaze state (Fig S7), they were particularly enhanced during gaze at visual stimuli associated with obtaining information.

This data suggested that the cortico-BG network’s information signals could be well suited to motivate the animal’s information-oriented behavior – a hypothesis we next tested through investigating the link between neurons and behavior and through direct pharmacological disruption.

First, we asked whether neural information signals strengthened *before* gaze shifts, consistent with a causal role in motivating information seeking, or *after* gaze shifts, potentially reflecting a sensory response to the stimulus being brought closer to the fovea (as has been reported in some neurons in the orbitofrontal cortex ^29^). To test this, we pooled data from all uncertainty-related neurons recorded in all three areas of the network in all tasks (n=222, 127, 129, in ACC, icbDS, and Pal; Fig S5,6; Table S1). We aligned neural activity at the time of each gaze shift onto or off of the visual stimulus (Fig 5B) and compared it to activity from matched control trials where these was no gaze shift at that time but which were matched in all other respects (i.e. the same neuron, CS, and cue, and similar initial gaze distance to the CS). The gaze shift trials and the matched control trials had similar gaze trajectories up until the moment of the gaze shift (Fig 5C). Strikingly, however, neural activity in the cortico-BG network was significantly altered long before the gaze shift, with significant enhancement of activity before gaze shifts onto the stimulus (Fig 5D, green, p < 0.05) and significant suppression of activity before gaze shifts off of the stimulus (Fig 5D, purple, p < 0.05). These gaze-shift related activity changes predominantly occurred when animals were anticipating information about an uncertain reward outcome; they were greatly reduced or absent when reward was certain to be delivered or omitted (Fig 5D). Also, these changes in activity grew stronger in successive time windows over the course of the gaze shift: they started before the gaze shift, grew stronger during the gaze shift but before the neurons received visual feedback from the change in eye position (given their visual response latencies > 0.1 sec), and became strongest after the gaze shift (Fig 5D; greater difference in activity between gaze shift on vs off in each time window compared to the previous window, p < 0.05, signed-rank test). Similar patterns were present for all three regions and for all CS locations and gaze shift directions (Fig S6).

To quantify each neuron’s modulation related to gaze and its timecourse relative to gaze shifts, we fit each neuron’s activity with a model of its responses to all of the CS and cue conditions (Methods). One set of parameters modeled the effects of gaze-related modulation (Fig 6A, top): a multiplicative change in response gain (e.g. enhanced information-related signals, similar to Fig 5A), an additive increase in firing rate (e.g. sensory or motor-related effects that have no effect on information signals), or a combination of both. A second set of parameters modeled the timecourse over which these modulations occurred relative to gaze shifts (Fig 6A, bottom): neurons could modulate their activity either before or after gaze shifts, and their modulation could come online either rapidly or gradually around the time of the gaze shift (similar to Fig 5D). The model accurately recovered the true effects and timecourses of gaze-modulation in simulated datasets (Fig S8).

**Figure 6.**
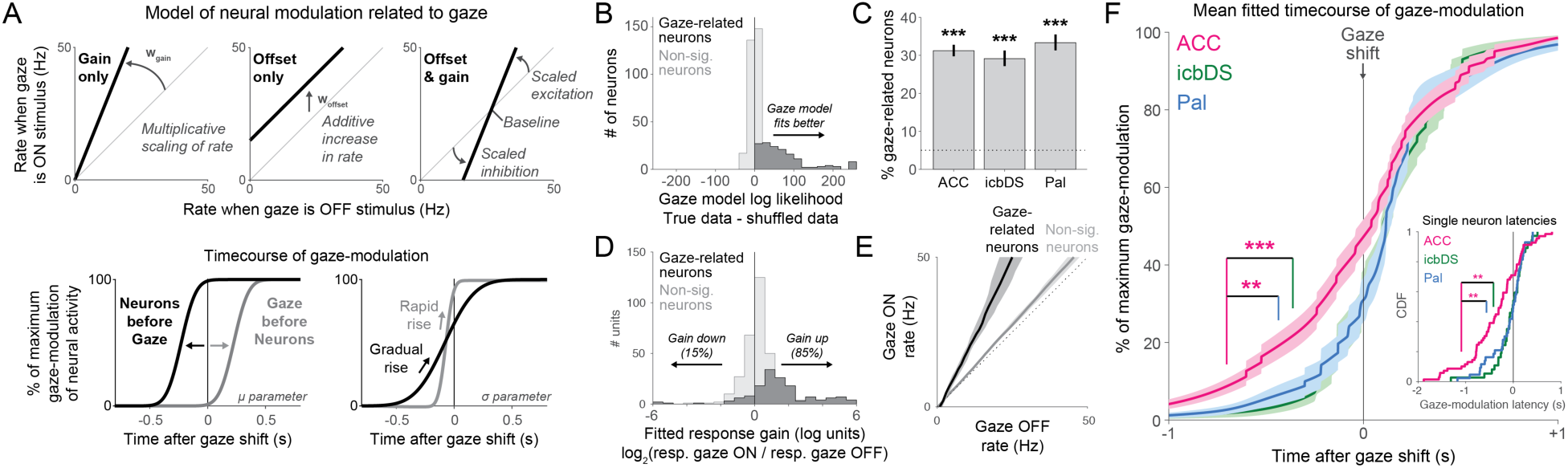
Prevalence and dynamics of gaze shift-related activity in the cortico-basal ganglia network. (**A**) Model of neural modulation related to gaze in single neurons. Top: model parameters fitting the effect of gaze on neural activity. Two parameters permit activity to be modulated by multiplicative scaling (left), additive increases or decreases in firing rate (middle), or a combination of both (right). Bottom: model parameters fitting the timecourse of gaze-modulation. Two parameters control the temporal offset between neural modulation and gaze shifts (left, neurons first vs. gaze first) and the rate at which neural modulation builds up over time (right, gradual vs. rapid). (**B**) Model fits indicated that 31% of neurons were significantly modulated in relation to gaze, indicated by significantly better log likelihood when fitting the true data compared to shuffled data (dark histogram). The remaining neurons had similar log likelihoods for true and shuffled data (light histogram). (**C**) Approximately 30% of neurons in each area were significantly gaze-related; all areas had more gaze-related neurons that expected by chance. (**D**) Fitted gaze-related response gains in each neuron, in log_2_ units (0 = no gain change; −1 = response is halved; +1 = response is doubled). Rare neurons fitted with complete reduction of responses during gaze (n=2 outliers) are plotted at −6. More neurons had gain increases than gain decreases, especially in gaze-related neurons (dark histogram). (**E**) Fitted relationship between gaze state and firing rate in gaze-related neurons (black) and non-significant neurons (gray) resembles an increase in response gain rather than a simple additive change in firing rate. Lines are medians, shaded areas are bootstrap 95% confidence intervals. (**F**) Mean fitted timecourse of gaze-modulation around the time of gaze shifts for neurons in each area with significant gaze-modulation (ACC, magenta; icbDS, green; Pal, blue). Asterisks indicate that the latency of gaze-modulation, defined at the time of 10% of maximal modulation, is significantly earlier in ACC than icbDS and Pal (p = 0.001 and p = 0.003, permutation tests including all gaze-modulated neurons; or p = 0.001 and p = 0.003, permutation tests including all neurons). Inset: cumulative distribution of gaze-modulation latencies for all individual neurons with significant gaze-modulation. The median latency is earlier in ACC than icbDS and Pal (p < 0.001 and p = 0.035, rank-sum tests including all neurons; or p = 0.002 and p = 0.006, rank-sum tests including all gaze-modulated neurons).

The model fits indicated that 31% of neurons had significant gaze-related modulation (permutation test, p < 0.05, Methods; 31%, 29%, 33% in ACC, icbDS, and Pal, Fig 3J; all areas above chance, binomial tests, p < 0.05, Fig 6B,C). Gaze-modulation was best fit as including a multiplicative change in response gain rather than simply an additive increase in firing rate, predominantly consisting of increases in gain (127 neurons) rather than decreases (22 neurons; Fig 6D). Thus, the strength of neural responses increased during gaze at the stimulus, by a median of 33% across all neural populations and by 133% in significantly gaze-related neurons (Fig 6D,E). These gain changes were present in all animals and tasks in which gaze was attracted by resolution of uncertainty (Fig S5). Furthermore, in all three areas the mean timecourse of gaze-modulation in these neurons began long before the gaze shift, reaching 30-50% of maximal modulation before the eye movement occurred (Fig 6F). The gaze-modulation began significantly earlier in ACC than icbDS and Pal. This was the case when comparing gaze-modulation latencies based on each area’s mean timecourse of modulation (Fig 6F, both p < 0.01, permutation tests), and also when comparing the latencies of modulation in individual neurons (Fig 6F inset, both p < 0.01, rank-sum tests). Thus, while uncertainty-related signals emerged earliest in Pal (Fig 1), it was their fluctuations in ACC that were first linked to motivation of future gaze behavior.

In sum, these findings indicate that the cortico-BG network can spontaneously change the strength of its information-anticipatory signals. These changes start in the ACC, continue in the BG, and are then immediately followed by gaze shifts toward or away from the object associated with resolution of uncertainty.

### Perturbations of network activity impair information seeking behavior in the manners predicted by network architecture

Given these findings, we hypothesized that information predictive neurons in the basal ganglia have a causal role in motivating gaze shifts to gain information. If so, then temporarily inactivating the basal ganglia subregions that contain these neurons should impair the motivation to seek information. We therefore trained monkeys to perform a task in which gaze shifts were required to gain information (gaze-contingent information task, Fig 7A). Monkeys fixated a spot of light and then continued to fixate during a delay period while a 50% reward CS was presented on either the left or right side of the screen. After the fixation point disappeared (‘go’ signal) the monkey was free to gaze in any manner they chose. On trials with an Info CS, gazing at the CS caused it to be immediately replaced by an informative cue indicating the trial’s outcome (reward or no reward). On trials with a Noinfo CS, gazing at the CS caused it to be replaced by a non-informative cue. Importantly, the monkeys’ gaze at the CS allowed them to gain information about the outcome but did not allow them to influence the outcome itself in any way (i.e. the outcome always occurred a fixed time after the ‘go’ signal regardless of whether and how they gazed at the CS). In addition, we used a fixed-duration delay period between CS onset and the go signal to allow animals to anticipate the moment when information would become available.

**Figure 7.**
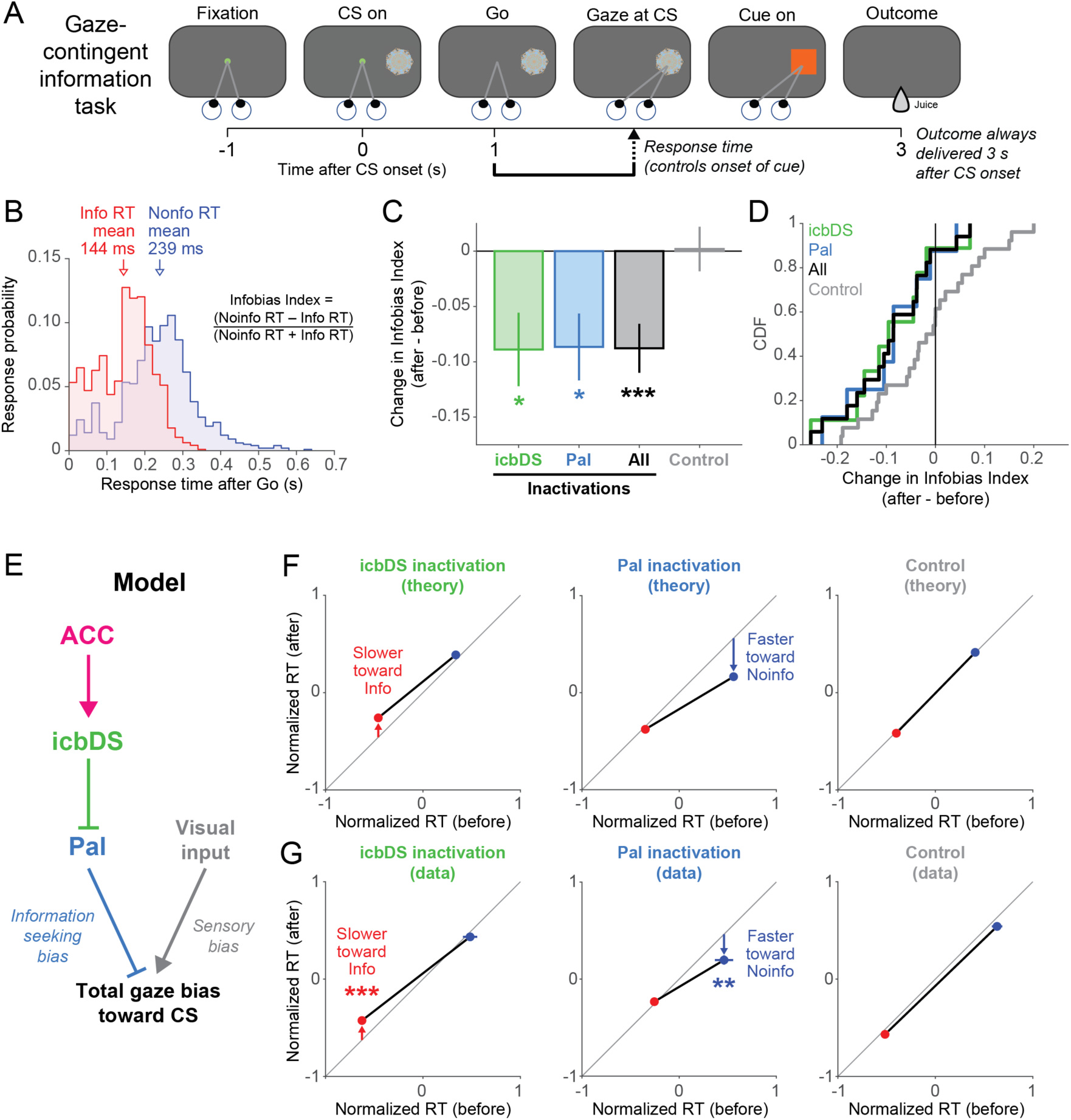
Perturbation of network activity impairs information seeking. (**A**) Gaze-contingent information task. Monkeys were shown a CS, waited for a ‘go’ signal, and then were allowed to gaze at it. Gazing at an Info or Noinfo CS caused it to be replaced with an appropriate informative or non-informative cue. Regardless of whether or when animals gazed at the CS, the outcome was delivered a fixed time after CS onset. (**B**) RT distribution for animal B. The animal had much faster RTs for Info trials (red) than Noinfo trials (blue). The animal often anticipated the time that information would become available, as indicated by the prevalence of anticipatory saccades especially on Info trials (e.g. RTs < 50 ms). This histogram includes all data that was collected when inactivations were not being performed. It shows n=1962/1966 (99.8%) of those RTs; not visible are four outliers from Noinfo trials (RT = 0.741, 0.748, 1.326, 1.335 s). Text indicates the equation for the Infobias Index. (**C**) Muscimol inactivation effect on RTs to contralateral CSs, quantified as the change in Infobias Index (after– before). There are significant reductions in Infobias Index for icbDS inactivations (green), Pal inactivations (blue), and all inactivations (black), but not control sessions (gray). *, **, *** indicate p < 0.05, 0.01, 0.001. Error bars are ± 1 SE. (**D**) Cumulative distributions showing each session’s inactivation effect on the Infobias Index for contralateral CSs. Inactivation sessions consistently reduce the information bias while control sessions do not. (**E**) Schematic model of a mechanism by which the cortico-BG network could govern information seeking gaze behavior. (**F**) Model predictions: the two BG areas should influence information seeking in distinct manners, such that icbDS inactivation slows RTs to obtain information (left, Info CS, red), Pal inactivation speeds RTs that will not obtain information (middle, Noinfo CS, blue), and controls have no effect (right). (**G**) Inactivation results, quantified by comparing normalized RTs (Methods) for the Info CS (red) and Noinfo CS (blue) before vs. after inactivation. icbDS inactivation slowed RTs to the Info CS without any significant effect on Noinfo RTs (left); Pal inactivation speeded RTs to Noinfo CS without any significant effect on Info RTs (middle); control sessions had no significant effect on RTs to either CS (right). Error bars are ± 1 SE.

Indeed, animals had strongly anticipatory behavior, at times shifting their gaze onto the CS at short latencies before they could have perceived and reacted to the ‘go’ signal (Fig 7B, response times (RTs) < ∼100 ms). Monkeys were highly motivated to seek information, shown by their much faster RTs to shift their gaze onto Info CSs than Noinfo CSs (Fig 7B). We quantified the response bias favoring the info CSs with an “Infobias Index” (Fig 7B) which was significantly positive in every session for every animal (n=43/43 sessions, all p < 0.05, permutation tests; Methods). Furthermore, we recorded from a subset of neurons using both the original and gaze-contingent versions of the information task and found that they had similar information-anticipatory activity in both tasks (Fig S9). This confirmed that information-anticipatory activity was present and well-positioned to regulate information seeking in this context where gaze shifts are instrumental actions required to gain information.

Consistent with the lateralized functions of basal ganglia circuitry ^30, 31^, we predicted that unilateral inactivations would reduce information seeking behavior directed toward objects in contralateral space (Supplementary Note). Indeed, unilateral injections of muscimol, a GABAa agonist, into either icbDS or Pal in the vicinity where information-anticipatory neurons were recorded caused the information seeking response bias to be significantly reduced in contralateral space (Fig 7C,D; icbDS, n=9 sessions, p = 0.028; Pal, n=8 sessions, p = 0.031; all inactivations, n=17, p = 0.0009; permutation tests, Methods; see Figs S2,S10 and Table S2 for injection sites and effects in each session). No significant change was observed in a control dataset consisting of sham and saline injections (Fig 7B, gray, n=26 sessions, p = 0.93, permutation test). Thus, relative to control sessions, inactivations caused the information seeking bias to be significantly reduced (p = 0.005, permutation test; Methods). In addition, inactivations had no significant effect on information seeking for ipsilateral CSs (all p > 0.4, permutation test, Fig S11). Thus, relative to control sessions, inactivations caused the information seeking bias to become lateralized – significantly shifted away from the contralateral side (p = 0.019, permutation test, Methods; Fig S11).

We further investigated the mechanism by which icbDS and Pal activity promote information seeking. Our data suggest that icbDS and Pal have reciprocal inhibitory connections and tend to encode information predictions in opposite manners, with icbDS neurons activated and Pal neurons commonly inhibited (Figs 1,2,S1). We therefore hypothesized that icbDS and Pal activity have opposite influences on motivated gaze behavior, such that information-oriented gaze shifts are motivated by icbDS activity and suppressed by Pal activity. A simple network model implementing this hypothesis (Fig 7E) reproduced the observed behavior on control sessions and predicted that inactivating icbDS and Pal should impair information seeking in distinct manners (Fig 7F). The icbDS is primarily active during the Info CS, so inactivation should slow gaze shifts to the Info CS while leaving responses to the Noinfo CS relatively intact (Fig 7F, left). Conversely, Pal is normally inhibited during the Info CS, so inactivation should leave responses to the Info CS relatively intact while speeding gaze shifts to the Noinfo CS (Fig 7F, middle). Both of these predictions were borne out in the data. Inactivation of icbDS slowed gaze shifts to the Info CS but did not significantly change RTs to the Noinfo CS (Fig 7G, left, Info CS p = 0.0003, Noinfo CS p = 0.64, rank-sum tests; significantly different change in normalized RT for Info CSs vs Noinfo CSs, indicated by a significant interaction term in a 2-way ANOVA using the factors CS type (Info or Noinfo) and epoch (pre- or post-injection), F_1,1808_ = 7.12, p = 0.008; Methods). Conversely, inactivation of Pal speeded gaze shifts to the Noinfo CS but did not significantly change RTs to the Info CS (Fig 7G, middle, Info CS p = 0.17, Noinfo CS p = 0.001, rank-sum tests; significantly different change in normalized RT for Info vs Noinfo, 2-way ANOVA, F_1,1955_ = 7.89, p = 0.005). Thus, a direct comparison between inactivations of the two areas revealed significantly different effects on behavior. Inactivation of icbDS caused significant slowing of normalized RTs relative to inactivation of Pal (permutation test, p = 0.0043). This occurred due to inactivations changing RTs in the predicted manner (significant effect of icbDS slowing saccades to the Info CS and Pal speeding saccades to the Noinfo CS, permutation test, p < 0.0001; Methods) and not in the orthogonal manner (no significant effect of icbDS slowing saccades to the Noinfo CS and Pal speeding saccades to the Info CS, permutation test, p = 0.3915). These changes in RTs included changes in anticipatory saccades, consistent with a disruption of information-anticipatory activity in these areas (Fig S12). Thus, icbDS activity motivated gaze shifts to gain information, while Pal activity suppressed motivation to gaze at objects that would not yield information.

Importantly, these inactivation effects on information seeking were not caused by generalized effects on overall motivation to perform the task, which could potentially be affected by inactivation of striatum and pallidum circuitry involved in primary reward seeking behavior. Specifically, icbDS inactivations slowed RTs to the Info CS without reducing measures of general motivation, while Pal inactivations speeded RTs to the Noinfo CS without increasing measures of general motivation (Fig S11). If anything, Pal inactivations speeded RTs to the Noinfo CS in spite of a modest reduction in general motivation to perform the task (Fig S11), consistent with previous Pal inactivations ^32–38^.

## DISCUSSION

### A neural network for information seeking

Our work demonstrates the existence of a neural network responsible for motivating actions to resolve uncertain situations by seeking knowledge about future rewards. Previous studies have identified cortical and basal ganglia networks that make conventional predictions about when future rewards are available and motivate behavior to seek those rewards ^13, 30, 39, 40^. Indeed, our monkeys had strong tendencies to gaze at visual stimuli based on their reward value. However, the subset of ACC-icbDS-Pal neurons we report here have relatively little response to reward value and hence their primary function is not likely to be control of such reward value-oriented behavior. Instead, they have a quite distinct function: they predict when information will become available to resolve reward uncertainty and motivate gaze behavior to obtain that information. This information seeking gaze behavior can be even more potent than the attraction of gaze to primary reward: our monkeys gazed much more avidly at the CS that provided informative cues than at any other stimulus, even CSs and cues that were associated with *double* its expected reward value. Our data show that this information seeking gaze behavior is tightly coupled to the cortico-BG network – fluctuations in the strength of its information-anticipatory signals are followed by immediate gaze shifts toward or away from the information-related stimulus, and artificial perturbations of its activity interfere with information seeking in the manner predicted by its neural signals and wiring.

Our data is crucial evidence for theories of reward learning, overt attention, and economic decision making which propose that objects and events in the world are assigned salience both by neural systems that track primary reward value and its uncertainty ^41–44^, and by a system that anticipates information to resolve uncertainty ^2, 6, 45–47^. Furthermore, our data demonstrates a neural mechanism through which information seeking can compete and interact with primary reward to drive ongoing behavior ^7, 9, 10, 48^.

In fact, information seeking goes hand-in-hand with primary reward seeking in natural environments. Most experimental studies of reward seeking begin with the presentation of a cue stimulus (or an environmental context) that tells the subject what reward to predict and what actions are needed to obtain it. However, rewards in natural environments can be scarce and uncertain, and fully predictive reward cues rarely come for free or materialize from thin air. In these situations, organisms must first seek and obtain information about the rewards that are available in their environment; only then can they predict the value of those rewards and use that value to motivate reward seeking behavior. In this sense, the cortico-BG network for information seeking may be critical to ensure that organisms seek out the reward-related cues in their environment that are necessary for the proper operation of the well-known networks that predict and seek primary rewards ^13, 30, 39, 40, 49^. Indeed, information-related neurons in all three areas were intermixed with other neurons that encoded the reward value of stimuli and outcomes, as expected from previous studies of these areas ^13, 22, 30, 39, 40, 49–52^. This suggests that information- and primary reward-related neurons are well-positioned to support each other’s computations. For instance, information-anticipatory activity in our tasks can be interpreted as ramping up to the expected time of a large reward prediction error (due to being informed of a pleasing or disappointing future outcome). Thus, while most uncertainty-related neurons in these areas do not encode reward prediction errors themselves ^22^, their activity could prepare local reward-processing networks to handle upcoming reward prediction error or surprise signals, processes in which ACC, dorsal striatum, and pallidum have been implicated ^40, 53–57^.

Importantly, monkeys expressed a strong preference for the information consistent with it having a high subjective value ^9^ even though the information did not have any ‘objective’ value, in the sense that it did not help the monkeys take action to gain more juice reward (a quantity called the “value of information” or “value of exploration” in decision theory and reinforcement learning ^58, 59^). An important area of future research will be how these two factors combine to guide behavior and whether they are implemented by shared or distinct neural networks. Notably, our study identifies a network that motivates seeking of a specific type of information (i.e. the presence or absence of an upcoming reward). It remains unknown whether this network could also motivate seeking of other types of information, such as information about instrumental contingencies (i.e. what action will be required to gain the reward^60^). In natural environments these types of information are likely to synergize with each other: if an agent has a subjective preference to resolve uncertainty in order to better *predict* future rewards, then the agent will likely be better at learning the objective value of resolving that uncertainty to better take actions and *control* rewards.

### Anterior cingulate cortex and information seeking behavior

While information-anticipatory signals were present in all three areas of the cortico-BG network, each area also had distinct features suitable for unique contributions to information seeking. Notably, fluctuations in ACC information signals were the earliest predictor of future behavior. ACC information signals changed several hundred milliseconds before gaze shifts, while BG signals changed more proximally to behavior. This finding supports and extends theories that ACC is especially important for motivating behavioral shifts to explore available prospects and learn their reward value and other properties ^61–63^, tracking their level of uncertainty and how it evolves over time as beliefs are updated in response to surprising outcomes ^12, 62, 64–68^, and using this information to decide how to control future cognition and behavior ^62, 69^. In particular, while it is well acknowledged that the ACC needs to receive a broad array of reward- and uncertainty-related information to perform these functions ^62, 69^, our data indicates that the ACC is not a mere passive recipient of this information; rather, the ACC is tightly linked to the emergence of motivational drive to actively seek out that information from the environment. Information-anticipatory activity would be especially useful from the perspective of theories that the ACC regulates foraging ^62^, because knowing the properties of the potential sources of reward in the environment is a fundamental requirement for making efficient foraging decisions. It would also be useful from the perspective of theories that the ACC regulates cognitive control ^69^, because one of the most crucial times to bring cognitive resources online is in preparation for receiving a new piece of information, in order to process it quickly and prepare an effective response.

### Implications for basal ganglia circuitry underlying motivated behavior

In addition, our findings indicate that information seeking behavior is motivated by a BG circuit mechanism that is analogous but distinct from the BG circuits that motivate conventional reward-seeking behavior. There are two key parallels. First, behavior-related fluctuations in BG information signals follow fluctuations in cortex and are proximal to behavioral gaze shifts. This is consistent with classic theories of cortico-BG circuits ^25^ and work on cortex-striatum interactions ^70–72^ suggesting that cognitive and motivational signals can be computed in cortex and then sent to BG whence they are processed and used to guide behavior. Second, the specific functions of each BG subregion in information seeking are consistent with classic BG circuit motifs (Fig 7E): icbDS and Pal neurons commonly encode information signals with opposite signs and these areas have opposite causal influences on behavior, such that icbDS activity speeds gaze shifts to gain information while Pal activity slows gaze shifts that will not provide information. This resembles analogous findings for BG areas involved in primary reward seeking: antagonizing D1 receptors in visuomotor dorsal striatum slows gaze shifts to gain large juice rewards ^73^, while inactivation of Pal speeds gaze shifts to gain small juice rewards ^35^.

Importantly, however, the BG mechanisms underlying physical reward- and information-oriented behavior are at least partially distinct at the neuronal and behavioral levels: first, when animals avidly gazed at the Info CS in anticipation of viewing the informative cue, they had near-zero licking behavior indicating that they were not anticipating juice reward; second, the cortical and BG neurons we identified that are linked to information-anticipatory behavior primarily anticipated the moment of gaining information, not the moment of gaining juice reward; third, inactivation effects on information seeking could not be explained as a result of generalized effects on juice reward seeking. Thus, this cortico-BG network appears to be specially focused on online information seeking behavior. This is in contrast to other BG circuits and interconnected areas involved in reward prediction errors and reinforcement, which commonly encode information and primary reward in a common currency ^6, 11^.

In addition, whereas classic theories of cortico-BG circuits classify Pal as an output structure ^25, 74^, our data extends recent results ^37, 38, 75^ by showing that Pal in fact responds earliest to uncertainty-related events. This is consistent with theories that BG rapidly selects salient stimuli to be used to guide future behavior ^76, 77^, perhaps based on input from areas specialized for rapid assessment of objects and their incentive properties, such as the amygdala^78^ and brainstem^79^. Our data supports a scenario in which (A) Pal rapidly signals a rough assessment of reward uncertainty; (B) ACC and icbDS first signal the precise graded level of uncertainty; (C) the resulting representation of uncertainty in all three areas ramps up to the time of its resolution by information, and drives ongoing information seeking behavior.

Given the link between the ACC-icbDS-Pal network and information seeking behavior, variations in the network’s activity could be responsible for the natural variations in information seeking behavior that are commonly found across individuals ^5, 10, 12^ and tasks ^3, 80^. In the same vein, it is notable that ACC and BG are implicated as sites of dysfunction and targets for treatment in human disorders of motivated behavior (such as obsessive-compulsive disorder ^81, 82^, Parkinson’s disease ^83^, and drug addiction ^84^) that are known to affect reward- and uncertainty-related behavior ^85–90^. Our results raise the possibility that these disorders and treatments may also affect the motivation to seek information about future events. While this has been little studied, there is evidence that Parkinson’s disease reduces the motivation to gather information needed for upcoming decisions ^91^ and impairs learning from early access to information about uncertain outcomes ^92^. Taken together, our work provides a foundation for understanding the neural network mechanisms by which information is detected, predicted, and used to motivate behavior.

## METHODS

### General procedures

Four adult male rhesus monkeys (*Macaca mulatta*) were used for behavioral, recording, and inactivation experiments (Animals B, R, Z, and W). All procedures conformed to the Guide for the Care and Use of Laboratory Animals and were approved by the Washington University Institutional Animal Care and Use Committee. A plastic head holder and plastic recording chamber were fixed to the skull under general anesthesia and sterile surgical conditions. The chambers were tilted laterally by 35-40° and aimed at the anterior cingulate and the anterior regions of the basal ganglia. After the animals recovered from surgery, they participated in the experiments.

### Data acquisition

While the animals participated in the behavioral tasks we recorded single neurons in the anterior cingulate cortex (ACC), internal capsule-bordering regions of the dorsal striatum (icbDS), and anterior pallidum including the ventral pallidum and the anteriormost part of the globus pallidus internal segment (Pal). Electrode trajectories were determined with 1 mm-spacing grid system and with the aid of MR images (3T) obtained along the direction of the recording chamber. This MRI-based estimation of neuron recording locations was aided by custom-built software (PyElectrode ^93^). In addition, in order to further verify the location of recording sites, after a subset of experiments the electrode was temporarily fixed in place at the recording site and the electrode tip’s location in the target area was verified by MRI (Fig S2).

Single-unit recording was performed using glass-coated electrodes (Alpha Omega). The electrode was inserted through a stainless-steel guide tube and advanced by an oil-driven micromanipulator (MO-97A, Narishige). Signal acquisition (including amplification and filtering) was performed using Alpha Omega 44 kHz SNR system. Action potential waveforms were identified online by multiple time-amplitude windows with an additional template matching algorithm (Alpha-Omega). Neuronal recording was restricted to single neurons that were isolated online. Neuronal and behavioral analyses were conducted offline in Matlab (Mathworks, Natick, MA).

Eye position was obtained with an infrared video camera (Eyelink, SR Research). Behavioral events and visual stimuli were controlled by Matlab (Mathworks, Natick, MA) with Psychophysics Toolbox extensions. Juice, used as reward, was delivered with a solenoid delivery reward system (CRIST Instruments). Juice-related licking was measured and quantified using previously described methods ^94^.

### Behavioral tasks

We analyzed data recorded from several behavioral tasks which can be grouped into two major categories: standard uncertainty tasks and information tasks.

The *standard uncertainty tasks* are described in detail in previous work ^21–23^. They each used a distinct set of fractal visual CSs with different associated outcomes. However, they all shared the following general outline. Animals were presented with a small white circular trial start cue at the center of the screen. In some tasks animals were required to fixate the trial start cue for a fixed duration (typically 0.5-1 s) for the trial to continue; if they failed to fulfill this requirement within a grace period (typically 5 s) the trial would be considered an error, they would receive a timeout, and the trial would repeat. In other tasks animal were not required to fixate the trial start cue; it was simply shown for a fixed duration (typically 1 s). After the trial start period, the trial start cue disappeared and a fractal visual conditioned stimulus (CS, 2° radius) appeared on the screen for a fixed duration (2.5 s). The CS was randomly positioned at one of three locations: the center of the screen, the left side of the screen, or the right side of the screen (at 10° or 12.5° eccentricity). In some sessions only the left and right locations were used. Animals were not required to gaze at or interact with the CS in any way. At the end of the CS period, the CS disappeared and simultaneously the trial’s outcome was delivered. Finally, there was an inter-trial interval during which the screen was blank (typically randomized between 1-8 s, with different durations for different animals and tasks). In some sessions, a small fraction of inter-trial intervals included the unexpected presentation of salient events, which could be appetitive (juice), aversive (an airpuff, ∼35 psi, delivered through a narrow tube placed ∼6-8 cm from the face ^22^), or audiovisual (an auditory tone sounding and the screen flashing white).

The standard uncertainty tasks primarily differed in their CSs, outcomes, and block structure:

- Task A ^21, 22^: Trials were presented in two distinct blocks. In the Probability block, there were five CSs associated with 0, 25, 50, 75, and 100% probabilities of 0.25 mL juice. In the Amount block, there were five CSs associated with 100% probability of 0, 0.0625, 0.125, 0.1875, and 0.25 mL juice. Hence for each CS in the Probability block there was a matched CS in the Amount block that was associated with an identical mean amount of juice, but for which the outcome was certain rather than probabilistic. Each block consisted of 20 trials (4 presentations of each of its 5 CSs, shuffled in a randomized order). The two blocks were presented repeatedly in an alternating manner, with each block continuing until its 20 trials were correctly completed and then immediately transitioning into the other block.
- Task B ^21, 23^: Same as task A, except it used three Probability CSs (0, 50, 100%) and the three corresponding Amount CSs (0, 0.125, 0.25 mL), and each block consisted of 6 or 9 trials (2 or 3 presentations of each of its 3 CSs). In some sessions, blocks also included interleaved choice trials in which two CSs were presented and animals were allowed to choose between them with a saccade; our analysis here is of non-choice trials.
- Task C ^21, 22^: Trials were presented in two distinct blocks. In the Appetitive block, there were three CSs associated with 0, 50, and 100% probabilities of 0.4 mL juice. In the Aversive block, there were three CSs associated with 0, 50, and 100% probabilities of airpuffs. Each block consisted of 12 trials (4 presentations of each of its 3 CSs).
- Task D ^22^: There were 9 CSs. Four CSs were associated with 25, 50, 75, and 100% probabilities of 0.4 mL juice. Four other CSs were associated with 25, 50, 75, and 100% probabilities of airpuff. One final CS was associated with no outcome (i.e. 0% probability of both reward and airpuff). The CSs were presented in a pseudorandom order.
- Task E: Three CSs were associated with 0, 50, and 100% probabilities of 0.25 mL juice. The CSs were presented in a pseudorandom order.

The *information tasks* follow the design in Fig 2A or are variants of this procedure. The task began with the appearance of a small circular trial start cue at the center of the screen which animals were required to fixate for a fixed duration (typically 0.5 or 1 s). The trial start cue then disappeared and was followed in succession by a CS (2° radius) that was displayed for a fixed duration (typically 1 s), which was then replaced by a cue of the same width and height at the same location that was displayed for a fixed duration (typically 2 s). The cue then disappeared, and simultaneously the outcome was delivered. The trial then completed with a 1 s inter-trial interval. The CSs were presented randomly on either the left or right side of the screen (10°). The CSs came in two types. The *Info CSs* predicted juice reward (typically 0.25 mL) with different probabilities (e.g. 0, 50, and 100%), and were followed by one of two informative cues whose color indicated the trial’s outcome. The *Noinfo CSs* also yielded juice reward with matched size and probability, but were followed by one of two non-informative cues whose colors were randomized on each trial and hence did not convey any information about the trial’s outcome. In some sessions Noinfo CSs were followed by a single non-informative visual cue; there was no apparent difference in behavior or neural activity between sessions with one or two non-informative cues and hence their data was pooled. The CSs were presented in a pseudo-random order.

We collected data using the following information tasks:

- Task IA: the task shown in Fig 2A, used to record the majority of neurons. There are three Info CSs and three Noinfo CSs, respectively associated with 0, 50, and 100% probability of reward.
- Task IB: the *gaze-contingent information task*, shown in Fig 7A. This followed the same general procedure as task IA, but with a few modifications. The trial start cue remained visible for a fixed duration after CS onset during which animals were required to maintain fixation on the trial start cue (typically for 1 s in animal B, 0.25 s in animal R, and 0.5 or 1 s in animal Z). Fixation breaks were treated as errors: the screen went blank, there was a 1-2 s penalty delay period, and then the trial repeated from the beginning. After the fixation period, the trial start cue disappeared and animals were free to move their eyes. The task then detected the first moment when animals gazed at the CS, defined as the eye position entering a square window centered on the CS (i.e. when horizontal and vertical eye positions were within 4° of the center of the CS). If animals gazed at an Info CS it was immediately replaced with the appropriate informative cue; if they gazed at a Noinfo CS it was immediately replaced with a non-informative cue; if they did not gaze at a CS, no cue was shown. Importantly, regardless of their gaze behavior, all stimuli disappeared and the outcome was delivered at the same, fixed time after CS onset on all trials in the session (typically 3 s). Thus, gazing at the CS gave animals access to the cues but did not give them earlier access to the juice reward. In the version of this task used for neuronal recording, we used the same visual CSs as in Task IA. Tasks IA and IB were typically pseudo-randomly interleaved in a trial-by-trial manner. At the start of each trial the current task was indicated to the animal by the color of the fixation point (white for IA, green for IB). In the version of this task used for inactivation experiments and controls, there were only two CSs – an Info CS and a Noinfo CS – that were both associated with 50% probability of 0.25 mL juice reward. This was to minimize the possibility that gaze behavior to the CSs could be influenced by different reward expectations or reward prediction errors induced by CS onset, by ensuring that that the probability, amount, and timing of juice reward was identical for all CSs on all trials.
- Task IC: used to record a subset of neurons in animal B. Similar to task IA but instead of 3 types of CSs (0, 50, and 100% probability of reward) there were 10 types of CSs, which were respectively associated with 25, 50, 75, or 100% probabilities of reward, with the equivalent probabilities of punishment, with no outcome (i.e. 0% probability of either reward or punishment), or with 50% probability of either reward or punishment. Info and Noinfo trials were presented in separate blocks. There were also minor changes in task timing and visual stimuli: the CS and cue periods were 1 s and 2.25 s duration, cues were presented as colored rectangles around the CS rather than as squares replacing the CS, and the CSs remained onscreen during the first 1.5 s of the cue period.
- Task ID: used to record a subset of neurons in animal R. Similar to task IA but with only 3 total CSs: Info 50% chance of 0.38 mL reward, Noinfo 50% chance of 0.38 mL reward, and a Certain CS which yielded a 100% chance of 0.19 mL reward. For this task, uncertainty signals were defined on Info trials as the ROC area comparing Info 50% CS trials to Certain CS trials, and on Noinfo trials as the ROC area comparing Noinfo 50% CS trials to Certain CS trials. There were also minor changes in task timing and visual stimuli: the CS and cue periods were 1.5 s and 1.5 s in duration, each individual CS was associated with two distinct cues, and the cues were square-shaped fractal stimuli rather than colored rectangles.

Information-related neural activity was typically similar in the standard and gaze-contingent versions of the information task (IA and IB), e.g. activity ramping up to the time the informative cue would become available and to the time a non-informed outcome would be delivered. Note, however, that the task design of the gaze-contingent task potentially induced a link between gaze and receipt of information: gaze behavior was not completely ‘free’ because it was required if the animal wanted to produce the cue, and the cue appeared with variable timing depending on the animal’s behavior. Therefore, to be conservative, data from the gaze-contingent task was excluded from all of our analysis of neural-behavioral links (Fig 5-6), and was only used for a subset of other analyses. First, we analyzed data from tasks IA and IB separately to compare them to each other in our supplementary analysis testing whether information signals are altered when access to information is explicitly contingent on gaze. Second, a neuron’s uncertainty coding in the time window 0.5 sec before outcome delivery was calculated by pooling data from tasks IA and IB, because by that time in the trial the task was no longer gaze-contingent because animals had already revealed the cue on the great majority of trials. All remaining analyses, including our main analysis of information signals (Fig 2-3), were restricted to data from task IA, except for a small number of neurons for which data from task IA was not available because they were only recorded in task IB (n=4 ACC neurons in monkey Z).

Analysis of neural activity in Fig 1 and S1 uses data from all neurons recorded using standard uncertainty tasks in which CSs were associated with all five reward probabilities (tasks A and D). Analysis of neural activity during information tasks (Fig 2,3) uses data from neurons recorded using an information task. Analyses of neural activity related to gaze behavior (Fig 5,6) pooled data from all neurons recorded in all tasks that contained trials of the types specified in the analyses (e.g. Fig 5 used neurons with data from 0, 50, and 100% conditions on both Info and Noinfo trials), except for the 4 ACC neurons described above from animal Z recorded only in the gaze-contingent information task, and 6 Pal neurons from animal W recorded in task A for which the gaze measurements were excessively noisy due to an error in configuring the eye tracker. If a neuron was recorded in multiple tasks, its data was fitted separately for each task and then pooled over tasks. Specifically, for each neuron, the model’s total log likelihood was calculated as the sum of the log likelihoods from the individual tasks, and the neuron’s gaze-aligned activity (e.g. Fig 5), fitted gaze-related gain change and fitted timecourse of gaze-modulation and latency of gaze-modulation (e.g. Fig 6) were calculated for each task and then averaged over tasks.

### Muscimol injections

On *muscimol injection* sessions, a 33-gauge cannula was inserted through a 23-gauge guide tube into a grid hole and to a depth previously identified to be in icbDS or Pal and to contain information-related neurons (see Fig S2 and Table S2 for coordinates of all injection sites). The other end of the cannula was connected to a 10 µL Hamilton syringe. Behavioral data from the gaze-contingent information task was collected in blocks of 70-150 correct trials. Before the injection we collected a ‘pre-injection’ behavioral data set from the animal performing the task, typically for one block (median: 96 correct trials, standard deviation: 24, range: 48-164). After recording the baseline data, we used a manual syringe pump (Stoelting) or automated syringe pump (Harvard Apparatus) to inject muscimol dissolved in saline. Muscimol concentrations were 8 mg/mL, injection rates were typically 0.1 µL/min (range: 0.09-0.2), and injection volumes are reported for each session in Table S2. After each injection we collected a ‘post-injection’ behavioral data set (median: 303 correct trials, standard deviation: 157, range: 43-839). All pre- and post-injection blocks of data were included in our analysis regardless of the animal’s response times or other gaze behavior, as long as the animal remained engaged in the task (i.e. generally initiating trials quickly and performing them correctly). On two sessions pre-injection data from the same day was not available, so to obtain a comparable baseline we used the first block of behavioral data collected from the same animal on days immediately before or after the session. On *saline injection* control sessions, the same procedure was followed except that only the saline vehicle was injected (with the same volumes previously used for muscimol injections). On *sham* control sessions, the same procedure was followed to mimic the procedure for injection experiments in every detail, except that no cannula was inserted and no injection was made. Specifically, we (1) set up the injection equipment above the animal, (2) mounted the microdrive, (3) closed the experimental booth and waited for the standard period of time that would be required to use the microdrive to advance the tip of the cannula to the target area, (4) started the behavioral task and ran it for the same duration and schedule as the ‘before’ condition on injection days, (5) stopped the task, entered the booth, and turned on the motor of the injection device to mimic the sounds and duration of time spent performing an injection, (6) turned on the task again and ran it for the same duration and schedule as the ‘after’ condition on injection days. The only difference was that in injection experiments the cannula was loaded into the microdrive and advanced to the target area, while in sham experiments it was not. Thus, sham sessions act as a control for the possibility that the observed alternations in information seeking were due to a generalized effect of the experimental procedure.

### Data analysis

All statistical tests were two-tailed unless otherwise noted. Neurons recorded in the standard uncertainty tasks were included in our dataset if they showed significant responsiveness to uncertainty (activity on uncertain reward CS trials significantly different from both 0% reward CS trials and 100% reward CS trials, rank-sum tests, both p < 0.05 and both differences with the same sign). For this purpose, activity was measured in a broad time window encompassing the CS period in order to avoid making any assumptions about the time course of neural responses (0.1-2.5 s after CS onset). The neuron’s sign of uncertainty coding was defined as +1 if its ROC area for discriminating between uncertain reward CS trials vs pooled data from 0% and 100% reward CS trials was > 0.5, and defined as −1 if its ROC area was < 0.5. Similarly, for the information task, to avoid making any assumptions about the nature of Info-, CS-, or cue-related activity, neurons were included in our dataset if their firing rates in a time window 0.5 sec before outcome delivery significantly discriminated between Noinfo certain vs. uncertain reward trials (ROC area ≠ 0.5, p < 0.05, rank-sum test), and its sign of uncertainty coding was set based on this activity in the analogous manner.

Neural activity was converted to normalized activity as follows. Each neuron’s spiking activity was smoothed with a causal exponential kernel (mean = 30 ms) and then z-scored and sign-normalized using the following procedure. The neuron’s average activity timecourse aligned at CS onset was calculated for each condition (defined here as each combination of CS and cue). These average activity timecourses from the different conditions were all concatenated into a single vector, and its mean and standard deviation were calculated. Henceforth, all future analyses converted that neuron’s firing rates to normalized activity by (1) subtracting the mean of that vector, (2) dividing by the standard deviation of that vector, (3) multiplying by the neuron’s sign of uncertainty coding. Thus, normalized activity of +1 in a given task condition means that the neuron’s firing rate deviated away from its average firing rate in the same direction that it responded to uncertainty, by an amount equivalent to 1 SD of its overall distribution of average firing rates during the task.

Neural uncertainty signals were calculated in specific time windows (e.g. pre-cue, pre-outcome, etc.) as the ROC area for distinguishing activity on uncertain reward CS trials (25, 50, and 75%) from pooled data from 0% and 100% certain reward trials. In the information tasks uncertainty signals were calculated separately for Info and Noinfo trials. To visualize their timecourses, they were calculated on neural activity at millisecond resolution after activity was smoothed with a gaussian kernel (SD = 50 ms) and sign-normalized based on the neuron’s sign of uncertainty coding on Noinfo trials in a 0.5 s pre-outcome window (Fig 3). The Informative Cue Anticipation Index was defined as the difference between its uncertainty signal for Info and Noinfo trials in a 0.5 s pre-cue time window (or for Fig 3A, visualized by calculating it separately at each time point). Hence the index was positive if a neuron had a higher uncertainty signal in anticipation of Info CSs, and negative if a neuron had a higher uncertainty signal in anticipation of Noinfo CSs. The Uncertain Outcome Anticipation Index was defined as the difference between its uncertainty signals computed on two different time windows on Noinfo trials: a 0.5 s pre-outcome window, and a 0.5 s post-cue window (0.15-0.65 s after cue onset). Hence the index was positive if a neuron’s uncertainty signal grew more positive between the cue and outcome, and negative if it grew more negative between the cue and outcome. Neurons were classified as information-responsive if their Informative Cue Anticipation Index was significantly different from 0 (p < 0.05, permutation tests conducted by comparing the index calculated on the true data to the distribution of indexes calculated on 20000 permuted datasets which shuffled the assignment of trials to Info and Noinfo conditions). Neurons were classified as having a significant Uncertain Outcome Anticipation Index using the analogous permutation test (p < 0.05, shuffling the assignment of data to the post-cue and pre-outcome time windows). For analysis of information-oriented gaze behavior, the same two indexes were calculated for each neuron except that instead of using neural data they used the behavioral gaze data from the last millisecond in each time window (equal to 1 for milliseconds when the animal’s gaze was classified as being in the stimulus window and 0 otherwise). Finally, to plot the timecourse of uncertainty signals from the population including neurons with different signs of uncertainty coding, the normalized uncertainty signal (Fig 3C) was calculated for each neuron using the equation: U_N_ = 0.5 + (U – 0.5)*S, where U_N_ is the normalized uncertainty signal, U is the uncertainty signal, and S is the neuron’s sign of uncertainty coding (−1 or +1). Thus, neurons with a positive sign of uncertainty coding (e.g. excited by uncertainty) had their uncertainty signals left intact, while neurons with a negative sign of uncertainty coding (e.g. inhibited by uncertainty) had their uncertainty signals ‘flipped’. This ensured that each neuron had a positive normalized uncertainty signal in the time window and task condition that was used to classify its sign of uncertainty coding.

#### Latency of uncertainty coding

In standard uncertainty tasks, each neuron’s smoothed normalized activity aligned at CS onset was further smoothed with a 101 ms causal boxcar kernel and then tested at each millisecond starting 50 ms after CS onset for whether it met the following criteria: (1) highly significant ROC area for distinguishing pooled data from uncertain reward CSs from the certain 0% reward CS (p < 0.005), (2) highly significant ROC area for distinguishing pooled data from uncertain reward CSs from the certain 100% reward CS (p < 0.005); (3) both ROC areas have the same ‘sign’ (i.e. both > 0.5 indicating activation by uncertainty or < 0.5 indicating inhibition by uncertainty). A neuron’s uncertainty coding latency was defined as the first millisecond after which it met these criteria for at least 24 consecutive milliseconds. In the information tasks, latencies were calculated in this manner separately for Info trials and Noinfo trials, and the neuron’s overall latency was set to be the shorter of the two. In information task ID, the first criterion was not applied because there was no task condition with a 0% chance of reward. See Fig S1 for the latencies and full ROC timecourses in all neurons with detected latencies. This method was chosen to produce latencies that resemble those seen in raw traces of neural activity, but the same key result (i.e. Pal having shorter latency than ACC and icbDS) was found with other latency detection methods (e.g. different smoothing methods, significance criteria, required number of consecutive time bins,etc). Each area’s latency was defined as the shortest latency of its single neurons after excluding the shortest 1% of single neuron latencies (rounding up) to make the analysis robust to a small number of false positives. Area latencies were compared by testing whether the difference between their latencies was significantly different from that expected by chance (p < 0.05, permutation test, conducted by comparing the latency difference calculated on the true data to the distribution of latency differences calculated on 20000 permuted datasets which shuffled the assignment of neurons between the two areas being compared).

#### Rough vs graded uncertainty coding

In standard uncertainty tasks, a neuron’s rough uncertainty activity was calculated as the difference in normalized activity between pooled data from all uncertain reward CSs and pooled data from the certain 0% and 100% reward CSs. Its graded uncertainty activity was calculated as the difference in normalized activity between data from the uncertain 50% reward CS and pooled data from the uncertain 25% and 75% reward CSs. Neurons were classified as having significant graded uncertainty coding if their graded uncertainty activity was significantly different from 0 (p < 0.05, rank-sum test). Areas were classified as having graded uncertainty coding if the number of neurons with significant graded coding was significantly different from chance levels (p < 0.05, binomial test).

#### Relating neural activity to gaze

Gaze was defined as being on the stimulus if it was within a small circular window (3° radius) centered on the trial’s CS location. We observed that there was modest but noticeable variance in eye tracker calibration from session to session. For instance, in some sessions the measured gaze location was consistently slightly to the left of the CS at all CS locations, while in other sessions it was slightly to the right. To correct for this, for each neuron and for each CS location, we centered the gaze window on the peak of a smoothed 2D histogram of eye positions collected from all milliseconds when the eye was within an 8° radius of the theoretical center of the CS (using gaze data from all trials that presented a reward-associated CSs and all times from 0.2 s after CS onset to 0.1 s after CS offset). This produced a good match between the gaze window used for analysis and the typical gaze positions around each CS location.

In our first analysis, we tested whether neural activity in each condition (defined as each combination of CS and cue) was altered depending on whether the gaze was on the CS, in a 0.5 s time window immediately before the animal was going to be informed of the outcome. For the standard uncertainty tasks, this was a pre-outcome window. For the information tasks, this was a pre-cue window on Info trials and a pre-outcome window on Noinfo trials; results were calculated separately for these two windows and then averaged. For each neuron and each condition, we did the following procedure. We calculated the neuron’s mean normalized activity in the time window at one millisecond resolution, separately for each gaze state (on or off). We then found all time points at which there was valid data for both gaze states (i.e. milliseconds where there was at least one trial with gaze on and one trial with gaze off). We then calculated the mean normalized activity for each time point separately for the ‘gaze on’ and ‘gaze off’ states, and then averaged across time points. This resulted in two measurements for each condition: the neuron’s average activity when gaze was on the CS, and its average activity at the same time points when the gaze was off the CS. This analysis was performed on all neurons where there was trial-to-trial variance in gaze behavior before receipt of information (i.e. at least one millisecond in the time window in which there was at least one trial with gaze on and one trial with gaze off) for all of the CSs. We then tested whether the mean difference in activity between the two gaze states was significantly different from 0 across the neural population (p < 0.05, signed-rank test). Note that this analysis essentially asks how neural activity differs across behavioral gaze states. We could have done an equivalent analysis in terms of the reverse relationship, by asking how well gaze states can be predicted from neural activity (e.g. in terms of decoding accuracy). However, our hypothesis was not that these neural populations have a 1:1 relationship with gaze states. Our hypothesis was that neural activity is linked to the level of motivation to seek information, which is one of many factors that compete to influence gaze (other motivational, attentional, perceptual, and motor factors include: expectation of juice reward, general arousal, task engagement, time since visual stimulus onset, perceived stimulus intensity, recent history of saccades, etc.). Therefore, we found it more interpretable to express the neural-behavioral link in terms of modulations of neural activity.

#### Gaze shift analysis

We measured the timecourse of the change in neural activity aligned at the time of gaze shifts. We detected all gaze shifts onto or away from the CS/cue stimulus using the following procedure. Gaze shifts off of the CS were defined as milliseconds where (1) the gaze started in a stable ‘on’ state, by being ‘on’ for at least 150 consecutive ms, (2) starting at the current millisecond, the gaze transitioned to a stable ‘off’ state, by being ‘off’ for at least 100 consecutive ms, (3) the gaze was not merely hovering near the edge of the stimulus, indicated by the gaze 100 ms after the putative shift being at a location at least 5° away from the center of the stimulus, (4) the putative gaze shift was not influenced by changes in the eye tracker signal related to blinks, indicated by blinks not occurring in a 100 ms time window before the putative gaze shift. The results were not sensitive to the detailed settings of these parameters; similar results were obtained with other parameters. Gaze shifts onto the CS were defined in the analogous manner, except that to make our analysis of pre-shift activity more conservative we subtracted 40 ms from the gaze shift time, so that the gaze shift time more closely approximated the time when the eye movement began rather than the time when the eye entered the gaze window (Fig 5C, note that time = 0 is approximately when the gaze distance to the stimulus begins to change, rather than when the eye enters the gaze window).

In order to compare neural activity around the time of gaze shifts to neural activity in similar conditions where no gaze shift occurred, we matched each gaze shift with a set of control non-shift events. Gaze shifts were only included in the analysis if at least one control non-shift event was found. For each gaze shift, its control non-shift events were defined as all time points in the data that met the following criteria: (1) were recorded from the same neuron, (2) were on trials that presented the same CS, (3) were on trials that presented the same cue, if it was a task that used cues, (4) were at the same time point in the trial as the gaze shift, (5) had spent the previous 150 consecutive ms in the same initial gaze state as the gaze shift, (6) instead of shifting, the gaze continued to stay in that same gaze state for at least 100 consecutive ms. We then selected a subset of these non-shift events to use as controls in our analysis, in order to match the mean the gaze-to-stimulus distance before the non-shifts as closely as possible to the gaze-to-stimulus distance before the gaze shift. To do this, we calculated the difference between the gaze-to-stimulus distance 50 ms before each non-shift event vs. 50 ms before the gaze shift. We then selected non-shift events with a procedure that yielded differences that were small and centered around zero. Specifically, we first selected the non-shift event that had the smallest difference.

We then iteratively added non-shift events by repeatedly selecting the event whose difference met the following criteria: (A) had the opposite sign of the mean difference of the currently selected non-shift events, and (B) had the smallest magnitude of all remaining non-shift events meeting that criterion. For instance, if the first selected non-shift event occurred when gaze was slightly further away from the stimulus than before the gaze shift, the second selected non-shift event would occur when the gaze was slightly closer to the stimulus than before the gaze shift. If no remaining non-shift events met those criteria, no further non-shift events were selected. This procedure yielded 36306 gaze shifts, with a mean of 3.5 control non-shift events per gaze shift (standard deviation: 3.0, range: 1-31). There was a close match between the mean time course of the gaze-to-stimulus distances before gaze shifts vs. before the control non-shift events (Fig 5C).

Each neuron’s activity related to each of its individual gaze shifts was quantified as its normalized activity aligned on the gaze shift minus its mean normalized activity aligned on that gaze shift’s associated control non-shift events. The neuron’s overall activity related to gaze shifts was quantified by averaging its activity for all individual gaze shifts, separately for each type of gaze shift and each condition being analyzed (e.g. Fig 5D had two types of gaze shifts: on and off; and three conditions: gaze shifts when the trial’s reward outcome was uncertain, known to be reward, or known to be no reward). Neurons were only included in an analysis if they had at least one gaze shift of each type in each of the conditions being compared (e.g. Fig 5D only includes neurons that had at least one gaze shift in each of the 2 x 3 = 6 combinations of gaze shift type and condition). Activity around the time of the gaze shift was quantified in three time windows relative to the gaze shift: before the gaze shift (−0.4 to −0.1 s), during the gaze shift (−0.1 to +0.1 s), and after the gaze shift (+0.1 to +0.4 s). The first two time windows contain activity that cannot be explained as a result of visual feedback following the gaze shift, as the uncertainty-related neurons in these areas almost exclusively had visual response latencies > 0.1 s (e.g. Fig 1).

#### Gaze-modulation model analysis

Each neuron’s activity during the CS and cue periods in each task was fit with a computational model. The model’s parameters were divided into three groups (see Fig S8 for further explanation and illustrations). First, **β** parameters specifying the neuron’s *responses to CSs and cues*: its timecourse of activity in each task condition during moments when gaze is off the stimulus. This is similar to a traditional PSTH, and represents the timecourse that neural activity would have if there was no gaze-modulation. Second, **µ** and **σ** parameters specifying the *timecourse of gaze-modulation*: what times activity should be modulated relative to a gaze shift onto the stimulus, and how strongly gaze should be modulated at those times. Third, **w_gain_**and **w_offset_** parameters specifying the *effect of gaze-modulation*: whether gaze induces a multiplicative change in response strength (‘gain change’), an additive increase in firing rate (‘offset’), or both. Importantly, we validated the model by confirming that it accurately recovered the true gaze-modulation parameters when it was fitted to simulated datasets, including simulations where (A) gaze-modulation effects were similar to those in the real data, (B) gaze-modulations had the same timecourse as the real data but a variety of different magnitudes, (C) gaze-modulations had the same magnitudes as the real data but a variety of different timecourses, (D) ‘null hypothesis’ simulations in which gaze-modulations were absent (Fig S8).

The model was defined as follows. The **β** parameters specified the mean firing rate when gaze was off the stimulus location. A separate parameter, **β**(c,t), was defined for each trial condition *c* and each time bin *t* during the trial. Thus, similar to a conventional set of PSTHs, they specified the full timecourse of the neural response to each stimulus. Trial conditions were defined as combinations of CSs and cues. For instance, in the information task shown in Fig 2 there were 7 conditions (Info 100% CS with reward cue, Info 50% CS with reward cue, Info 50% CS with no-reward cue, Info 0% CS with no-reward cue, Noinfo 100% CS with non-informative cue, Noinfo 50% CS with non-informative cue, Noinfo 0% CS with non-informative cue). Time bins were defined as 50 ms bins spanning from 0.2 s after CS onset to 0.1 s after outcome delivery onset. Thus, a neuron recorded with a 3 sec total duration from CS onset to outcome delivery was fit with 7 conditions x 58 time bins = 406 distinct **β** parameters.

The **µ** and **σ** parameters specified the timecourse of gaze-modulation (Fig S8). Specifically, at each millisecond of each trial, a GazeMod variable specified the degree to which neural activity was modulated by gaze (0 <= GazeMod <= 1, 0 = no modulation, 1 = maximal modulation). GazeMod was computed by convolving a binary Gaze variable (1 = gaze is on the stimulus, 0 = gaze is off the stimulus) with a Gaussian kernel with mean = **µ** and standard deviation = **σ**. Thus, **µ** controlled the temporal offset between neural activity and gaze (whether neural activity is modulated before the gaze shifts onto the stimulus, or vice versa) and **σ** controlled the gradualness of gaze-modulation around a gaze shift (low **σ** = rapid onset of gaze-modulation, high **σ** = gradual onset). Thus, the timecourse of gaze-modulation around the time of a gaze shift took the form of a cumulative Gaussian function (Fig S8). The model’s gaze-modulation latency was calculated as the time relative to gaze onset when its timecourse of gaze-modulation reached 10% of its maximal value (i.e. when GazeMod = 0.1); other criteria produced the same main result (i.e. ACC before icbDS and Pal).

The **w_gain_** and **w_offset_**parameters specified the effect of gaze-modulation (Fig S8). Specifically, **w_gain_**caused gaze to multiplicatively scale firing rates, while **w_offset_**caused gaze to add or subtract from firing rates. Mathematically, the firing rate in a time bin with a given mean GazeMod was equal to: **β** x (1 + GazeMod x **w_gain_**) + GazeMod x **w_offset_**. Putting the parameters together, the neural firing rate in time bin *t* of trial *tr* that was in condition *c* was modeled as:

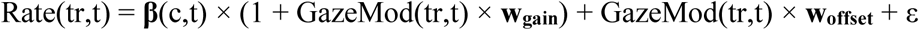

…where GazeMod(tr,t) is the mean GazeMod on trial tr in time bin t, and ε is a normally distributed noise term. Thus, if **w_gain_** = **w_offset_**= 0 then there was no gaze-modulation. If one of these parameters was non-zero, then gaze had a strictly multiplicative or additive effect on firing rate. Finally, if both parameters were non-zero, then the model multiplicatively increased the gain of neural responses relative to a baseline firing rate (e.g. during gaze, excitations above baseline are stronger excitations, and inhibitions below baseline are stronger inhibitions; Fig S8).

The model was fitted using the method of maximum likelihood, i.e. finding the parameters that maximize the probability that the model would produce the observed data. We did this by deriving the log likelihood function, its gradient, and its Hessian, and using them as input to the Trust Region Reflective Algorithm for function optimization to find the parameters that locally maximized the log likelihood (using its implementation in the ‘fmincon’ function in Matlab).

The parameters were constrained so that **µ** was in [-1,+1] and **σ** was in [0,1]; the other parameters were unconstrained. The initial parameter settings were **β**= 0, **µ** = 0, **σ** = 0.1, **w_gain_**= 0, **w_offset_** = 0; different initial parameter settings gave similar results. The optimization algorithm was allowed to continue for each session (here, defined as each set of data collected from a particular neuron with a particular task) for up to 100 function evaluations. The optimization algorithm successfully converged on 459/460 (99.8%) of sessions.

A neuron was considered to be significantly modulated by gaze if the log likelihood of the gaze model was significantly higher than expected by chance under the null hypothesis that there was no relationship between neural activity and gaze (p < 0.05, permutation test with 200 permutations). This was tested by comparing the log likelihood of the model fitted to the neuron’s true data to the distribution of log likelihoods of the model fitted to shuffled datasets in which the neural data was exactly the same as the original dataset, but the gaze data were randomly shuffled among trials that shared the same task condition. For instance, suppose the 75% reward CS was presented on trials 5, 12, 14, 22, and 24. The neural data on those trials would be kept the same, but the gaze data would be shuffled so that the gaze data from trial 5 might now be assigned to trial 22; the gaze data from trial 12 might now be assigned to trial 14; etc. Thus, shuffling destroyed any trial-to-trial relationships between neural activity and gaze, while leaving the neural and gaze data intact *per se*. The neuron was classified as significantly gaze-modulated if the log likelihood of the model fit to the true dataset was greater than the log likelihoods of the model fits to at least 191/200 of the permuted datasets. Note that this is a one-tailed test, because the shuffling cannot (in expectation) improve the quality of the fit. That is, if there was a true neural-gaze relationship in the original data then shuffling should worsen the fit, while if there was no such relationship then shuffling should not affect the quality of the fit; in neither case could shuffling improve the fit.

The latency of gaze-modulation was compared across areas using two methods which gave consistent results. First, we calculated each area’s mean timecourse of gaze-modulation by averaging the fitted timecourses for each neuron in that area with significant gaze-modulation (Fig 6F) and using this to calculate the area’s latency in the same method used for single neuron timecourses. We then calculated whether the per-area latencies were significantly different (p < 0.05, permutation test conducted by comparing the difference between the area latencies with the distribution of such differences computed from 2000 permuted datasets in which the assignment of neurons to the two areas was randomly shuffled). Second, we directly compared the distribution of fitted latencies from all single neurons with significant gaze-modulation in the two areas (Fig 6F inset, rank-sum test, p < 0.05).

#### Quantification of information seeking behavior during inactivations

Response times (RTs) were computed online and used to determine when the CS was replaced by the cue, defined as the time between the ‘go’ signal (i.e. the trial start cue’s disappearance) and the gaze entering the response window around the CS. To improve the accuracy of RT measurements for our offline analysis, RTs were recomputed offline using response windows that were corrected for session-to-session variability in eye tracker settings using the procedure described above (i.e. centering the window on the observed peak gaze location separately for each CS location and each session). We then analyzed the RTs from all correctly performed trials in which the animal made a response and there was at least rough agreement between the online and offline RTs (i.e. within 0.2 s of each other). These criteria were met by nearly all correctly performed trials (n=17968/18035; 99.6%). We then quantified the information seeking bias using an Infobias Index based on the mean RTs for the Info and Noinfo CSs:

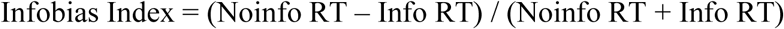

…which was computed separately for each session, and separately for each of the 2 x 2 combinations of time in session (pre- vs post-injection) and CS location relative to injection site (contralateral vs ipsilateral). We then derived two additional measures. For session and each CS location, we defined the change in Infobias index as the difference between post-injection and pre-injection Infobias indexes. We defined the change in Infobias laterality as the difference between the changes in Infobias Index for the contralateral and ipsilateral sides.

To test whether icbDS and Pal inactivations affected information seeking behavior (Fig 7C), we computed the mean change in Infobias index and tested whether it was significantly different from zero (p < 0.05, permutation test conducted by comparing the true mean change in Infobias Index to the distribution of mean changes in Infobias Index computed on 20000000 permuted datasets in which pre- and post-injection data were shuffled with each other). For pooled data from all inactivation sessions and for control sessions (Fig 7C) we used the same procedure, except to be conservative we included an additional correction for any potential main effects of animal, by using weighted means such that each animal’s inactivation and control data were weighted by the number of inactivation sessions that animal contributed to the dataset. The same key results were obtained in uncorrected data (significant change in contralateral Infobias Index during inactivation sessions, p = 0.0015, signed-rank test; no significant change in Control sessions, p = 0.8786, signed-rank test; changes in inactivation sessions significantly different from changes in control sessions, p = 0.0297, rank-sum test).

To test how inactivations interfered with information seeking behavior, we analyzed RTs separately for Info and Noinfo CSs. RTs were normalized by z-scoring all RTs separately for each session and CS location. Then for each area and each CS type we tested whether there was a significant difference between the pre- vs. post-injection RT distributions for contralateral CSs (p < 0.05, ranked-sum test). We further tested if the changes in RTs were significantly different for the two CS types (p < 0.05, interaction term from a 2-way ANOVA using the factors CS type (Info or Noinfo) and epoch in session (pre- or post-injection)). Finally, to directly compare the effects of injections in the two areas, we first computed each area’s overall change in mean normalized RT averaged across the Info and Noinfo CSs (computed as ΔRT_overall_ = 0.5*(mean(Info after) – mean(Info before)) + 0.5*(mean(Noinfo after) – mean(Noinfo before))) and tested whether this was significantly different between the two areas (permutation test, conducted by comparing the difference between the ΔRT_overall_ of the two areas and comparing it to the distribution of differences computed on 2000 permuted datasets in which each session’s pre- and post-injection data were shuffled with each other). We then computed the component of this overall change in RTs that occurred in the hypothesized manner, i.e. icbDS injection slowing RTs to the Info CS or Pal injection speeding RTs to the Noinfo CS, as ΔR_Thypothesized_ = (mean(icbDS Info after) – mean(icbDS Info before)) – (mean(Pal Noinfo after) – mean(Pal Noinfo before)). We also computed the component that occurred in the orthogonal manner to our hypothesis, i.e. icbDS injection slowing RTs to the Noinfo CS or Pal injection speeding RTs to the Info CS, as ΔR_Torthogonal_ = (mean(icbDS Noinfo after) – mean(icbDS Noinfo before)) – (mean(Pal Info after) – mean(Pal Info before)). We tested these components for significance using the same type of permutation test (comparing them to the distribution of the components calculated from 2000 permuted datasets in which each session’s pre- and post-injection data were shuffled with each other).

To test how inactivations might be expected to interfere with information seeking given the broad strokes of the cortico-BG circuit, we implemented a simple computational model of the network and used it to generate a simulated RT for each trial in our dataset (Fig 7E). We then analyzed the simulated RTs in the same manner as the real RTs. Note that this model is intended as a simple test of whether information seeking behavior and its perturbation by icbDS and Pal inactivations are consistent with a straightforward implementation of the typical cortico-BG information signals we observed and the signs of excitatory/inhibitory connections between areas. This model is not intended to emulate the detailed dynamics of neural activity within or between areas or to emulate the detailed circuitry of visual processing and saccade generation. In this model, each neural population in each hemisphere was represented by a single simulated neuron whose firing rate *r* on each trial was:

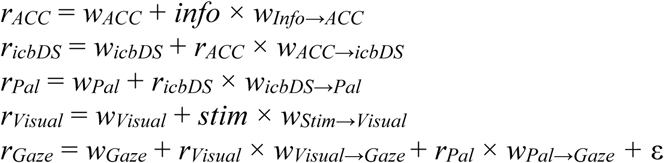

…where for each area X the variable *w_X_* represents the area’s baseline firing rate and *w_Y→X_* represents the weight of incoming input from area Y to area X in the same hemisphere, *info* = 1 for trials with an Info CS and 0 for trials with a Noinfo CS, *stim* = 1 for trials with a contralateral CS and 0 for trials with an ipsilateral CS, and ε is Gaussian noise drawn independently on each trial with mean = 0 and standard deviation = 30. Here, Visual represents a population of neurons responsive to visual input for the purpose of directing eye movements (e.g. visually responsive neurons in the frontal eye field, lateral intraparietal sulcus, and/or the superior colliculus ^95–97^), and Gaze represents a population of neurons that directly control eye movements (e.g. saccade-related neurons in the deep layers of superior colliculus ^95^). Note that the connections in this model are not meant to imply that effects are necessarily mediated in the brain by direct monosynaptic connections; the connections correspond to the net effect of both direct and indirect influences of activity in one area on another area. The baseline and input weights of each cortico-BG area were selected to approximately match the typical firing rates of information-related neurons in that area in the moment before cue delivery in the information task, and to be consistent with the excitatory/inhibitory nature of each connection in the cortico-BG network, as follows. Baseline rates were: ACC = 2, icbDS = 0, Pal = 50, Visual = 0, Gaze = 0. Input weights were: Info→ACC = +20, Stim→Visual = +100, ACC→icBDS = +1, icbDS→Pal = −2, Visual→Gaze = +1, Pal→Gaze = −1. Finally, the simulated RTs for each trial were generated by taking the firing rates from the Gaze neuron in the contralateral hemisphere to the CS and mapping them onto the distribution of RTs in the real data (so that the trial with the K-th highest firing rate from the Gaze neuron was given the K-th fastest RT observed in control sessions in the real data). These simulated RTs were then analyzed in the same manner as the real data. Inactivations were simulated by multiplying the firing rate of the inactivated area (icbDS or Pal) by a scaling factor of 0.7, representing a 30% reduction in firing rate; other scaling factors produced qualitatively similar results (i.e. icbDS inactivations predominantly slowing Info CS RTs and Pal inactivations predominantly speeding Noinfo CS RTs).

### Anatomy and tracer injections

#### Surgery, tissue preparation, and microscopy

Experiments were carried out on separate animals from those used for electrophysiology. Some of these injections were analyzed for other anatomical studies of the striatal-pallidal network (e.g. refs ^26, 98^). Procedures were conducted in accordance with the Institute of Laboratory Animal Resources Guide for the Care and Use of Laboratory Animals and approved by the University Committee on Animal Resources at the University of Rochester. Adult male macaque monkeys (data from 1 *Macaca nemestrina,* 2 *Macaca fascicularis* shown here) were tranquilized by intramuscular injection of ketamine (10 mg/kg). For a subset of animals, MRI (3 T) T1 or T2 turbo spin echo scans (0.5 mm × 0.5 mm × 1.42 mm) were obtained before surgery. For the others, serial electrode penetrations were made to locate the anterior commissure, as described previously ^99^. These images and recordings were used to calculate the anterior/posterior, dorsal/ventral, and medial/lateral coordinates for each tracer injection from stereotaxic zero.

Animals received ketamine 10 mg/kg, diazepam 0.25 mg/kg, and atropine 0.04 mg/kg IM in the cage. A surgical plane of anesthesia was maintained by either intravenous injections of pentobarbital (for recordings, initial dose 20 mg/kg, i.v., and maintained as needed) or via 1-3% isoflurane in 100% oxygen via vaporizer. Temperature, heart rate, and respiration were monitored throughout the surgery. Monkeys were placed in a Kopf stereotaxic, a midline scalp incision was made, and the muscle and fascia were displaced laterally to expose the skull. A craniotomy (∼2-3 cm^2^) was made over the region of interest and small dural incisions were made only at injection sites.

Monkeys received an injection of one or more of the following anterograde/bidirectional tracers: lucifer yellow, fluororuby, or fluorescein conjugated to dextran amine (LY, FR, or FS; 40–50 nl, 10% in 0.1M phosphate buffer (PB), pH 7.4; Invitrogen); wheat germ agglutinin conjugated to horseradish peroxidase (WGA; 40–50 nl, 4% in distilled water; Sigma, St. Louis, MO); *Phaseolus vulgaris-*leucoagglutinin (PHA-L; 50 nl, 2.5%; Vector Laboratories); or tritiated amino acids (AA, 100 nl, 1:1 solution of [^3^H] leucine and [^3^H]-proline in dH2O, 200 mCi/ml, NEN). These tracers do not cross-react with one another, and thus an individual animal can serve in multiple experiments, reducing the total number of animals needed. Tracers were pressure-injected over 10 min using a 0.5-µl Hamilton syringe. Following each injection, the syringe remained in situ for 20–30 min. Twelve to 14 days after surgery, monkeys were again deeply anesthetized and perfused with saline followed by a 4% paraformaldehyde/1.5% sucrose solution in 0.1 M PB, pH 7.4. Brains were postfixed overnight and cryoprotected in increasing gradients of sucrose (10, 20, and 30%). Serial sections of 50 µm were cut on a freezing microtome into 0.1 M PB or cryoprotectant solution, as described previously ^100^.

One in eight sections was processed free-floating for immunocytochemistry to visualize the tracers. Tissue was incubated in primary anti-LY (1:3000 dilution; Invitrogen) or anti-FR (1:1000; Invitrogen in 10% NGS and 0.3% Triton X-100 (Sigma-Aldrich) in PB for four nights at 4 °C. Following extensive rinsing, the tissue was incubated for 40 min in biotinylated secondary anti-rabbit antibody made in goat (1:200; Vector BA-1000) followed by incubation with the avidin–biotin complex solution (Vectastain ABC kit, Vector Laboratories). Immunoreactivity was visualized using standard DAB procedures. Staining was intensified by incubating the tissue for 5–15 s in a solution of 0.05% 3,3′-diaminobenzidine tetra-hydrochloride, 0.025% cobalt chloride, 0.02% nickel ammonium sulfate, and 0.01% H2O2. Sections were mounted onto gel-coated slides, dehydrated, defatted in xylene, and cover-slipped with Permount.

Using darkfield light microscopy, brain sections, injection sites, and dense pallidal terminal fields were outlined under a 1.6, 4.0, or 10× objective using a Leitz or Leica microscope with Neurolucida software (MBF Bioscience). Terminal fields were considered dense when they could be visualized at a low objective (1.6×) ^100^. Retrogradely labeled input cells were identified under brightfield microscopy (20×). StereoInvestigator software (Micro-BrightField) was used to stereologically count labeled cells with an even sampling (64%). On the additional three cases (45LY, 113FS, 40LY), cells were charted in representative select frontal sections using the same parameters. Other sections were visually inspected for labeling.

### Reporting

The datasets generated during and/or analyzed during the current study and custom code for mathematical algorithms are available from the corresponding author on reasonable request.

## AUTHOR CONTRIBUTIONS

J.K.W. contributed to conceptualization and study design, led data collection for electrophysiology and pharmacology, and contributed to data analysis; E.S.B.-M. contributed to study design and led data analysis and writing the manuscript; S.R.H. performed the anatomical analysis; K.Z. contributed to data collection for electrophysiology and pharmacology and data analysis; J.P. contributed to data collection for electrophysiology; S.N.H. provided anatomical data and supported the anatomical analysis; I.M. contributed to conceptualization, study design, and data collection for electrophysiology, and provided overall project supervision and funding acquisition; and all authors contributed to writing the manuscript.

## ACKNOWLEDGEMENTS

We are grateful to Dr. Noah Ledbetter for assisting in data acquisition, Dr. Timothy Holy for helpful discussions, and Ms. Kim Kocher for fantastic animal care and training. This work was supported by the National Institute of Mental Health under award number R01MH110594 and R01MH116937, the Edward Mallinckrodt, Jr. Foundation, and the McDonnell Center for Systems Neuroscience.

## Supplementary Information

**Figure S1.**
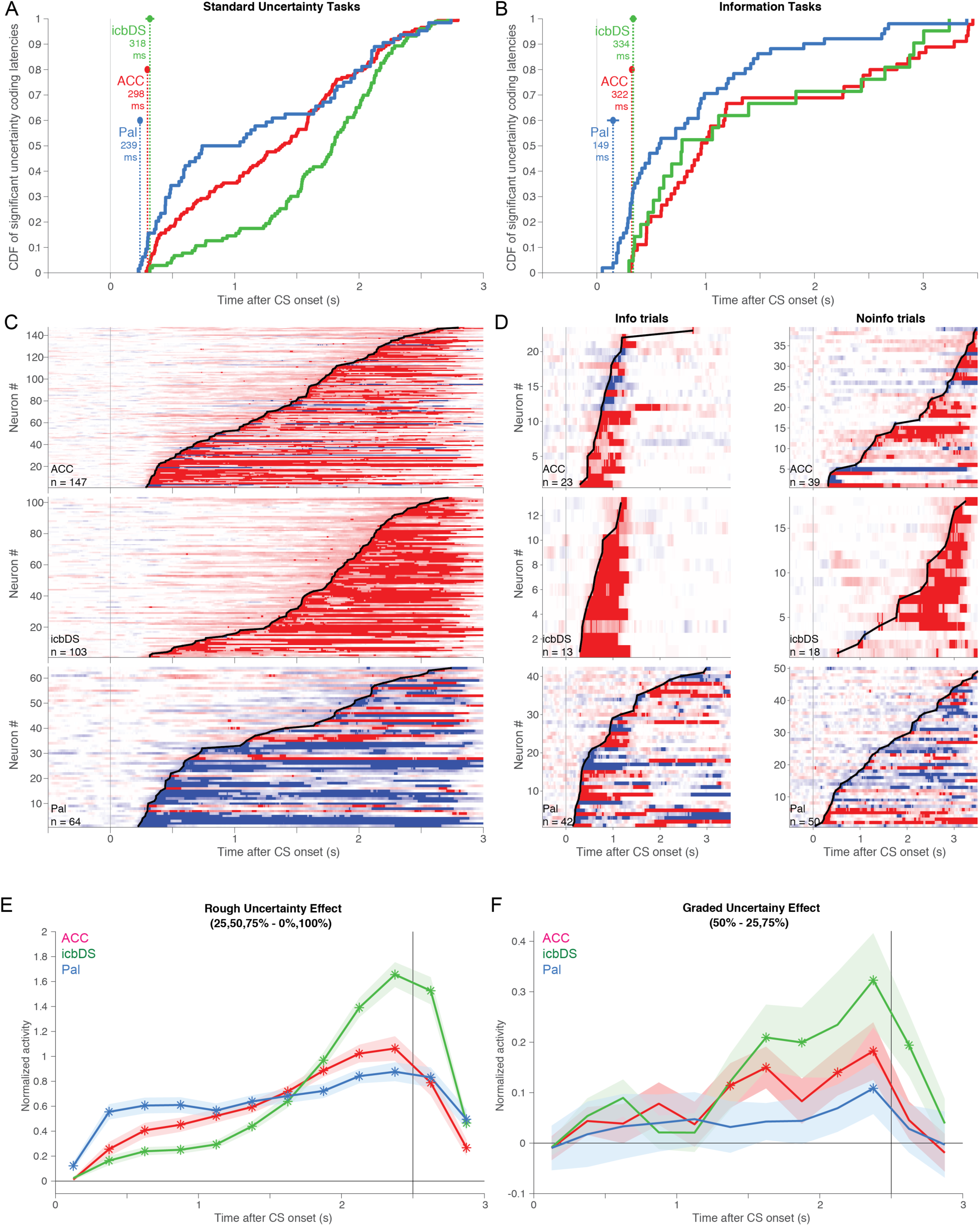
Detailed properties of uncertainty signals: latency, sign, and emergence of rough vs. graded coding. (**A**) Cumulative distribution of latencies for all single neurons recorded during standard uncertainty tasks that had detected latencies of uncertainty coding. Circles, dashed vertical lines, and text indicate the estimated population latency of uncertainty coding in each area (error bars are ± 1 bootstrap SE). Uncertainty signals emerge first in Pal, followed by ACC and icbDS at similar latencies. The population latency is significantly shorter in Pal than both ACC and icbDS (p = 0.005 and p = 0.001, permutation tests) and not significantly different between ACC and icbDS (p = 0.35). (**B**) Same, for neurons recorded during information tasks. Latencies were calculated separately for Info and Noinfo trials, and each neuron’s overall latency of uncertainty coding was defined as the shorter of the two. Note that the power to detect latency differences between areas is considerably smaller in this dataset due to the smaller number of neurons. Nonetheless, the data shows a similar pattern: the population latency is significantly or trending to be shorter in Pal than both ACC and icbDS (p = 0.018 and p = 0.15) and not significantly different between ACC and icbDS (p = 0.63). (**C**) Heat map of uncertainty coding over time for the neurons in (A). Top: ACC; middle: icbDS; bottom: Pal. Each row is a neuron. Black lines indicate each neuron’s detected latency of uncertainty coding. Color indicates the neuron’s uncertainty coding at each time during the task (red = more active for uncertain than certain CSs, blue = less active for uncertain than certain CSs, white = no uncertainty coding; light colors = non-significant uncertainty coding, dark colors = significant uncertainty coding). Uncertainty coding is quantified based on the ROC area for using the neuron’s activity to discriminate between uncertain vs certain CSs. For this latency analysis we conservatively calculate the uncertainty coding at each time point as the least extreme of two ROC areas: one for discriminating all uncertain CSs from the 100% reward CS, and the other for discriminating all uncertain CSs from the 0% reward CS. Uncertainty coding is predominantly excitatory in ACC, almost exclusively excitatory in icbDS, and predominantly inhibitory in Pal. (**D**) Same, for neurons recorded during information tasks. Data is shown separately for Info and Noinfo trials (left and right columns). For this analysis we calculate the uncertainty coding at each point as the ROC area for discriminating uncertain from certain reward trials. Note that the “n” is different because neurons with detectable latencies on Info trials did not always have detectable latencies on Noinfo trials, and vice versa. (**E**) Population average rough uncertainty signal for each area during standard uncertainty tasks, defined as the difference in mean normalized activity between uncertain CSs (25%,50%, and 75%) and certain CSs (0% and 100%). The shaded area is ± 1 SE. Asterisks indicate significant differences from 0 (signed-rank test, p < 0.05). Consistent with the latency analysis, the rough uncertainty signal reaches significance first in Pal and later in ACC and icbDS. The population average rough uncertainty signal was significantly higher in Pal than either ACC or icbDS in all time bins within the first 1 sec after CS onset (all p < 0.05, rank-sum tests), and conversely, was significantly higher in both ACC and icbDS than Pal in all time bins in the last 0.75 sec before outcome delivery (all p < 0.05, rank-sum tests). This indicates that the shorter latency of uncertainty signals in Pal was due to Pal having a different timecourse of uncertainty coding, consisting of faster but ultimately weaker signals (i.e., not due to Pal simply having overall stronger signals than other areas, or due to Pal having the same average signal strength as other areas but concentrated into small fraction of its uncertainty coding neurons that are endowed with especially strong signals that allow their latencies to be detected earlier). (**F**) Population average graded uncertainty signal in each area, defined as the difference in mean normalized activity between the maximally uncertain CS (50%) and the moderately uncertain CSs (25% and 75%). There is a trend for a graded uncertainty coding at all time points in all areas, but it grows and reaches significance first in ACC and icbDS, and later in Pal. Thus, while rough uncertainty coding emerged first in Pal, graded uncertainty coding became significantly prevalent first in ACC and icbDS.

**Figure S2.**
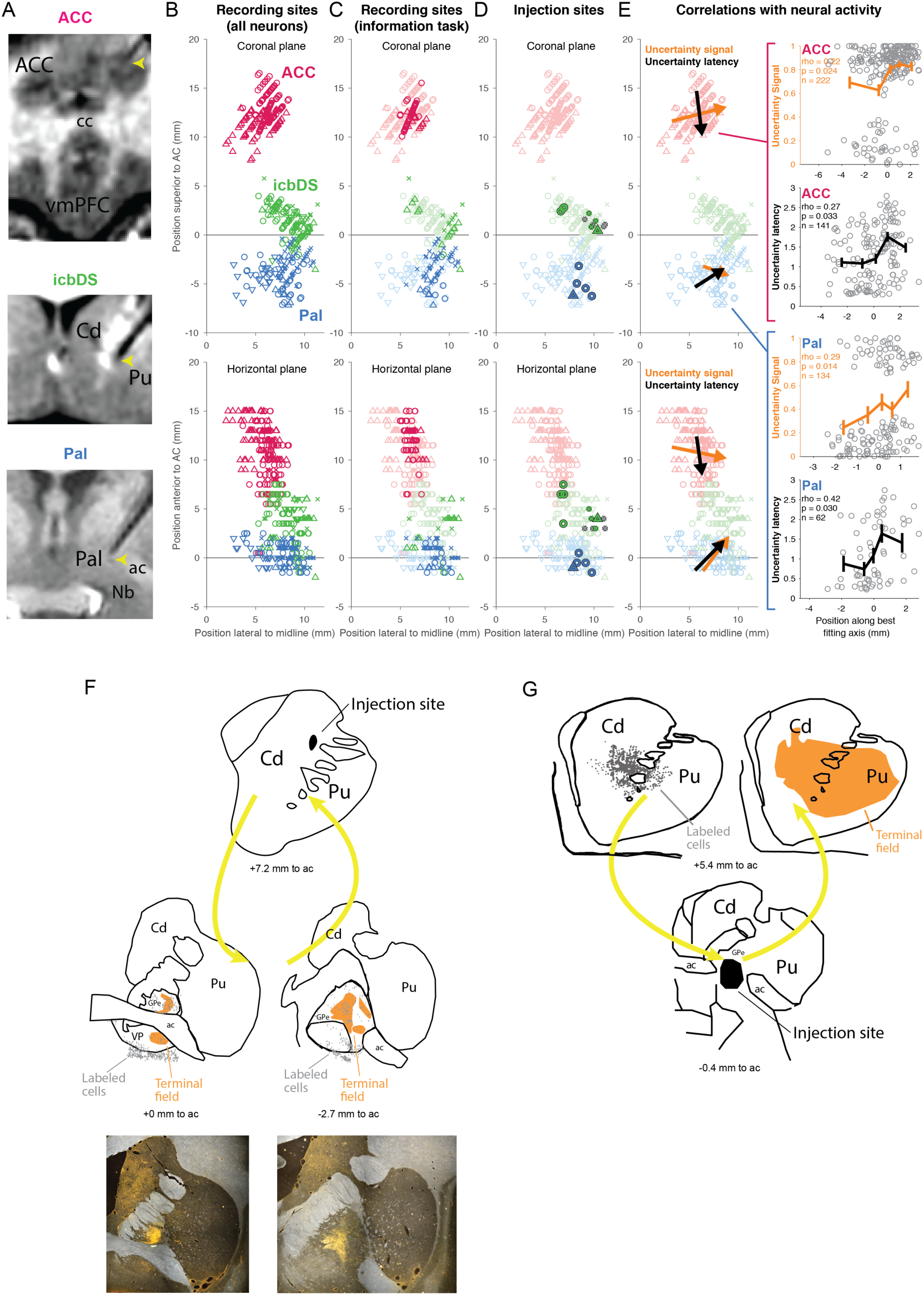
Anatomical tracing, reconstruction of recording sites, and correlation with neural responses. (**A**) **MRI verification of recording sites.** MRIs were taken immediately after recording uncertainty coding neurons with the electrode still in place. Shown are coronal views in which the electrode track is visible as a black ‘shadow’ on the MRI; the yellow arrow indicates the location of the electrode tip. Top: Recording site in ACC, at a location symmetrical to the ACC in the opposite hemisphere. Middle: Recording site in icbDS, at a location intermediate between the body of the caudate and putamen. Bottom: Recording site in Pal, with the tip adjacent to the ventral boundary of the anterior commissure. Abbreviations: cc, corpus callosum; vmPFC, ventromedial prefrontal cortex; Cd, caudate; Pu, putamen; ac, anterior commissure; Nb, nucleus basalis. (**B**) **Anatomical locations of recording sites.** Reconstructed 3D coordinates of each neuron in the dataset, shown for all areas (indicated by colors) and all animals (indicated by symbols: animals B, R, W, and Z are circles, crosses, downward triangles, and upward triangles). Coordinates are relative to the midline, superior tip of the anterior commissure (AC). *Top* shows coordinates in the coronal plane. *Bottom* shows coordinates in the horizontal plane. The three areas where uncertainty-responsive neurons were found were clearly anatomically distinct from each other, and were located in similar, overlapping locations in all animals. (**C**) **Anatomical locations of recording sites for each task type.** Light colors indicate neurons recorded in standard uncertainty tasks. Dark colors indicate neurons recorded in information tasks. There is considerable overlap between recording sites for the two task types. (**D**) **Anatomical locations of injection sites.** Same as (B) with an overlay showing the reconstructed coordinates of the injection sites. Colors indicate injection type (green, icbDS muscimol; blue, Pal muscimol; gray, icbDS saline). The injection sites for icbDS and Pal were clearly distinct from each other and overlapped the locations of uncertainty-responsive neurons in their respective areas. The injection sites for saline overlapped with the injection sites for muscimol in the same area. (**E**) **Anatomical relationships to neural responses.** We plot all significant correlations between neuron locations and response properties (p < 0.05, permutation test). Each relationship is indicated by an arrow in the left column (viewed in the coronal plane (top) and in the horizontal plane (bottom)), and a corresponding scatterplot on the right column. ***Left:*** Colored arrows in each area indicate relationships between neuron location and one of four response properties: uncertainty signal (orange), uncertainty signal latency (black), fitted gaze-modulation latency (none significant), fitted gaze-modulation gain change expressed in log_2_ units (none significant). Each arrow passes through the center of mass of the locations of the neurons recorded in that area, and is directed along the vector in 3D space that best predicts the response property. Specifically, we fitted each response property with linear regression as a function of neural coordinates in 3D space (ResponseProperty = β_0_ + β_1_*Lateral + β_2_*Anterior + β_3_*Superior+ ε). We then pointed the arrow in the direction of the unit vector v = <β_1_, β_2_, β_3_> / |<β_1_, β_2_, β_3_>|, and scaled the arrow’s length by the standard deviation of the Euclidean distance between the neuron locations and their center of mass. P-values and significance were assessed with a permutation test (by comparing the log likelihood of the fit to the true dataset with the distribution of log likelihoods of fits to 20000 permuted datasets generated by shuffling the relationship between the 3D coordinates of neurons and the responses of those neurons). ***Right:*** Each plot shows a single significant relationship. Each dot is a single neuron’s response property. The x-axis indicates each neuron’s location relative to its area’s center of mass, projected onto the best-predictive unit vector for that area. The y-axis indicates that neuron’s response property in the task. Lines with five data points indicate mean ± SE of the response property for the neurons that were located in each quintile of the neuron locations along the best-fit line. Text indicates rank correlation, the p-value of the linear regression that determined the vector (permutation test described above), and the number of neurons for the comparison. Uncertainty latency relationships have fewer neurons than other response properties because not all neurons had detectable latencies (Fig S1). In ACC, there were tendencies for superior-lateral neurons to have stronger uncertainty signals (orange, p = 0.024) and for posterior-inferior neurons to have longer latencies of uncertainty signals (black, p = 0.033). In Pal, there were tendencies for anterior-lateral neurons to have stronger uncertainty signals (orange, p = 0.014) and longer latencies of uncertainty signals (black, p = 0.030). Certain neuronal signals and functions of the ACC may be organized in a dorsal-ventral manner (e.g. based on their location across the dorsal and ventral banks of the cingulate sulcus). Therefore, we explicitly tested whether the coding properties of ACC neurons varied significantly with their position along the dorsal-ventral axis. Consistent with our analysis in Fig S2E, there was a significant correlation such that uncertainty coding neurons located more dorsally had more positive uncertainty signals (n = 222, rho = +0.16, p = 0.017). There were no significant correlations with other coding properties (uncertainty coding latency in neurons with detectable latencies, n = 141, rho = 0.02, p = 0.795; gaze-modulation latency, n = 221, rho = - 0.02, p = 0.729; gaze-modulation gain factor, n = 221, rho = 0.00, p = 0.978). (**F**) **Additional tracer injection into icbDS confirms bidirectional icbDS-Pal projections.** Top: the tracer fluororuby was injected into the internal capsule-bordering region of the dorsal striatum (black area, top). Retrogradely labeled cell bodies (gray dots, bottom) and anterogradely labeled terminal fields (orange shaded areas, bottom) were found in Pal, including both the ventral pallidum (below the anterior commissure) and the anterior globus pallidus external segment (above the anterior commissure). Text indicates the anterior-posterior position relative to the midline anterior commissure. Bottom: photomicrographs show anterogradely labeled terminal fields in Pal. (**G**) **Tracer injection into Pal confirms bidirectional icbDS-Pal projections.** The tracer Lucifer Yellow was injected into the border of the globus pallidus external segment and ventral pallidum (black area, bottom). Retrogradely labeled cell bodies (gray dots, top left) and anterogradely labeled terminal fields (orange shaded areas, right) were found in the dorsal striatum, including the internal capsule-bordering regions of the caudate and putamen. Text indicates the anterior-posterior position relative to the midline anterior commissure. Abbreviations: Cd, caudate nucleus; Pu, putamen; ac, anterior commissure; GPe, globus pallidus external segment; VP, ventral pallidum.

**Figure S3.**
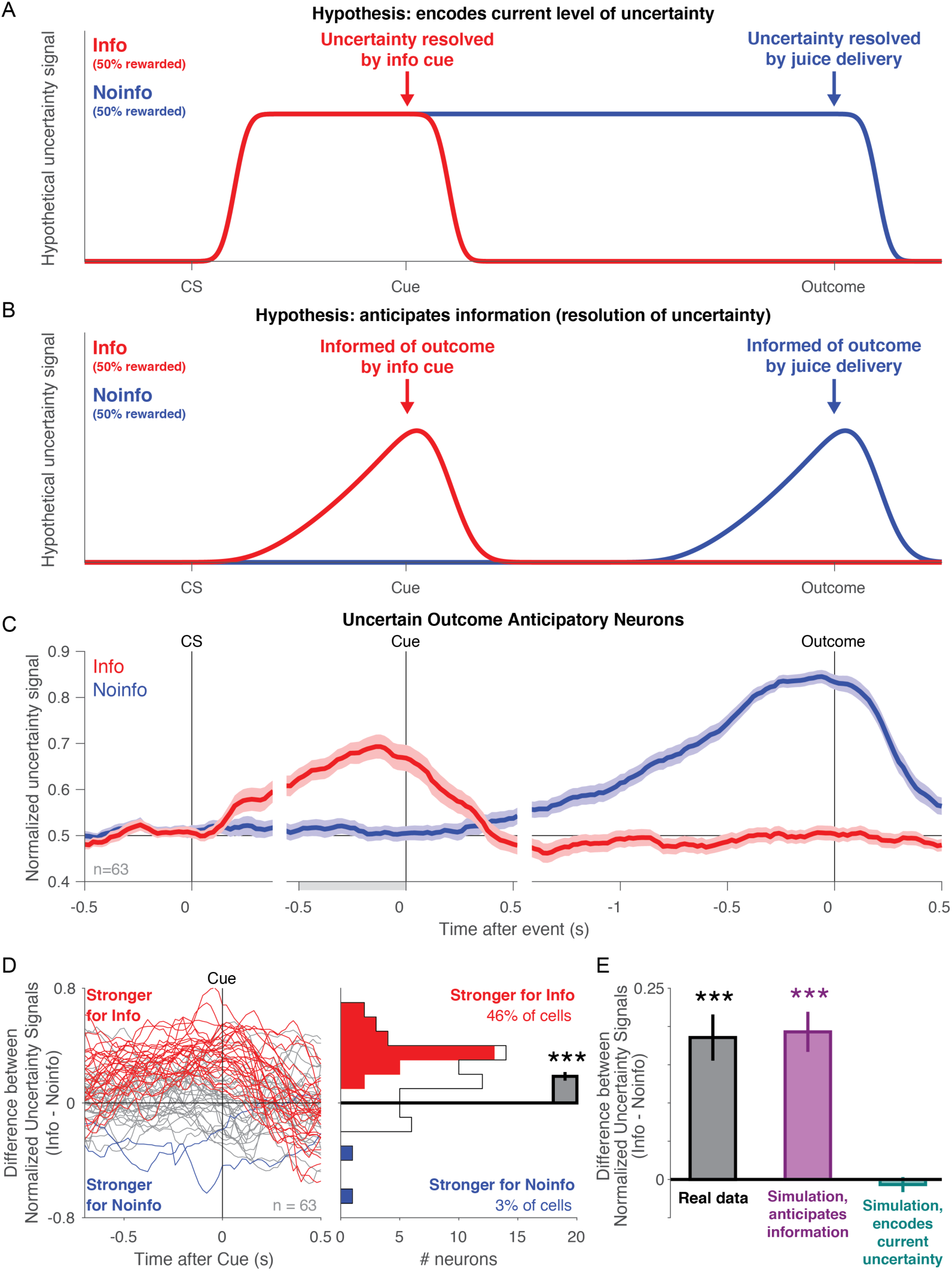
Information-anticipatory activity cannot be explained by encoding of the current level of uncertainty. (**A**) Hypothetical timecourse of uncertainty signals for a neuron encoding the animal’s current level of uncertainty about the size of the trial’s reward outcome. The neuron would respond equally to both the Info 50% reward CS and Noinfo 50% reward CS, because the reward is equally uncertain in both situations (i.e. they both predict identical reward distributions: 50% chance of reward, 50% chance of no reward). The only difference between the conditions would occur after the CS period ended, based on the time the neuron terminated its response. On Info trials, the neuron would stop responding after the Info cue (because the cue informs the animal of the upcoming reward, and hence fully resolves the uncertainty), but on Noinfo trials, the neuron would only stop responding after outcome delivery (because the cue does not predict the outcome, and hence the uncertainty is only resolved after outcome delivery). (**B**) Hypothetical timecourse of uncertainty signals for a neuron that anticipates the time of receiving information to resolve reward uncertainty. Same as Fig 2B. The neuron should clearly differentiate between Info and Noinfo 50% reward CSs during the CS period, with a stronger uncertainty signal for the Info 50% reward CS. (**C**) Observed timecourse of normalized uncertainty signals in the uncertainty coding neurons that had significant Uncertain Outcome Anticipation Indexes. Same as Fig 3C. Importantly, these neurons were not selected in any manner based on their activity on Info trials or their activity during the CS period; they were only selected based on their outcome-anticipatory activity on Noinfo trials. Nonetheless, these neurons clearly differentiated between Info and Noinfo trials during the CS period, with stronger uncertainty signals during the Info CS. (**D**) Difference between timecourses of normalized uncertainty signals on Info trials vs. Noinfo trials (i.e., red curve in panel C – blue curve in panel C) for each of these neurons. Red, blue, and gray lines indicate neurons with significantly positive, negative, or non-significant differences (p < 0.05, permutation test on the activity in a 500 ms window before cue onset). A large fraction of the neurons have higher normalized uncertainty signals for Info, in a manner that ramps up to the time of the cue. (**E**) Quantification of the mean difference between normalized uncertainty signals in the 500 ms before cue onset. Red, blue, and white cells correspond to the colors in panel D. Nearly half of cells have significant positive differences (46% of cells, higher than expected by chance, p < 0.001, binomial test). Only 2 neurons have significant negative differences (3% of cells, not significantly different from chance, p = 0.47). The average difference over all neurons is +0.186, which is significantly greater than 0 (gray bar, p < 0.001, signed-rank test). (**F**) Direct test of whether neural activity is better explained by anticipation of information or encoding of the current level of uncertainty. Bars indicate the mean ± SE of the mean differences plotted in panels D,E. The gray bar represents this value computed using the real data (same as the gray bar in panel E). The other bars represent this value computed using simulated datasets representing the two competing hypotheses. The simulated datasets for each hypothesis were generated by starting with the real dataset, finding pairs of task conditions that *should* have equivalent activity according to that hypothesis, and by drawing new firing rates for each trial in the first condition from the distribution of firing rates in the second condition, to force those conditions to be equivalent in the simulated data. This procedure was done separately for each neuron. For the simulations representing ‘anticipates information’, activity during the CS period on Noinfo trials should be equal to activity on Info certain trials (i.e. little or no activity). Hence we set the rates on all Noinfo trials to be drawn from the distribution of rates on Info certain trials. For the simulation representing ‘encodes current uncertainty’, Info and Noinfo trials should have equivalent activity during the CS period. Hence we set the rates on Info certain trials to be drawn from the distribution of rates on Noinfo certain trials, and set the rates of Info uncertain trials to be drawn from the distribution of rates on Noinfo uncertain trials. The results were clear: the mean difference in the real data was highly similar to the mean difference in the simulated ‘anticipates information’ dataset (purple bar), and clearly distinct from the simulated ‘encodes current uncertainty’ dataset (blue bar).

**Figure S4.**
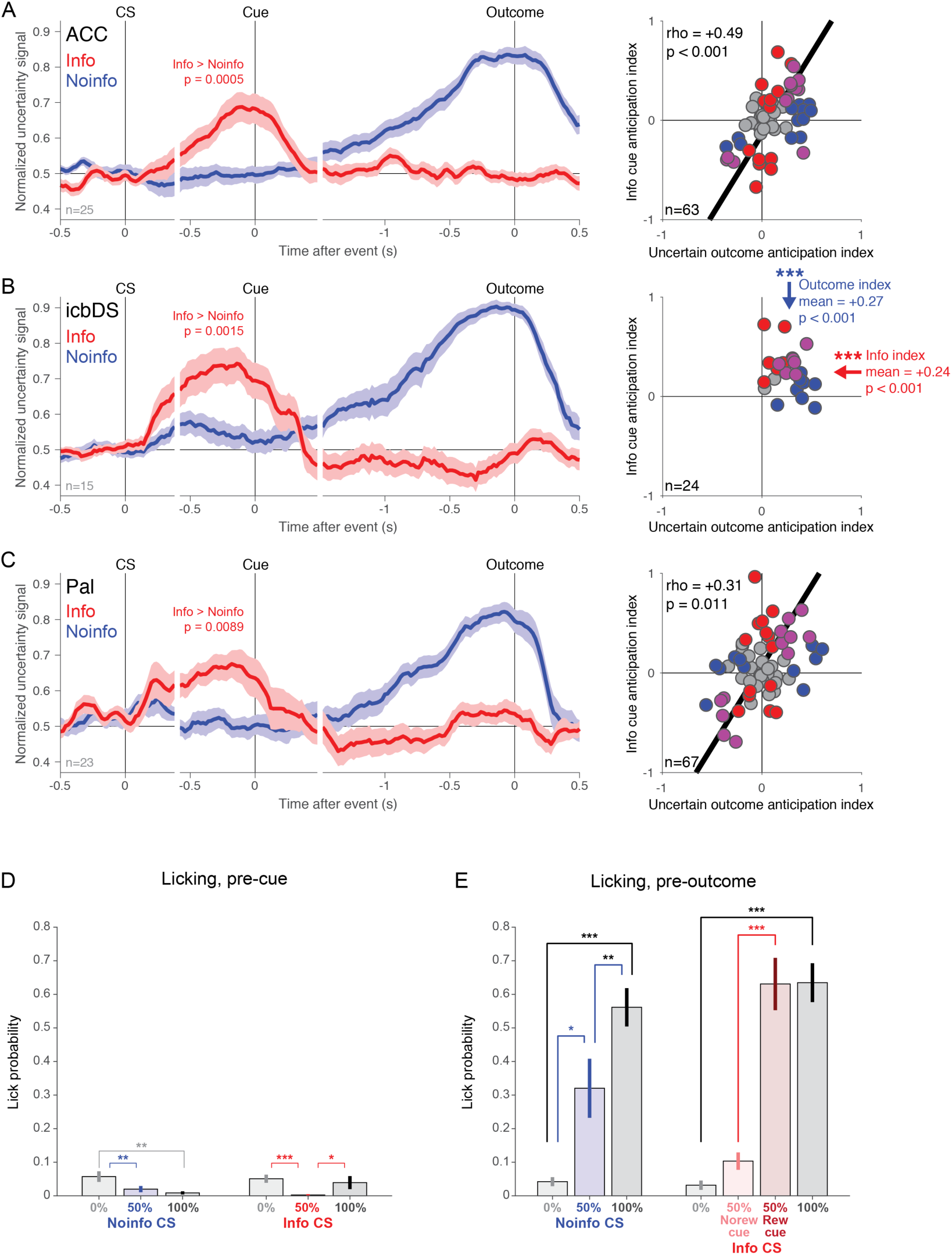
Information-anticipatory activity in all areas, and absence of information-anticipatory licking behavior. (**A,B,C**) Test for information-anticipatory activity in ACC (A), icbDS (B), and Pal (C). *Left:* same format as Fig 3C for each area. The average uncertainty signal of neurons that had a significant ramping uncertainty activity measured only using Noinfo trials (i.e. cells with a significant Uncertain Outcome Anticipation Index, p < 0.05, permutation test). Note that these neurons are selected solely based on their activity anticipating the outcome on Noinfo trials. Even so, in all areas these populations show a strong uncertainty signal in the same direction anticipating the cue on Info trials. Thus, these neural populations had information-anticipatory activity resembling the theoretical pattern in Fig 2B. *Right:* same format as Fig 3D for each area. Best-fit lines from type 2 regression are plotted for all areas with significant correlations (p < 0.05); arrows indicate significant mean indexes different from 0 (p < 0.05). All areas show coding indexes consistent with information-anticipatory activity, though in different manners due to the different signs of neural coding in icbDS vs ACC and Pal. icbDS shows an especially strong pattern of information-anticipatory activity. icbDS neurons only encode uncertainty with a positive sign, and nearly all neurons are in the upper right quadrant, indicating positive anticipation of both informative cues and uncertain outcomes (mean Informative Cue Anticipation Index = +0.24, mean Uncertain Outcome Anticipation Index = +0.27; both significantly greater than 0, signed-rank tests, both p < 0.0001). These two indexes are generally consistent across neurons with relatively low variability so there is no significant correlation between them (rho = −0.27, p = 0.195). By contrast, ACC and Pal populations included subsets of neurons with different signs and variable strengths of uncertainty coding in standard uncertainty tasks (Fig S1). As a result, information-anticipatory activity should not necessarily lead to non-zero population average indexes, but should result in the two indexes being correlated, indicating that individual neurons anticipate both informative cues and uncertain outcomes in similar manners. Indeed, in both areas there is a strong and significant correlation such that cells with a more positive Informative Cue Anticipation Index also have a more positive Uncertain Outcome Anticipation Index (ACC: p < 0.001; Pal: p = 0.011) – the same pattern seen in Fig 3D for the population of all uncertainty-responsive neurons across the network. (**D**) Mean fraction of trials of each type in which a lick was detected in the 500 ms before cue onset. Data are from n=18 sessions in which licking was measured and there was reliable differential licking that distinguished 100% reward from 0% reward trials. Licks occurred before the cue on ∼5% or fewer trials, indicating that animals had little or no expectation that juice would be delivered at the time of the cue. Notably, the Info 50% reward CS evoked intense gaze and neural activity in anticipation of the informative cue (Fig 2-4), but evoked near-zero licking, indicating that the information-related behavior and neural activity cannot be accounted for by expectation of juice reward. If anything, there was slightly but significantly less licking for the Info 50% reward CS than for the other Info CSs. There were significant but small differences in licking between some other conditions as well (Noinfo 50% vs 0%, p = 0.005; Info 50% vs 0%, p = 0.001; Info 50% vs 100%, p = 0.032). (**E**) Same as (A), for the 500 ms before outcome delivery. Licks occurred on a large fraction of trials, indicating that animals expected juice to be delivered at the time of the outcome. Licking was generally consistent with the mean reward in each condition. On Noinfo trials licking significantly increased with reward probability (100% > 50%, p = 0.003; 50% > 0%, p = 0.044; signed-rank tests). On Info trials licking was significantly greater after reward was cued than after no-reward was cued, both for trials in which reward was initially uncertain (50%→reward > 50%→no reward, p = 0.001) and trials when reward was initially certain (100% > 0%, p < 0.001).

**Figure S5.**
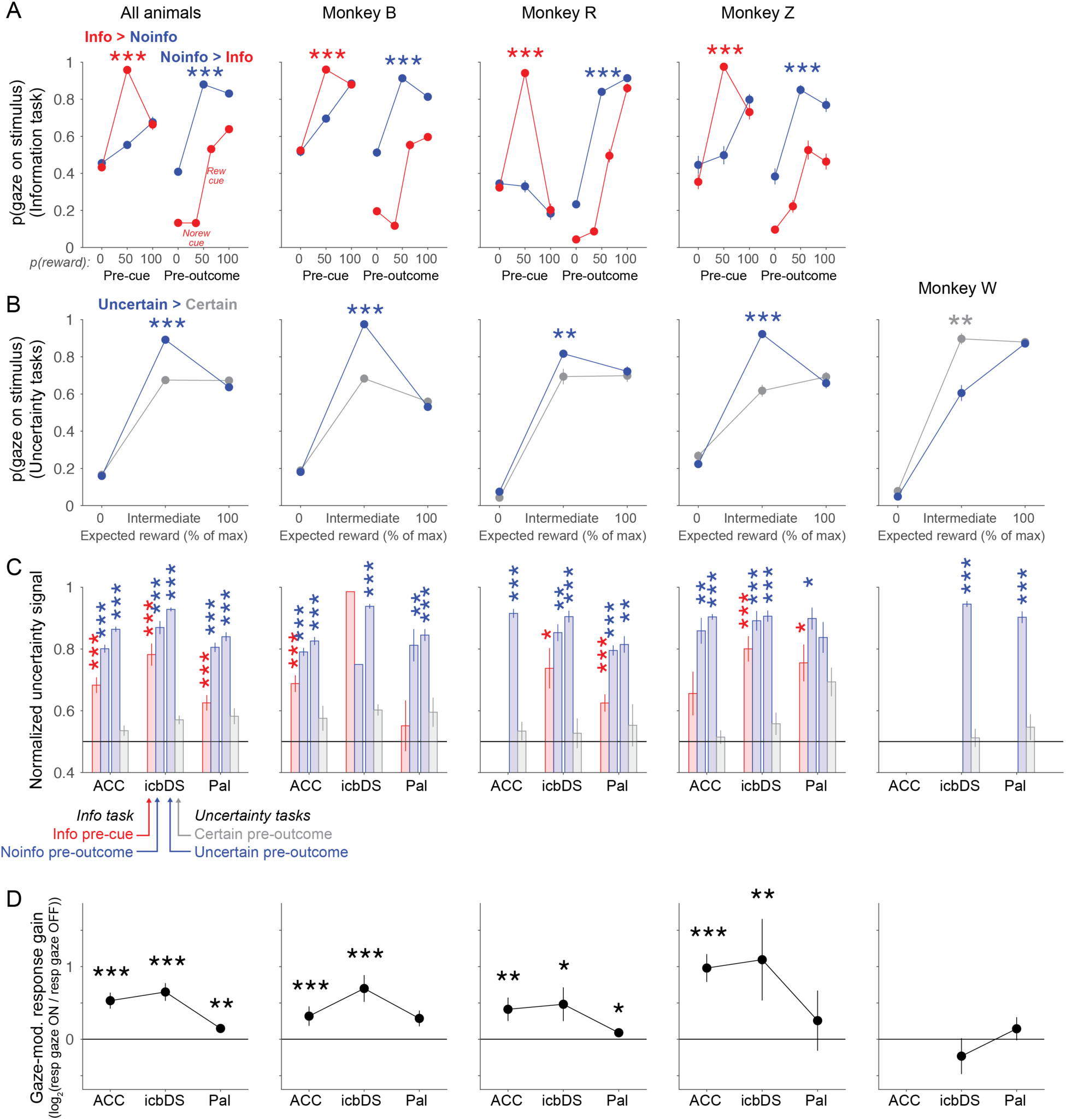
Gaze behavior and neural activity for each animal and area. The three animals that contributed the great majority of the data – animals B, R, and Z – showed strong and consistent information-anticipatory behavior in the information task (A) and standard uncertainty tasks (B); showed strong and consistent information-anticipatory neural signals in ACC, icbDS, and Pal (C); and showed similar and consistent links between fluctuations in their gaze behavior and fluctuations in the gain of their information-related neural signals (D). In addition, we were able to find a single animal (animal W) that lacked normal uncertainty-related gaze behavior, and were able to collect neural data from BG in the standard uncertainty task (B). This animal appeared to have neural uncertainty signals with the usual magnitude (C) but they lacked the usual coupling to gaze behavior (D). (**A**) Gaze behavior in the information task, averaged over all animals (left column) and in each individual animal (other columns). The fraction of the time spent with gaze on the stimulus is plotted for each Info (red) and Noinfo (blue) CS and cue condition in 500 ms time windows before the cue (left) and outcome (right). Info 50% CS pre-outcome gaze is plotted separately for trials when the no-reward cue (left data point) or reward cue (right data point) was shown. Error bars are ± SE. The pooled data from all animals shows clear information-anticipatory gaze behavior, with greater gaze on trials with uncertain outcomes than certain outcomes, significantly greater gaze pre-cue in anticipation of informative than non-informative cues (left, 50% Info CS (red) > 50% Noinfo CS (blue); *** indicates p < 0.001, signed-rank test) and significantly greater gaze pre-outcome in anticipation of uninformed than already-informed outcomes (right, Noinfo cues following Noinfo 50% CS (blue) > Info cues following Info 50% CS (red); p < 0.001). Similar behavior occurred in all three animals (B, R, and Z). (**B**) Gaze behavior in standard uncertainty tasks as a function of the expected reward value of the CSs, for blocks of trials where intermediate-value CSs were associated with uncertain rewards (blue, e.g. 50% chance of 0.25 mL or 0 mL juice) or certain rewards with equal expected value (gray, e.g. 100% chance of 0.125 mL juice). Gaze was measured in a 500 ms time window before the outcome. Animals B, R, and Z had significantly higher gaze at uncertain CSs than at certain CSs with the same expected value (*, **, *** indicate p < 0.05, 0.01, 0.001, signed-rank tests), consistent with their information-anticipatory behavior in the information task. By contrast, animal W had significantly lower gaze at uncertain than certain CSs. (**C**) Population average neural uncertainty signals in all animals, areas, and tasks. Uncertainty signals in each area are quantified as ROC areas for distinguishing between task conditions, as follows. Information task: Bar 1 (red), pre-cue Info uncertain vs certain CSs; Bar 2 (blue), pre-outcome Noinfo uncertain vs certain CSs. Standard uncertainty tasks: Bar 3 (blue), uncertain reward blocks, pre-outcome uncertain CSs vs 0,100% CSs; Bar 4 (gray), certain reward blocks, pre-outcome intermediate value CSs vs 0,100% CSs. Asterisks indicate significance (*, **, *** indicate p < 0.05, 0.01, 0.001, signed-rank tests). The pooled data from all animals shows that strong and significant uncertainty signals are present in all areas both pre-cue on Info CSs trials (red) and pre-outcome on Noinfo CS trials and in uncertain reward blocks of standard uncertainty tasks (blue), and that no significant uncertainty signals are present in certain reward blocks in standard uncertainty tasks (gray). Similar patterns are found in all animals, areas, and tasks, although not all reach significance due to small samples sizes in some cases. Notably, strong, significant pre-outcome uncertainty coding was present in all animals, including those whose gaze was attracted either uncertain CSs (B, R, Z) and certain CSs (W). (**D**) Median fitted gaze-modulation gain parameters in all animals and areas. Gains are expressed in log units (log_2_(rate when gaze is ON / rate when gaze is OFF)) and are quantified as population medians ± bootstrap SE. Asterisks indicate significance (signed-rank tests). The pooled data from all animals shows large increases in neural response gain around gaze shifts onto the CS in both ACC and icbDS, and a more modest but still highly significant increase in Pal. Similar patterns were seen in all animals whose gaze was attracted to uncertain CSs (B, R, and Z). However, animal W whose gaze was significantly less attracted to uncertain CSs lacked significant neural gain changes around the time of gaze shifts. Notably, the gain change in icbDS was lower in animal W than in all three other animals (p = 0.001, 0.056, and 0.001 vs animals B, R, and Z, respectively).

**Figure S6.**
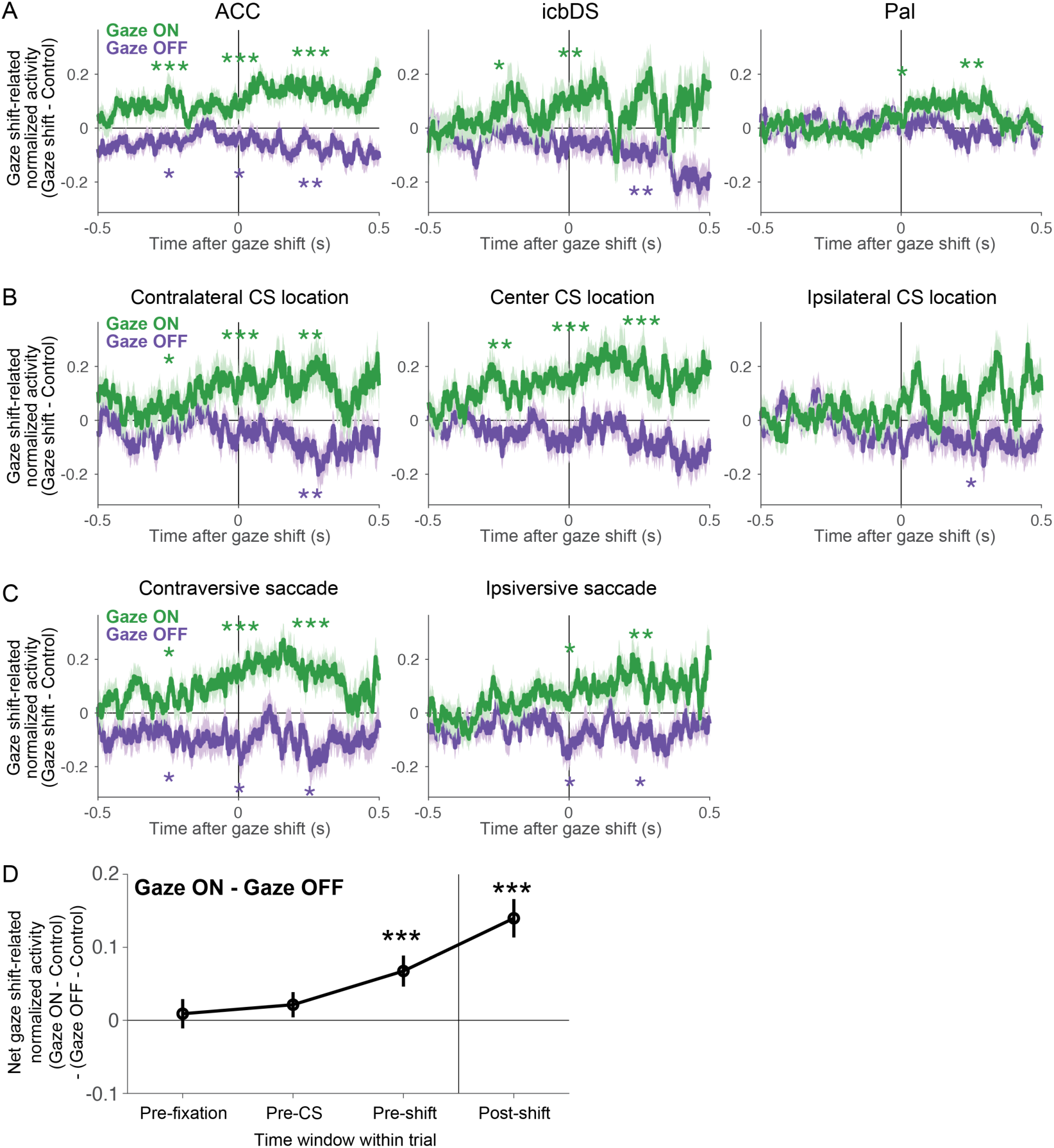
Neural activity during gaze shifts for each area and gaze shift type. (**A**) Population average changes in normalized activity related to gaze shifts, separately for each area. Same format as Fig 5D, but only showing data from gaze shifts when reward was uncertain (since gaze-related activity was much weaker when reward was certain (Fig 5)). Data is shown for all neurons with at least one gaze shift onto and one gaze shift off of the stimulus, and appropriate control non-shift events for each of these shifts, when reward was uncertain (ACC n=191, icbDS n=113, Pal n=121). All areas have normalized activity that increases around gaze shifts onto the stimulus (green) and decreases around gaze shifts off of the stimulus (purple), with these effects growing over the course of the gaze shift. The effects tend to reach significance first in ACC and later in icbDS and Pal, consistent with the latencies from the model-based analysis (Fig 6). All areas at least one effect reaching significance before or during the gaze shift (i.e. too soon for effects to have been responses to altered visual feedback after the gaze shift). (**B**) Population average changes in normalized activity related to gaze shifts, shown separately for CSs presented on each of the three possible positions on the screen (contralateral to the recorded neuron, center of screen, or ipsilateral to the recorded neuron). To allow valid comparisons between the curves, this plot is calculated using the n=131 neurons that could contribute data to all six curves (i.e. neurons that had one gaze shift onto the CS and off of the CS, and appropriate non-shift controls, in each of the three locations). Gaze-related activity was similar for all CS locations. There is a trend for stronger and earlier activity for gaze shifts onto contralateral and central CSs compared to ipsilateral CSs, but it did not reach significance. Importantly, if neural activity was simply encoding horizontal gaze location or stimulus location rather than motivational features of the stimuli, then activity would be substantially different after the gaze shift depending on whether the gaze was held on a CS located at the contralateral, central, or ipsilateral locations. Instead, neural activity had similar trends to be enhanced when gaze was at all the three locations, and to be suppressed after gaze shifts away from all three locations (no significant difference between locations in paired comparisons between gaze-shift related activity after the gaze shift, all p > 0.05 for both gaze shifts on and off the stimulus, signed-rank tests). This indicates that neural activity was not simply encoding gaze location or stimulus location. (**C**) Same as (B), but split based on whether the gaze shift’s saccade vector was contraversive or ipsiversive to the recorded neuron. To ensure that all gaze shifts were clearly contraversive or ipsiversive, this analysis is restricted to gaze shifts with magnitude >= 5°. Again, to allow valid comparisons between the curves, this plot is calculated using the n=134 neurons that could contribute data to all four curves. Gaze-related activity was similar for all saccade vectors. Importantly, if neural activity was simply encoding saccade execution or saccade direction rather than motivational features of the stimuli, then they should have similar saccade-related activity regardless of whether the saccade was toward or away from a CS (if encoding saccade execution) and this activity should preferentially occur for specific saccade directions (if encoding saccade direction). Instead, activity was significantly enhanced during and after all saccades toward CSs but was significantly suppressed during and after all saccades away from CSs, and this occurred regardless of whether the saccade was contraversive or ipsiversive. This indicates that neural activity was not simply encoding saccade execution or direction. (**D**) Demonstration that gaze shift-related activity was closely time-locked to gaze shifts, in a manner suitable for regulating ongoing gaze behavior. This is important because we found that gaze-related activity was present for > 500 ms before and after the gaze shift (A-C). This raised the possibility that neural activity might also be related to gaze shifts over much longer timescales. For instance, neurons might have slow fluctuations in neural firing rate lasting for several minutes that covary with similarly slow fluctuations in the animal’s tendency to make certain kinds of gaze shifts or to be generally engaged in the task. To test this, we calculated the mean gaze shift-related change in activity using activity in four different 500 ms time windows on each trial when the gaze shifts occurred: (1) before the onset of the fixation point, (2) before CS onset, (3) immediately before the gaze shift, (4) immediately after the gaze shift. We calculated this separately for gaze shifts onto the stimulus and gaze shifts off of the stimulus, and then calculated the difference between them (gaze shift on – gaze shift off) as a summary measure of the net strength of activity related to gaze shifts. To allow valid comparison between the data points, this analysis used the n=302 neurons that could contribute data to all data points in all curves. The net gaze shift effect was not significantly different from zero in the pre-fixation and pre-CS time windows, but was significantly increased in the pre-gaze shift window, and significantly increased further in the post-gaze shift window. Thus, changes in neural activity were closely time-locked to upcoming gaze shifts.

**Figure S7.**
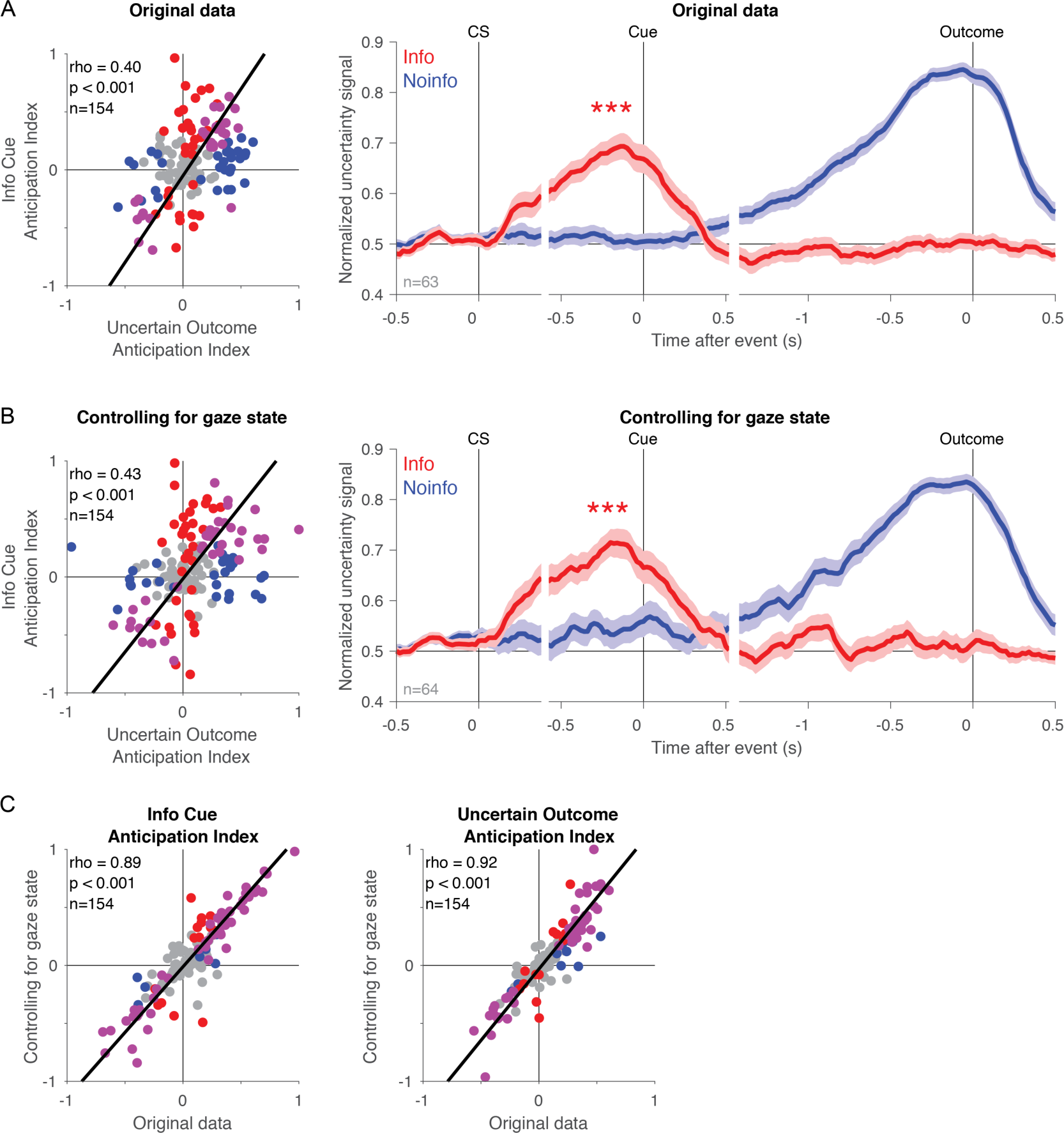
Analysis of information signals while controlling for gaze state. Our analysis in Fig 5 demonstrates that while information signals are modulated by gaze, information signals are still clearly present even after neural data is split separately into times when the animal’s gaze is on the stimulus vs. off the stimulus. This suggests that the presence of information signals cannot be explained as a mere side-effect of gaze-related activity. However, it remained possible that the average strength and/or timecourse of information signals (as quantified in our original analysis) might be substantially affected by the animal’s gaze behavior. Animals have characteristic patterns of gazing on vs. off the stimulus at each time of the trial and on each trial type. To control for this, we used our model of gaze-modulation from Fig 6. We used the fitted model for each neuron to remove the estimated influence of gaze on neural activity. Essentially, we adjusted each neuron’s activity to be ‘as if’ the gaze state had been constant at all times on all trials. Specifically, for each neuron and each millisecond of each trial, we used the neuron’s fitted model to estimate R_real gaze_, what its mean firing rate should be at each millisecond of each trial, given that trial’s true time-varying gaze behavior. We also estimated R_constant gaze_, what its mean firing rate *would have been* if the animal’s gaze state had been constant on the stimulus throughout the trial. We then calculated the difference between these values, which represents the model’s prediction about how gaze was affecting the neuron’s firing rate at that moment in time (i.e. R_gaze effect_ = R_real gaze_ – R_constant gaze_). Finally, we subtracted this gaze effect from the neuron’s original firing rate (i.e. R_controlling for gaze state_ = R_original_ – R_gaze effect_). We then repeated our analysis on this gaze state-controlled dataset. (**A**) Information signals in the original dataset. Left: same as Fig 3D. Right: same as Fig 3C (**B**) Information signals in the dataset controlling for gaze state. The results are highly similar to the original data. The only noticeable effect is a mild increase in uncertainty signals throughout the task, particularly at times when the animal’s gaze was often away from the stimulus in the original dataset. There was little or no effect at the key moments when animals were anticipating information, as expected because the animal’s gaze was already almost constantly on the stimulus at those moments in the original dataset. (**C**) The Informative Cue Anticipation Index and Uncertain Outcome Anticipation Index are highly correlated between the two datasets (rho > 0.88, both p < 0.001).

**Figure S8.**
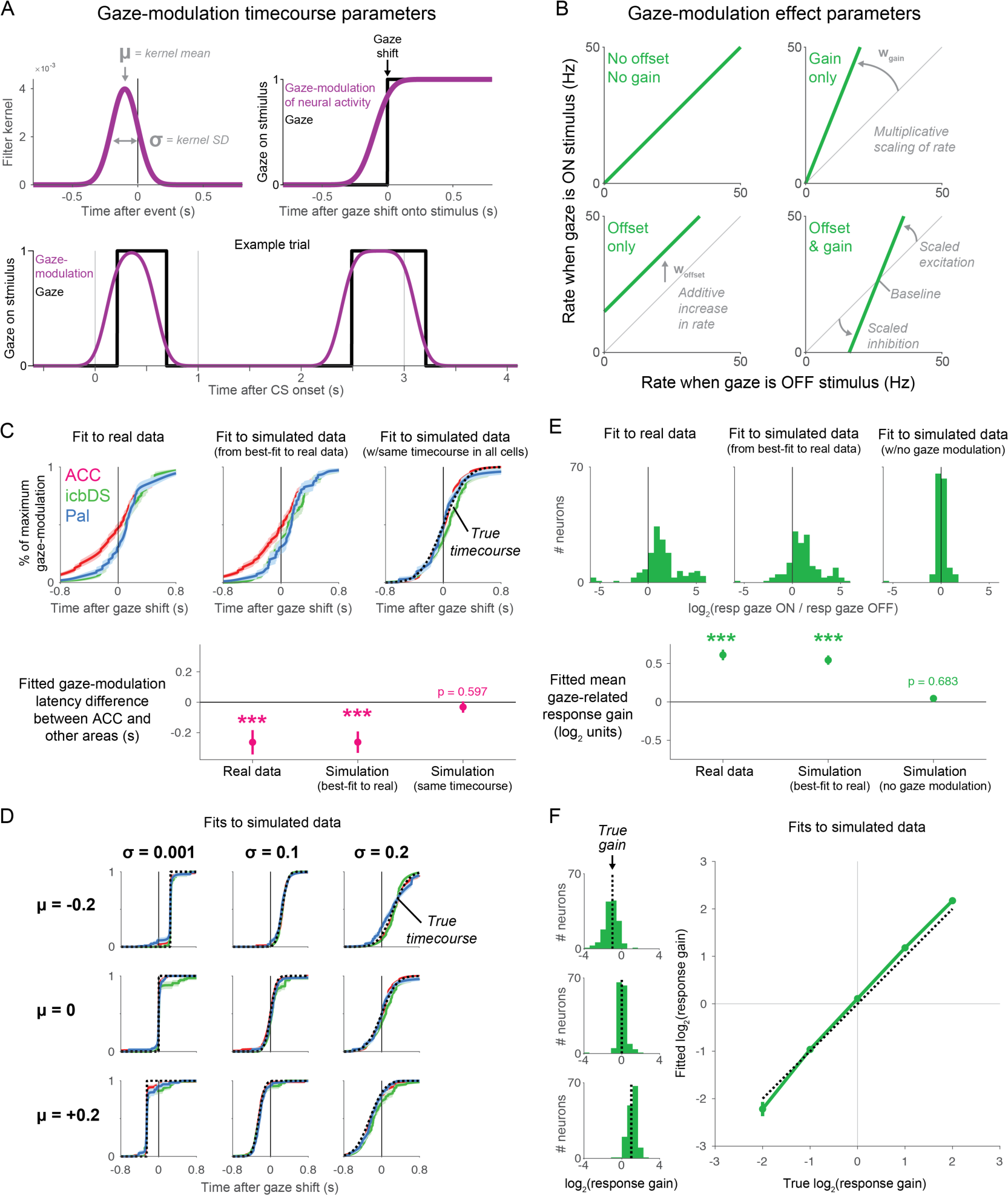
Model of gaze-related modulation in single neurons. The model fitted each neuron’s activity as the combination of three factors: fitted timecourse of neural response to each trial type (defined by the trial’s CS and cue) when the gaze is off the stimulus, fitted timecourse of gaze-modulation from when gaze is on the stimulus (A), and fitted effect of gaze-modulation (B). Our simulations indicate that the model fits accurately recover a close approximation of the true timecourse and effect of gaze modulation. This is true for simulations mimicking the properties of the real data (C,E), simulations representing null hypotheses that gaze effects are absent (C,E), and simulations representing alternative hypotheses that timecourses and effects of gaze modulation are different from those seen in the real data (D,F). (**A**) Model parameters controlling the timecourse of gaze-modulation (**µ** and **σ**). Gaze (black) was defined at each millisecond of each trial as 1 if the eye was within 3° of the center of the trial’s stimulus location and 0 otherwise. GazeMod (purple) was defined by smoothing Gaze with a Gaussian kernel specified by two parameters, the mean **µ** and standard deviation **σ** (*top left*). Thus, after a gaze shift onto the stimulus, the timecourse of GazeMod was a sigmoid, cumulative Gaussian function rising from 0 to 1, with its temporal shift controlled by **µ** and the gradualness of its rise controlled by **σ** (*top right*). For example, during the same example trial shown in Fig 5, the model’s GazeMod goes up and down in temporal alignment with changes in Gaze (*bottom*). GazeMod then controls the degree to which firing rate is affected by the parameters in (B). For instance, when GazeMod = 0 firing rate is unaffected by gaze; when GazeMod = 0.5 firing rate is affected half-maximally by gaze; when GazeMod = 1 firing rate is maximally affected by gaze. (**B**) Model parameters controlling the effect of gaze-modulation (**w_gain_** and **w_offset_**). These allow GazeMod to cause a multiplicative scaling of neural responses (**w_gain_**), an additive increase or decrease in neural activity (**w_offset_**), or both. Specifically, the model’s firing rate is specified by the equation Rate = Rate_GazeOFF_*(1 + GazeMod\***w_gain_**) + GazeMod* **w_offset_**. Thus, if gaze is on the stimulus, it has different effects on firing rate depending on these two parameters. If **w_gain_**= 0 and **w_offset_** = 0 then firing rate is unaffected (*top left*). If **w_gain_** > 0, neural activity is multiplicatively scaled up in proportion to **w_gain_**(*top right*). If **w_offset_** > 0, neural activity is additively increased by **w_offset_** (*bottom left*). Finally, if both **w_gain_** and **w_offset_** are non-zero, then neural responses (i.e. deviations in neural firing rate from a baseline level) are multiplicatively scaled up in proportion to **w_gain_** (*bottom right*). (**C**) Model performance: fitted timecourses of gaze-modulation for real and simulated data. Fits are shown for the real data (*left*), a simulated dataset generated from the best-fitting model to the real data (*middle*), and a second simulated dataset generated from the same model except that its gaze-modulation timecourses are forced to be the same, identical curves for all neurons in all areas (*right*, enforced timecourse indicated by black dashed curve). Simulated data was generated from a given model by calculating its predicted mean spike count for each time bin of each trial, then drawing the corresponding simulated spike count from a Poisson distribution with that mean. Fitting results are displayed as the average over all neurons that were significantly modulated by gaze in the real data. The fit to the real data indicates that gaze-modulation begins earlier in ACC than icbDS or Pal (*left*). The fits to the simulated data indicate that the model-fitting procedure is effective at recovering the true timecourse of gaze-modulation. Thus, in a simulation where the true timecourse is known to be equal to the curves on the left, the fits accurately recover a close approximation of those curves (compare *middle* vs *left*), including a similarly significantly earlier latency in ACC than icbDS and Pal (*bottom*). Likewise, in a simulation where the true timecourse is known to be identical in all neurons in all areas, the fits in all areas accurately recover a close approximation of that timecourse (*right*, compare colored curves to black curve), resulting in no difference in latency between areas (*bottom*). (**D**) Model performance: fitted timecourses for simulated data with a variety of true timecourses. As in the right panel in (C), each dataset was simulated from the best-fitting model to the real data, but with its gaze-modulation timecourse parameters forced to be the same for all neurons in all areas. The true timecourses had **µ** parameters making them occur before, simultaneous, or after gaze shifts (top, middle, bottom) and **σ** parameters making them be instantaneous, gradual, or very gradual (left, center, right). The model recovered close approximations of the true timecourse of gaze modulation (even when the true timecourse was very temporally sharp (left)) and correctly fitted similar timecourses to all three areas (all colored curves similar to the black curve). (**E**) Model performance: fitted effect of gaze modulation for real and simulated data. Fits are shown for the real data (*left*), a simulated dataset generated from the best-fitting model to the real data (*middle*), and a second simulated dataset generated from the same model except that it has no gaze-modulation because **w_gain_** and **w_offset_** parameters are both set to zero for all neurons in all areas (*right*). Same plotting conventions as the histogram of fitted gains in the main text. Fitting results are displayed for all neurons that were significantly modulated by gaze in the real data. The fit to the real data indicates that gaze-modulation often includes changes in response gain, typically consisting of an increase in gain but with some variability across neurons (left). The fits to the simulated data indicate that the model-fitting procedure is effective at recovering the true effect of gaze-modulation. Thus, in a simulation where the true gain changes are known to be equal to the distribution on the left, the fits accurately recover a close approximation of that distribution of gain changes (compare *middle* vs *left*), including a similarly significantly positive mean gain change (*bottom*). Likewise, in a simulation where there is known to be no true gaze-modulation, the fits accurately indicate that all neurons have little or no change in gain (*right*) and hence that the mean gain change is not significantly different from zero (*bottom*). (**F**) Model performance: fitted effect of gaze modulation for simulated data with a variety of true effects. *Left*: distribution of gain changes fit to single neurons in simulations where the true log_2_(gain) was set to −1, 0, or 1 for all neurons (black dashed lines), meaning that neural responses are halved, unchanged, or doubled during gaze. The distributions of fitted gain changes are clustered around the true gain change (compare green distribution to black dashed lines). *Right*: relationship between true and mean fitted gain change across multiple simulations. The fitted gain changes (green) approximate the true gain changes (black dashed identity line).

**Figure S9.**
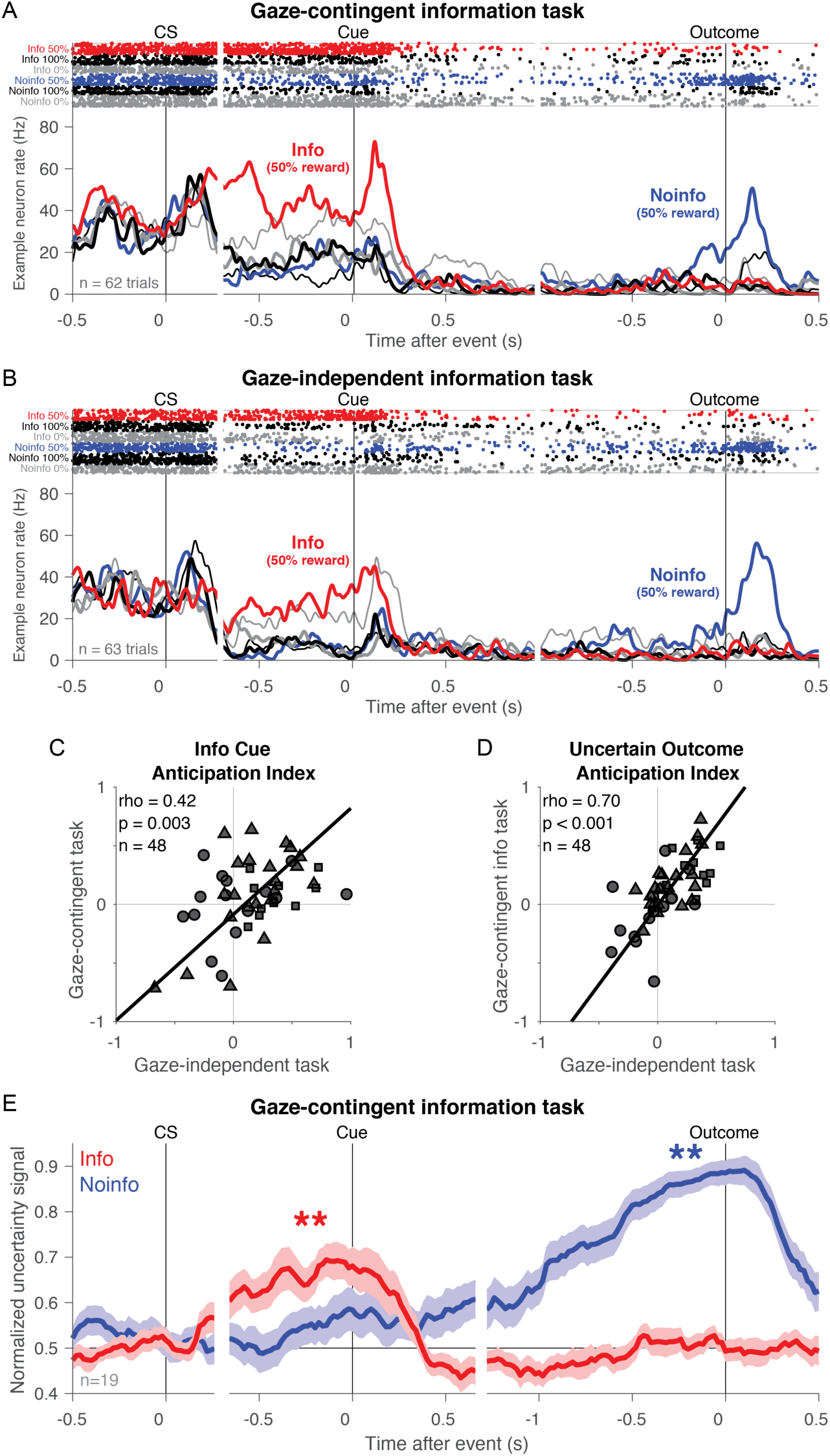
Information-anticipatory signals are present when access to information is contingent on gaze. (**A,B**) Activity of an example ACC neuron recorded in both the gaze-contingent information task (A) and the gaze-independent information task (B). Colored lines of different thicknesses indicate activity on Info trials (red thick, black thick, gray thick: 50%, 100%, and 0% reward) and Noinfo trials (blue thick, black thin, gray thin: 50%, 100%, and 0% reward). This neuron had similar information-related activity in both tasks, with clear information-anticipatory activity during the CS period in both tasks. Notably, in anticipation of cue onset, its firing rate on Info 50% trials was higher than its firing rates on Noinfo 50% trials, on Info Certain trials, and on Noinfo Certain trials (p = 0.0047, p = 0.0044, p = 0.0138 respectively in the gaze-contingent task, and p = 0.0011, p = 0.0002, and p = 0.0042 in the gaze-independent task). (**C,D**) Coding indexes are highly similar in the two tasks. Shown are the correlations between the gaze-contingent task and gaze-independent task measurements of the Info Cue Anticipation Index (C) and Uncertain Outcome Anticipation Index (D). Both are significantly positively correlated. Symbols indicate areas (triangle, square, and circle indicate ACC, icbDS, and Pal). For completeness and to any potential for avoid selection bias, this analysis includes all neurons recorded in both tasks; we obtained similar results if analyzing only neurons with significant uncertainty coding (n=40, Info Cue Anticipation Index rho = +0.35, p = 0.026; Uncertain Outcome Anticipation Index rho = +0.68, p < 0.001). (**E**) Timecourse of normalized uncertainty signals in the gaze-contingent task for uncertainty coding neurons with significant Uncertain Outcome Anticipation Indexes. Same format as Fig 3C. Information-anticipatory activity is significantly present before the cue (p = 0.01, signed-rank test). Note that this data further confirms our finding (Fig 5A, Fig S7) that neural information-anticipatory signals are not merely an artifact of allowing the animal to gaze freely at the stimulus. Information-anticipatory activity occurred during the CS period of the gaze-contingent task – a time when the animal was not allowed to gaze at the CS, because they were required to maintain fixation at the center of the screen until they received the ‘go’ signal. Thus, information-anticipatory activity occurred when animals were anticipating information, even at times when they were not allowed to express their state of anticipation with an action.

**Figure S10.**
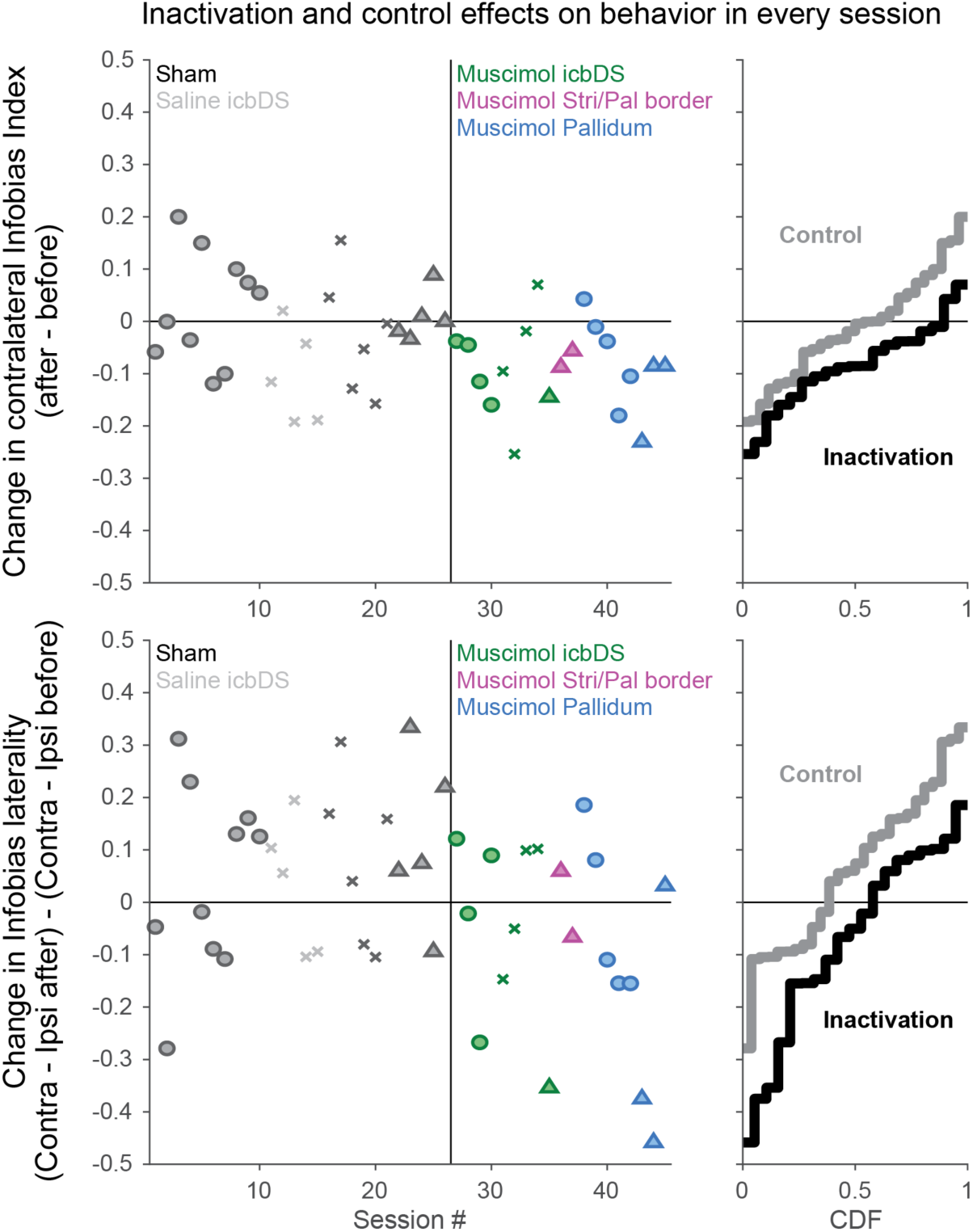
Inactivation results from each session. Sessions are sorted by type (sham, saline, muscimol), animal (B, R, Z), and area (icbDS, Pal, Stri/Pal border region). The two Stri/Pal border region injections were excluded from our main analysis because they were done in a border region between the striatum and pallidum that was not clearly identifiable as strictly icbDS or Pal (between the ventral portion of icbDS and the dorsal portion of the anterior globus pallidus above the anterior commissure; Table S2). However, they did not alter our results or conclusions if they were included in our analysis and are shown here for completeness. *Top:* change in Infobias Index for contralateral CSs. Left: data for single sessions. Right: cumulative distributions of the data plotted on the left. Inactivations resulted in a more negative change in Infobias Index relative to control (as shown in Fig 7). *Bottom:* change in laterality of Infobias Index (Infobias Index contra – Infobias Index ipsi). Inactivations resulted in the laterality of information seeking behavior being shifted more negatively (i.e. away from the contralateral side) relative to control (as shown in Fig S11C).

**Figure S11.**
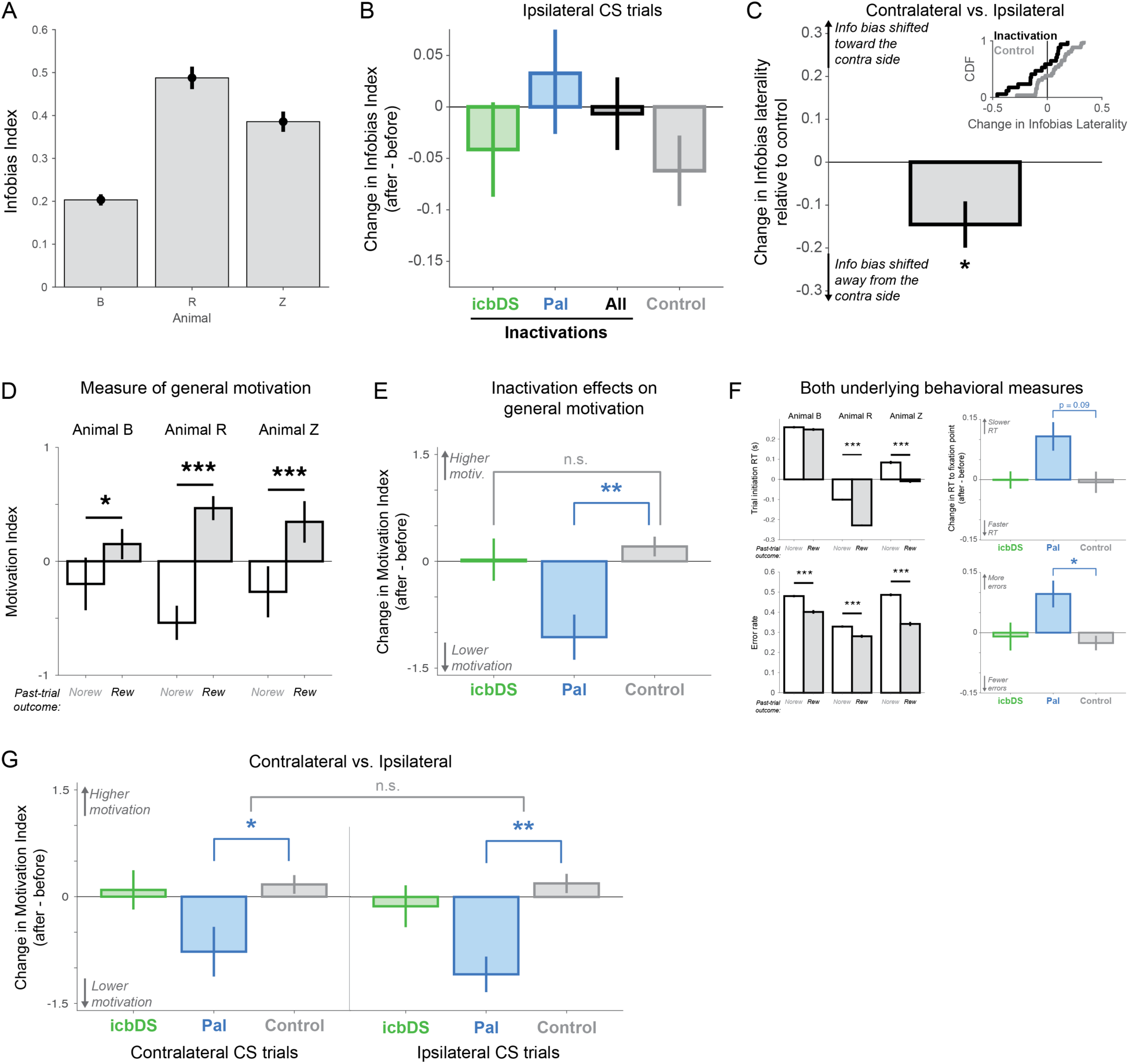
Inactivation effects on laterality of information seeking behavior, general motivation, and effects in each session. (**A**) Mean Infobias Index in each animal. All animals had significantly positive Infobias Indexes in the mean over all sessions shown here and in all individual sessions (n=43/43; all p < 0.05, permutation tests). (**B**) No significant change in Infobias Indexes for ipsilateral CSs. Same format as Fig 7C, but for ipsilateral CSs. There is no significant change resulting from inactivations and no significant difference between inactivations and control. (**C**) Inactivations change the laterality of information seeking behavior relative to control sessions. *Inset*: cumulative distributions of change in infobias laterality for inactivations (black) and controls (gray). Change in infobias laterality was quantified as: (change in Infobias Index for contralateral CSs) – (change in Infobias Index for ipsilateral CSs). *Bar plot*: difference in the change in infobias laterality between inactivations and controls. Error bars are ± bootstrap SE. The change in infobias laterality was more negative for inactivations than controls (p = 0.019, permutation test), indicating that information seeking behavior was shifted away from the contralateral side relative to controls. (**D-G**) **Inactivation effects on general motivation and reward responsiveness.** We quantified the animal’s motivation to perform the task using two measures of general motivation that have been used in previous studies: the response time to initiate the trial on correctly performed trials, and the probability of making an error during the trial ^35, 101, 102^ (F). To summarize these measures and gain statistical power to detect any potential small effect of inactivation on motivation, we created a composite Motivation Index (D,E) pooling these measures by z-scoring each measure within each animal, averaging the two measures within each session, and then flipping its sign (so that positive Motivation Index indicates higher motivation to perform the task). (**D**) Mean Motivation Index for each animal, plotted separately for trials in which the previous trial was rewarded (black) or non-rewarded (white). If the Motivation Index is a valid indicator of the animal’s motivation then we expect it to indicate that most animals are more motivated to perform the task after receiving rewards. This result is commonly reported in monkeys unless tasks are designed to specifically encourage the opposite behavior ^103^. Indeed, this was the case. By definition the mean Motivation Index over all trials is near zero, but all animals had a significantly higher Motivation Index after receiving reward than no reward (rank-sum tests, all p < 0.05). (**E**) Mean change in Motivation Index (after – before), plotted separately for icbDS inactivation, Pal inactivation, and control sessions (colored bars). During control sessions there was no significant change in measures of motivation. icbDS inactivation sessions also had no apparent change in motivation in any measure, and their change in Motivation Index was not significantly different from control (p = 0.414, permutation test). Pal inactivation sessions had a trend for reduced motivation and their change in Motivation Index was significantly different from control (p = 0.008, permutation test). (**F**) Similar results occurred for both individual motivational measures. When the past trial was rewarded, animals had faster trial initiation RTs and lower error rates (rank-sum tests, all p < 0.001 except for the trial initiation RT in animal B). Conversely, Pal inactivations tended to induce slower trial initiation RTs and higher error rates relative to control sessions, as either significant effects or non-significant trends (fixation RT, p = 0.092; error rate, p = 0.018). Note that the error rate includes both trials while the animal failed to fixate and trials when the animal fixated but then broke fixation. We obtained similar results when considering either one of these two types of errors individually (only failures to fixate, p = 0.028; only fixation breaks, p = 0.043). Note that the trial initiation RTs could be negative. This occurred when the animal gazed at the fixation window in an anticipatory manner before the fixation point appeared. This behavior appeared to reflect a situation in which the animal was highly motivated to complete the task, animals did this most frequently if they had received a reward on the previous trial (especially in animals R and Z), and this caused the trial to be initiated at the earliest possible time. This property of the data was not critical for our results; we obtained the same pattern of results in this figure if we enforced non-negative trial initiation RTs by setting all negative RTs to 0. (**G**) Similar results occurred for both locations in space. On both contralateral CS trials and ipsilateral CS trials, the Motivation Index was unaffected by icbDS inactivation (contra p = 0.67, ipsi p = 0.253, no significant difference between them, p = 0.573) and was similarly and significantly reduced by Pal inactivation (contra p = 0.028, ipsi p = 0.004, no significant difference between them, p = 0.170; permutation tests).

**Figure S12.**
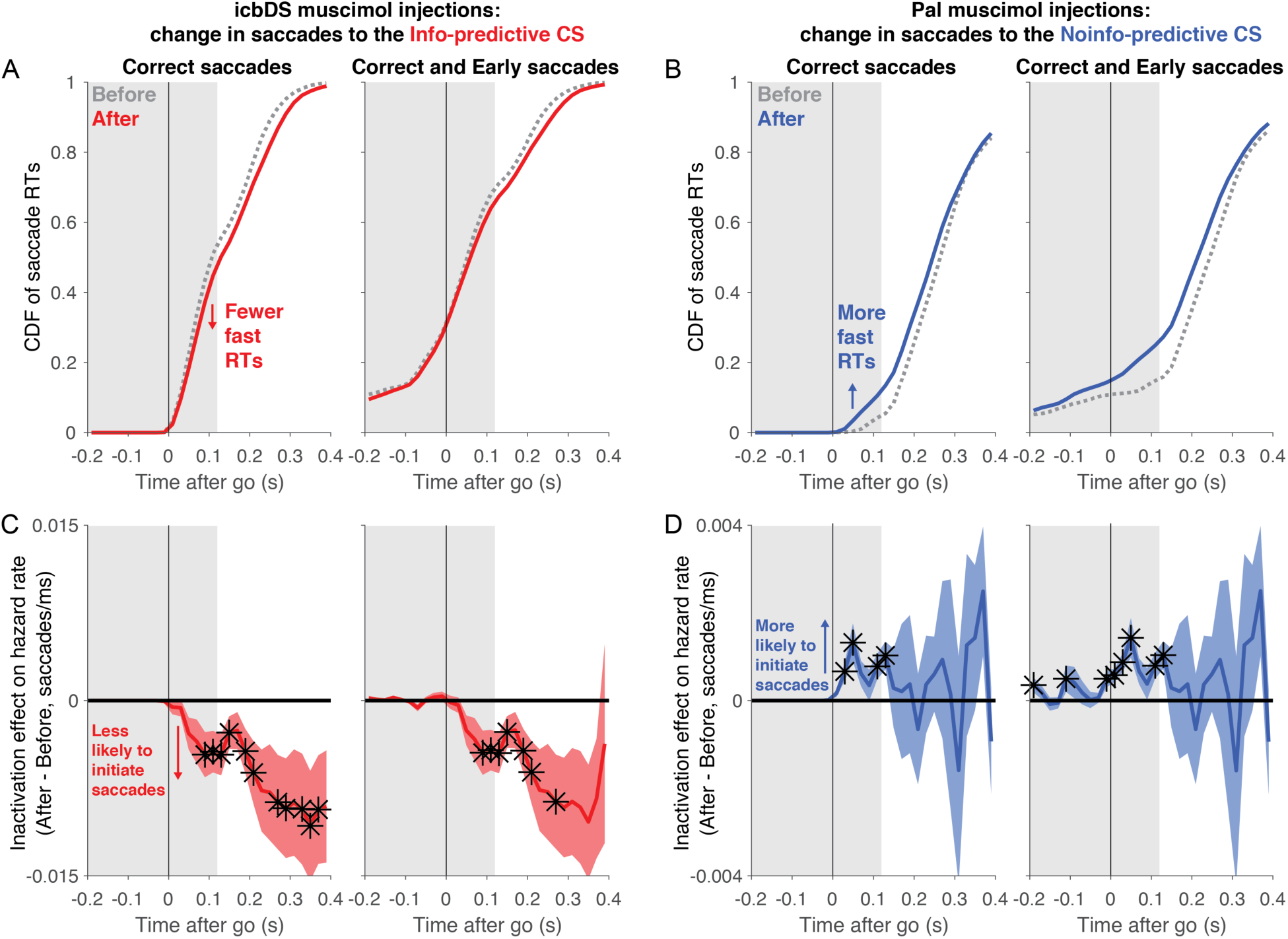
Inactivation of icbDS and Pal influences anticipatory behavior. Our analysis thus far showed that inactivations of icbDS and Pal altered saccadic response times to Info and Noinfo CSs, respectively. Here we investigate in more detail how these inactivations affected saccade generation and when in the trial they had their effects. In particular, did they alter anticipatory saccades the animals generated while awaiting the ‘go’ signal? To address this, we first observed that the histograms of saccadic RTs permitted a relatively clear distinction between times in the trial when saccades were predominantly ‘anticipatory’ vs. ‘reactive’. In the example histogram in Fig 7B, the initial ∼120 ms after the go signal had a relatively low rate of responses, which was stable or gradually increasing over time. This is suggestive of anticipatory saccades, prepared based on the animal’s internal estimate of when the go signal would occur. Then at ∼120-150 ms there was a sudden, sharp increase in the response rate. This is suggestive of reactive saccades, generated as a reaction to the onset of the go signal. To quantify this distinction, we wanted to estimate the time after the go signal when behavior switched from anticipatory to reactive. Therefore, we made a conservative estimate of the minimal reactive RT. We built a histogram of all contralateral RTs and estimated the onset of the sharp peak by finding the millisecond after the go signal at which there was the largest difference between the frequency of saccades in the next 15 ms compared to the previous 15 ms. To be conservative, we then subtracted 15 ms from this estimate to ensure that it excluded the great majority of reactive saccades. The resulting estimates of the minimal reactive RT in this task were reliable and consistent across animals (124 ± 2 ms, 119 ± 2 ms, 124 ± 1 ms in animals B, R, and Z, respectively; mean estimate ± bootstrap standard error). Therefore, in this analysis we examined the timecourse of inactivation effects on saccade generation and whether these effects were present at times when saccades were predominantly anticipatory (< 120 ms after the go signal, gray shaded area in panels A-D). We examined inactivation effects on the cumulative distribution of RTs across time (**panels A,B**) and for a more precise measurement, we quantified their effects on the instantaneous probability of saccade initiation (also known as the hazard rate), which is more direct measure of perturbation effects on behavior at each moment in time^104^ (**panels C,D**). We performed this analysis on all correct trials (**left part of panels A-D**). To gain further insight into anticipatory behavior, we repeated this analysis on all correct trials plus error trials in which the animal made a premature saccade to the CS (**right part of panels A-D**, error saccades coded as ‘negative RTs’ because the saccade occurred before the go signal). Note that our hypothesis – that icbDS and Pal activity affect the motivation to seek information – does not make a strong prediction about whether inactivations should affect the likelihood or timing of premature saccade errors. For instance, if a manipulation increases the motivation to seek information, this could *increase* the rate of premature saccades (by making the Info CS more attractive), or *decrease* the rate of premature saccades (by making the animal more motivated to complete each trial correctly in order to obtain information), or cause *no change* in premature saccades (e.g. by both effects counterbalancing each other). Nonetheless, examining these trials may give further insight into inactivation effects on anticipatory behavior. (**B**) Cumulative distribution of saccadic response times to the contralateral Info CSs before (gray) and after (red) icbDS inactivation, for all correct trials (left) and correct trials plus premature saccades (right). As expected from our findings in Fig 7, the inactivation shifted the RT distribution to have fewer fast RTs (red < gray). The CDFs visibly diverged within the first 120 ms after the go signal (gray shaded area), suggestive of an effect on anticipatory behavior. (**C**) Same, for saccades to the contralateral Noinfo CS before (gray) and after (blue) Pal inactivation. As expected, the inactivation shifted the RT distribution to have more fast RTs (blue > gray). The CDFs visibly diverged within the first 120 ms after the go signal, suggestive of an effect on anticipatory behavior. (**C,D**) Inactivation effects on the instantaneous probability of saccade initiation (i.e. the hazard rate) as a function of time relative to the go signal. This was calculated as follows. In each session and each before/after condition, we calculated the probability distribution of contralateral RTs with 1 ms precision and then smoothed it with a causal exponential kernel (τ = 30 ms; causal smoothing ensures that estimates of effects on presumed anticipatory behavior (RTs < 120 ms) are not influenced by later events (RTs > 120 ms)). We then calculated the hazard rate in the standard manner as the probability of a response at each point in time given that a response has not yet occurred (i.e. hazardrate(t) = p(RT = t) / p(RT ≥ t)). Finally, we calculated the inactivation effect on the hazard rate in each session as (hazard rate after – hazard rate before) and binned the results in 20 ms time bins. Plotted is the mean ± SE of the single session effects. Black asterisks indicate time points with significant effects (signed-rank test, p < 0.05). (**D**) icbDS inactivation effect on instantaneous probability of saccades to the Info CS (n=9 sessions). After inactivation there was a substantial reduction in saccade probability. In the analysis of correct trials (left) this effect started almost immediately after the go signal and continued for several hundred milliseconds. Notably, the effect first reached significance in the time bins starting at 80, 100, 120, and 140 ms after the go signal, overlapping with the gray shaded area. No significant effects were found at these or nearby times in the analogous analyses of control sessions, or for ipsilateral saccades on inactivation sessions. Similar results occurred in the analysis that included premature saccades (right). This suggests an effect on anticipatory saccades. (**E**) Pal inactivation effect on instantaneous probability of saccades to the Noinfo CS (n=8 sessions). After inactivation there was a substantial increase in saccade probability. In the analysis of correct trials (left) this effect reached significance almost immediately after the go signal and at later times as well (time bins starting at 20, 40, 100, and 120 ms), suggestive of anticipatory behavior. No significant effects were found at these times or nearby times in the analogous analyses of control sessions, or for ipsilateral saccades on inactivation sessions. In the analysis including premature saccades (right), this effect also reached significance at some times before and adjacent to the go signal (time bins starting at −200, −120, −20, 0, and 20 ms). This suggests an effect on anticipatory saccades.

**Table S1.** Neural dataset from each animal and task. Shown for each animal (B, R, Z, and W) and area (ACC, icbDS, and Pal) are the total number of neurons recorded, the number of neurons recorded in 1, 2, or 3 different tasks, and the number of neurons recorded in each of the tasks (standard uncertainty tasks A-E and information tasks IA-ID).

| Animal | Area | #cells | #tasks recorded |  |  | standard uncertainty tasks |  |  |  |  | information tasks |  |  |  |
| --- | --- | --- | --- | --- | --- | --- | --- | --- | --- | --- | --- | --- | --- | --- |
|  |  |  | 1 | 2 | 3 | A | B | C | D | E | IA | IB | IC | ID |
| All | All | 484 | 423 | 53 | 8 | 140 | 112 | 87 | 25 | 25 | 77 | 67 | 42 | 36 |
|  | ACC | 222 | 204 | 12 | 6 | 52 | 33 | 78 | 15 |  | 22 | 24 | 42 |  |
|  | icbDS | 127 | 103 | 23 | 1 | 63 | 49 | 9 |  | 7 | 24 | 24 |  |  |
|  | Pal | 135 | 116 | 18 | 1 | 25 | 30 |  | 10 | 18 | 31 | 19 |  | 36 |
| B | All | 204 | 174 | 25 | 5 | 56 | 45 | 45 | 25 |  | 22 | 22 | 42 |  |
|  | ACC | 113 | 101 | 8 | 4 | 12 | 9 | 36 | 15 |  | 13 | 13 | 42 |  |
|  | icbDS | 58 | 44 | 13 | 1 | 34 | 29 | 9 |  |  | 1 | 1 |  |  |
|  | Pal | 33 | 29 | 4 |  | 10 | 7 |  | 10 |  | 8 | 8 |  |  |
| R | All | 117 | 104 | 13 |  | 27 | 12 |  |  | 25 | 27 | 15 |  | 36 |
|  | ACC | 24 | 23 | 1 |  | 17 | 8 |  |  |  |  |  |  |  |
|  | icbDS | 22 | 21 | 1 |  | 6 |  |  |  | 7 | 10 | 10 |  |  |
|  | Pal | 71 | 60 | 11 |  | 4 | 4 |  |  | 18 | 17 | 5 |  | 36 |
| Z | All | 114 | 97 | 14 | 3 | 44 | 18 | 42 |  |  | 28 | 30 |  |  |
|  | ACC | 85 | 80 | 3 | 2 | 23 | 16 | 42 |  |  | 9 | 11 |  |  |
|  | icbDS | 22 | 14 | 8 |  | 16 | 1 |  |  |  | 13 | 13 |  |  |
|  | Pal | 7 | 3 | 3 | 1 | 5 | 1 |  |  |  | 6 | 6 |  |  |
| W | All | 49 | 48 | 1 |  | 13 | 37 |  |  |  |  |  |  |  |
|  | icbDS | 25 | 24 | 1 |  | 7 | 19 |  |  |  |  |  |  |  |
|  | Pal | 24 | 24 |  |  | 6 | 18 |  |  |  |  |  |  |  |

**Table S2.** Inactivation parameters from each session. Shown are each session’s animal, treatment (muscimol or saline), volume injected (uL), area injected (icbDS, Pal, or two sites that were at an intermediate location on the Border between striatum and pallidum; Fig S10), and reconstructed coordinates of the injection site (same coordinate system as in Fig S2, i.e. positions are in mm and are relative to the midline superior tip of the anterior commissure).

| Animal | Treatment | Volume (μL) | Area | Injection site relative to AC (mm) | | |
| --- | --- | --- | --- | --- | --- | --- |
|  |  |  |  | Anterior | Lateral | Superior |
| B | Muscimol | 2 | icbDS | 3.5 | 6.9 | 2.7 |
| B | Muscimol | 2.5 | icbDS | 7.5 | 6.9 | 2.7 |
| B | Muscimol | 2 | icbDS | 6.5 | 6.9 | 2.8 |
| B | Muscimol | 1 | icbDS | 6.5 | 6.6 | 2.4 |
| R | Muscimol | 3 | icbDS | 5 | 9.5 | 2.2 |
| R | Muscimol | 2 | icbDS | 3 | 10.2 | 1.4 |
| R | Muscimol | 2 | icbDS | 3 | 10.2 | 1.4 |
| R | Muscimol | 2 | icbDS | 4 | 9.7 | 0.8 |
| Z | Muscimol | 1.5 | icbDS | 4 | 10.4 | 0.5 |
| Z | Muscimol | 2.5 | Border | 1 | 6.2 | 3.3 |
| Z | Muscimol | 2 | Border | 2 | 6.8 | 2.4 |
| B | Muscimol | 1.4 | Pal | -0.5 | 9.1 | -5.4 |
| B | Muscimol | 1 | Pal | -0.5 | 8.2 | -5 |
| B | Muscimol | 0.8 | Pal | -1.5 | 9.8 | -6.2 |
| B | Muscimol | 0.8 | Pal | -1.5 | 9.8 | -6.2 |
| B | Muscimol | 0.545 | Pal | 0.5 | 8.4 | -3.2 |
| Z | Muscimol | 0.545 | Pal | -1 | 7.9 | -6.2 |
| Z | Muscimol | 0.636 | Pal | -1 | 7.9 | -6.2 |
| Z | Muscimol | 0.636 | Pal | -1 | 7.9 | -6.2 |
| R | Saline | 2 | icbDS | 4 | 9.7 | 0.8 |
| R | Saline | 2 | icbDS | 3 | 9.9 | 1 |
| R | Saline | 2 | icbDS | 3 | 9.1 | 1.6 |
| R | Saline | 2 | icbDS | 3 | 11.1 | 1.1 |
| R | Saline | 2 | icbDS | 4 | 11 | 0.9 |

### Supplementary Note: design and logic of perturbation experiments

Here we detail two methodological points: the reasoning for perturbing the targeted brain structures and the logic of the experimental design of perturbations and controls.

#### Targeting brain structures for perturbation

We targeted icbDS and Pal for perturbation because, by combining known anatomy and physiology with our own anatomical and neurophysiological findings, we were able to make strong hypotheses about how their activity may be causally linked to information seeking behavior, and hence were able to have a clear framework for interpreting the results. While our results also indicate a link between ACC and information seeking, our findings do not allow us to make a firm hypothesis about the effects of perturbing that area and what it would teach us about the underlying neural mechanisms. Specifically:

1. Our data suggest that Pal and icbDS are well-positioned to influence the network’s access to information-related signals and generation of behavioral output. By contrast, even if ACC was inactivated, our data suggest that there would likely be an intact pathway for information signals to reach icbDS and Pal and influence behavior. Notably, our latency analysis shows that (rough) uncertainty signals appear at the shortest latency in Pal, our anatomical data indicates that this could be transmitted directly to icbDS by Pal→icbDS projections, and our gaze-modulation analysis indicates that icbDS and Pal change their activity significantly closer to the time of upcoming gaze shifts than ACC, suggesting a more proximal link to behavior. Taken together, our data suggests that ACC activity may be well-suited for a supervisory role in information anticipatory behavior, but that even without ACC, icbDS and Pal may still be capable of generating simple, rough information-anticipatory signals and using them to guide behavior – which could well be sufficient to perform the task used in perturbation experiments.
2. Our study, and our model of perturbation effects on the circuit, is focused specifically on the subset of neurons that encode signals related to uncertainty. This allows us to make clear predictions for icbDS perturbations, since icbDS uncertainty-related neurons have very consistent response patterns (i.e. exclusively excitatory information signals with a relatively consistent timecourse across neurons). This also allows us to make fairly clear predictions for Pal perturbations. Pal has more diverse responses, and indeed, Pal perturbations also produced significant effects on measures of the animal’s level of general motivation (e.g. error rates). However, these are known effects that have been produced in previous studies using Pal inactivations ^32–38^. Hence, we designed our task so these would not confound our key test of information seeking. By contrast, ACC contains a diverse array of neurons related to different cognitive and motivational functions, such as the subjective value of an option, specific task-relevant features of options, rewards, punishments, etc., most of which can be encoded with either excitations or inhibitions (as shown by studies from many groups ^39, 40, 49–52^ including our own ^22^). This makes it difficult to predict what side-effects to expect, especially as ACC inactivations have been rarely reported in the primate literature. Therefore, if ACC perturbations altered our measure of information seeking, it could also have complex side-effects on motivated behavior that make the results difficult to interpret.
3. Dorsal striatum and Pal have projections within the same hemisphere. Thus, perturbations would be expected to produce effects that are (to a first order) localized to a single side of the brain, and hence preferentially affect contralateral behaviors. This is necessary for the logic of our experiments, as the comparison of ‘contra’ vs ‘ipsi’ conditions is a vital control to show that the perturbations have a specific effect on localized brain circuitry rather than a generalized effect on all behavior. By contrast, ACC projects to both hemispheres, including its own counterpart in the opposite hemisphere^105^, and these fiber pathways are capable of transferring neural activation between hemispheres^106^. Perturbations would therefore potentially alter both hemispheres, perhaps in distinct manners, making it more difficult to form a clear hypothesis about the results.

#### Design of perturbation experiments and controls

Our study was designed so that the key logic of our conclusions is based primarily on measuring perturbation effects by comparison with control conditions collected from the same session and comparison between different types of perturbations. This is a common approach used in systems neuroscience, especially in experiments using animal models and behaviors that allow the use of a task with control conditions such as comparisons of behavior before vs. after perturbation, comparison of effects on different classes of sensory stimuli or motor responses, and/or comparison of perturbations in different brain areas (e.g. ^35, 38, 107–112^). In our case, we employ all of these types of controls. Specifically, our study was designed for (A) testing the presence and validity of perturbation effects via internal controls, by comparing conditions within each experiment and testing whether they have specific relationships predicted by our hypothesis (before vs. after perturbation, contralateral vs. ipsilateral CS location, CS-directed vs. trial-initiating saccades), (B) testing the regional specificity of effects, via comparing perturbations in the two brain areas and testing whether they have the specific, distinct effects predicted by our hypothesis (icbDS vs. Pal). Below, we set out this logic in greater detail.

1. We must test whether effects on behavior are caused by the perturbation – i.e. by the injection of a substance into the brain. As an alternative, behaviors may be merely generalized side-effects caused by the experimental apparatus for injection (e.g. the weight of the apparatus on the head causing attention to be biased to a specific location in space). We ruled this out by computing all key behavioral measures as differences between behavior both before vs. after the injection. Such effects cannot be generalized effects of the apparatus because it was present throughout the entire experiment. As another alternative, behaviors may change due to generalized effects of time (e.g. behavior changing over the course of a session as the animal becomes satiated). We ruled this out in two manners. First, we used a comparison between different types of perturbations. Perturbation experiments in icbDS and Pal were carried out using the same experimental timecourse, so any main effect of time should occur equally in the two datasets. Instead, perturbing icbDS vs. Pal resulted in specific, significantly different effects on behavior that were predicted by our hypothesis. Second, as an additional test, we carried out sham injection experiments that followed exactly the same experimental timecourse as muscimol injections, so any main effect of time should occur equally in these datasets as well. Instead, there was no significant ‘before vs. after’ difference in our measure of information seeking during sham sessions, and sham sessions were significantly different from muscimol sessions. Thus, we conclude that the measured perturbation effects were indeed caused by the perturbation.
2. We must test whether effects are caused by perturbing the targeted area. As an alternative, effects may be merely non-specific effects of perturbing the brain, or the local region of the basal ganglia, rather than of perturbing the specifically targeted nuclei. To rule this out, we used two controls. First, we compared perturbation effects between contralateral vs ipsilateral space. If perturbations caused a generalized effect on brain function then it would have similar effects for saccades to either side. Instead, perturbation effects were significantly stronger in space contralateral to the perturbation target, as predicted by our hypothesis. Second, we compared perturbation effects between the two targeted areas. If effects were simply due to perturbing the rough vicinity of this area of the basal ganglia *per se*, then we would expect similar effects from perturbing icbDS vs Pal. Instead, perturbing the two targets impaired information seeking in specific, opposite manners as predicted by our hypothesis.
3. We must test whether effects are caused by perturbing information seeking behavior. As an alternative, effects may have been due to a more general effect on motivation (e.g. saccades becoming faster due to higher general motivation to seek juice). To rule this out, we used two internal controls. We computed two independent measures of general motivation – latency to saccade to the fixation point to initiate the trial and probability of completing the trial successfully. If perturbations affected information seeking due to a generalized effect on motivation, they should have a consistent effect on these measures. Instead, we found that this was not the case: icbDS perturbations slowed saccades to the Info CS despite no reduction in general motivation, while Pal perturbations speeded saccades to the Noinfo CS despite no enhancement of general motivation (in fact, this speeding of saccades occurred despite a modest reduction of general motivation, consistent with the results of previous Pal studies).

Thus, we conclude that the perturbation effects were caused by the perturbations having an effect on the targeted areas and thereby altering information seeking behavior. Future research may further interrogate these areas to discover the specific biochemical process by which our perturbations affected icbDS and Pal neurons and information seeking behavior. Our results are consistent with injections having their effect by the classic action of muscimol via agonizing GABAA receptors and hence inhibiting striatal and pallidal neurons. This is based the fact that muscimol is a highly potent and selective GABA_A_ agonist^113, 114^ and GABA_A_ receptors potently inhibit striatal and pallidal neurons ^115^. In addition, there are two lines of evidence from our study supporting this possibility. First, the observed effects on information seeking behavior are as predicted from a perturbation that has its effect via inactivating nearby neurons, including: reduction in measures of information seeking from both icbDS and Pal injections, the specific manner in which icbDS injections affected saccades to the Info CS, and the specific manner in which Pal injections affected saccades to the Noinfo CS (as illustrated by our model and its predictions in Fig 7). Second, the observed effect of muscimol injection in Pal causing a modest reduction in measures of general motivation (e.g. error rates), which is consistent with previous studies which injected muscimol into Pal and compared it to other substances ^32, 33, 37^.

